# Regulation of Toxic RNA Foci and Mutant *DMPK* Transcripts: Role of MBNL Proteins and RNA Decay Pathways

**DOI:** 10.1101/2023.09.28.559487

**Authors:** Xiaomeng Xing, Robert Markus, Tushar Ghosh, Sarah Buxton, Daniel J. Nieves, Marzena Wojciechowska, J. David Brook

## Abstract

Myotonic dystrophy type 1 (DM1) is a progressive, multisystemic disorder caused by an expansion of CTG repeats in the 3’ untranslated region of the *DMPK* gene. When transcribed the mutant RNAs accumulate in affected tissues appearing as distinct foci when visualised by *in situ* hybridisation. The RNA foci are aggregates of CUG repeat-containing RNAs that sequester RNA-binding proteins, particularly muscleblind-like (MBNL) proteins, leading to their dysfunction and causing downstream molecular and cellular defects. Here we show the double knock-out of *MBNL1* and *2* prevents RNA foci formation and nuclear retention of mutant *DMPK* mRNA in DM1 cells as well as promoting their degradation and nuclear export. Using stochastic optical reconstruction microscopy (STORM), we find the presence of both large foci and micro foci in DM1 cells. Large foci consist of multiple DMPK transcripts, while many micro foci are (CUG)n fragments. The absence of MBNL proteins not only prevents the aggregation of multiple *DMPK* transcripts into large foci, but also promotes their degradation and nuclear processing. However, although a substantial amount of MBNL1 proteins are bound to the mutant transcripts, the pools of free MBNL1 proteins are similar in DM1 nuclei to those in controls. Furthermore, we have identified several factors that are involved in the control of mutant *DMPK* mRNA turnover, including XRN2, EXOSC10, UPF1 and STAU1. Our study indicates that these factors are implicated in the RNA foci accumulation and the degradation of mutant *DMPK* mRNA. UPF1 and STAU1 may have additional roles beyond degradation, impacting the nuclear processing of mutant *DMPK* mRNA. Our study also highlights the critical role of MBNL proteins in regulating mutant *DMPK* mRNA metabolism: the absence of MBNLs in DM1 appears to expedite the processing of mutant *DMPK* mRNA mediated by these RNA decay factors.

**Significance statement:** Our investigations uncovered valuable data on the RNA foci dynamics in DM1, revealing the intricate mechanisms that underlie their formation, stability, and turnover. Our findings also contributed to delineate the complex pathways involved in the transportation and degradation of the mutant mRNA and provided insights into the critical role played by MBNL proteins in these processes. Studying the degradation mechanism of mutant *DMPK* mRNA in myotonic dystrophy may provide a foundation for comprehending the mechanisms of RNA degradation in other diseases caused by short tandem repeat (STR) mutations, such as Huntington’s disease, Fragile X syndrome, and several types of ataxia. Additionally, the use of cutting-edge STORM technology can provide a valuable tool for investigating RNA foci in other STR expansion disorders.

## Introduction

Myotonic dystrophy type 1 (DM1) is a progressive, multisystemic disorder that affects approximately 1 in 8,000 individuals worldwide [1, 2]. It is caused by an expansion of CTG repeats in the 3’ untranslated region of the *DMPK* gene, which, when transcribed results in the accumulation of mutant RNA foci in affected tissues [3]. The RNA foci are abnormal aggregates of CUG repeat-containing RNA that sequester RNA-binding proteins, particularly muscleblind-like (MBNL) proteins, leading to their functional loss and causing downstream molecular and cellular defects [4-6].

MBNL proteins are key regulators of alternative splicing, and their sequestration by RNA foci disrupts their normal function, leading to mis-splicing events that contribute to DM1 pathology [6, 7]. Mis-splicing affects the expression of genes involved in muscle function, metabolism, and development, leading to the clinical symptoms of DM1, including myotonia, muscle weakness, cardiac conduction defects, and cognitive impairment [8-10]. Recent studies have shown that restoring MBNL protein function can reverse some of the molecular and cellular defects in DM1 models, highlighting the importance of understanding the role of MBNL proteins the disease pathogenesis [11-13].

The pathogenesis of DM1 is associated with nuclear retained CUG expansion *DMPK* mRNA. However, little is known about the degradation of it, and what enzymes are responsible. Although several therapeutic approaches [12, 14-16] have appeared to eliminate nuclear foci when visualised using *in situ* hybridisation, they have not successfully reversed the entire pathogenesis of DM1 as it is not clear what happens to the mutant RNA transcripts after the dissolution/disappearance of nuclear foci. Thus, having a better understanding of the metabolism of mutant *DMPK* mRNA and the effect of the (CUG)^exp^ RNA -MBNL protein complex on mutant *DMPK* mRNA decay could have important therapeutic implications for DM.

RNA foci, pivotal contributors to DM1 pathology, exhibit a dynamic nature characterized by turnover, aggregation, and dispersion [17-20]. Although distinct, large RNA foci feature prominently in many molecular studies of DM1 [19, 21-23], few studies have explored their turn-over, particularly the identification of break-down products originating from these structures. This can be attributed to the inherent resolution constraints and lack of detection sensitivity of conventional microscopy, which preclude accurate estimations of size and distribution (detected by fluorescence or intensity) for these breakdown products. To this end, we set out to visualize and quantify the break-down products of large foci, using single-molecule localization microscopy (SMLM). SMLM methods such as stochastic optical reconstruction microscopy (STORM) can achieve resolutions in the 20- to 30-nm range by exploiting the temporal separation of individual fluorescing molecules [24, 25]. SMLM overcomes the limitations of conventional microscopy, facilitating nanoscale resolution to determine the dynamics of RNA foci.

It is well-established that most mRNAs, presumably including the wild-type *DMPK* mRNA, go through deadenylation-dependent decay in the cytoplasm [26, 27]. However, due to the presence of long expanded CUG repeats in the 3’ UTR and association with MBNL, the mutant *DMPK* mRNA has prolonged nuclear residence and is therefore initially subject to nuclear RNA surveillance [28-30]. Nuclear RNA decay is primarily conducted by exonuclease XRN2 in the 5’ → 3’ direction [31, 32] or/and by the nuclear exosome (with catalytic subunit EXOSC10/RRP6) in the 3’ → 5’ direction [33-36]. Studies by Garcia et al. (2014) suggest that the nonsense-mediated mRNA decay (NMD) pathway which principally operates in the cytoplasm, plays a conserved role in regulating CUG repeat RNA transcript levels and toxicity [37]. For example, inactivating UPF1 protein by RNAi in human DM1 patient fibroblasts causes an increase in the number of toxic RNA foci [37]. Therefore, NMD could be directly involved in the mutant *DMPK* RNA decay as the presence of expanded CUG repeats in *DMPK* mRNAs may trigger NMD due to the extended distance between the stop codon and the poly(A) signal [37-39]. Additionally, the RNA hairpin structures formed by the expanded CUG repeats in the 3’UTR of *DMPK* mRNA are very likely to be the binding sites of STAU proteins [40, 41]. Therefore, mutant *DMPK* mRNA may be subject to Staufen-mediated mRNA decay (SMD) [40, 42, 43] as well. Taken together, we hypothesised that several decay pathways might play a role in the degradation of mutant *DMPK* RNA.

In this study we have sought to examine RNA foci dynamics in DM1, to gain a better understanding of the machinery governing mutant RNA degradation and transportation, and decipher the pivotal role played by MBNL proteins in these processes. To achieve this, we have conducted the dual knockout of *MBNL1* and *MBNL2* in DM1 fibroblast cells via CRISPR/Cas9 genome engineering, resulting in the ablation of full-length MBNL1 and MBNL2 proteins. Using STORM technology, we have studied the breakdown products of (CUG)^exp^ RNA foci in DM1 cells and tissues, and in MBNL-deficient DM1 cells. Additionally, we have investigated the participation of XRN2, EXOSC10, UPF1 and STAU1 in the degradation and nuclear processing of mutant *DMPK* mRNA and the impact of MBNL proteins on these pathways.

## Results

### The double knock-out of *MBNL1 & 2* reduces nuclear foci in DM1 cells

Two MBNL-deficient DM1 cell lines (P4C1 and P4B6) were generated following CRISPR/Cas9-mediated *MBNL* knock-out in DM1 fibroblast cells, KBTeloMyoD (KB). The CRISPR-Cas9 target sites were in exon 4 (258nt) of *MBNL1* and exon 6 (154nt) of *MBNL2* (***SI Appendix,* Table S4** and **Fig. S1**). Our observations showed that MBNL1 was completely absent from P4C1 and truncated in P4B6 resulting in the loss of domains encoded by several exons, including ZnF 3-4, while MBNL2 was significantly truncated in both P4C1 and P4B6 cell lines with the loss of domain encoded by constitutive exon 6 (**Fig. 1** A) (***SI Appendix,* Fig. S2, Fig. S3, Table S5 and Fig. S4**). Thus, the endogenous expression of both *MBNL1* and *MBNL2* was effectively disrupted in both cell lines.

**Fig. 1.**
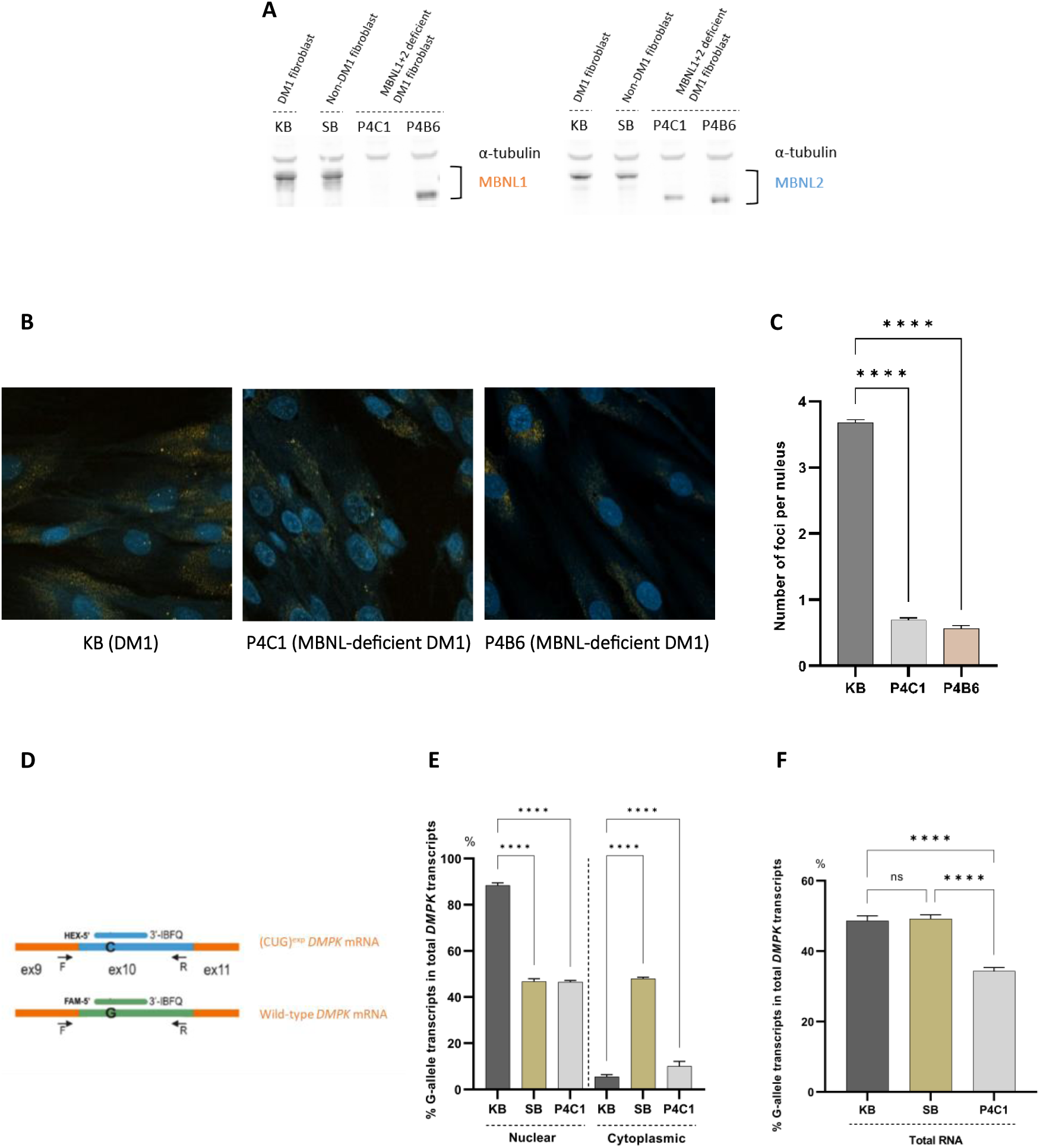
Double knock-out of *MBNL1* and *2* reduces foci formation and nuclear retention of mutant *DMPK* mRNA. (A) Total protein extracts from KB (a DM1 patient cell line), SB (a non-DM control cell line), as well as P4C1 and P4B6 (MBNL-deficient DM1 cell lines) were analysed by Western blotting to detect the expression of MBNL1 and MBNL2 proteins. α-tubulin was used as a loading control. (B) Images of KB, P4C1 and P4B6 cells following RNA-FISH using Cy3-(CAG)_10_ probe (amber) for the localisation of CUG repeats in the mutant *DMPK* mRNA and Hoechst (blue) to indicate the cell nuclei. (C) Effect of *MBNL1* and *2* double knock-out on nuclear foci. Histograms show data compiled from the RNA *in situ* hybridisation in KB (n=1351), P4C1 (n=1108) and P4B6 (n=1196) cells. n indicates the total number of cells analysed. (D) Scheme of probe design for digital PCR assay. FAM and HEX-labelled probes to bind within the rs527221 SNP site in the exon 10 of *DMPK*. They have the same nucleotide composition except for the single nucleotide difference (C or G) present in the middle of each probe. The probes had Iowa Black® FQ (IBFQ) as a quencher. (E) dPCR quantification of mutant (CUG)^exp^ *DMPK* transcripts as a percentage of total *DMPK* RNA, using allele-specific probes following total RNA extraction from KB, SB, and P4C1. (F) dPCR quantification of mutant (CUG)^exp^ *DMPK* transcripts in the nucleus and cytoplasm as a percentage of total *DMPK* RNA, using allele-specific probes following segregating RNA from the nucleus and cytoplasm of KB, SB, and P4C1. The G-allele *DMPK* transcript represents the mutant (CUG)^exp^ *DMPK* transcript in DM1. The experiment was performed in triplicate to ensure the reproducibility of the results. Statistical significance was determined by one-way or two-way ANOVA (mean ± SD, **** for P <0.0001) or indicated as not significant (NS) for P>0.05.

To determine the effect of *MBNL1* and *2* double knock-out on nuclear RNA foci, RNA fluorescence *in situ* hybridisation (RNA-FISH) was performed, using a Cy3-(CAG)_10_ labelled probe. A significant reduction in nuclear RNA foci was produced in the two MBNL-deficient DM1 cells lines compared to unmodified DM1 cells. The number of foci was decreased by ∼89.4% and ∼85.7% in P4C1 and P4B6, respectively (**Fig. 1**). Thus, our data suggests that the absence of MBNL1 and 2 is sufficient to prevent nucleation of the predominant fraction of mutant *DMPK* RNA into foci.

To measure accurately the CTG repeat lengths within the human *DMPK* gene and ensure they had not been truncated in the derived lines, we used Cas9-assisted targeting of chromosome segments (CATCH) in conjunction with Nanopore DNA sequencing. As shown in ***SI Appendix,* Fig. S6**, there was significant variation in the *DMPK* CTG expansion in KB cells as well as in the two MBNL-deficient DM1 cell lines. KB demonstrated CTG repeat lengths ranging from 6 to 2574 repeat units, with a median of 284 units for the expanded repeat lengths (>50 units). P4C1 displayed CTG repeat lengths ranging from 6 to 1553 units, with a median of 940 units for the expanded repeat lengths. P4B6 exhibited CTG repeat lengths ranging from 6 to 346 units, with a median of 291 units for the expanded repeat lengths.

These results give us confidence in the validity of using P4C1 or P4B6 cells for subsequent experiments. However, P4C1 provides a better representation of the DM1 cell line as the *DMPK* CTG repeat lengths are more comparable to KB, whereas P4B6 generally lacked the longer CTG repeat expansions (***SI Appendix,* Fig. S6**). Additionally, P4C1 completely lacked the MBNL1 protein, while a truncated version was detectable in P4B6 (**Fig. 1** A) (***SI Appendix,* Fig. S2**). Given these advantages, P4C1 was chosen for subsequent experiments.

We have previously exploited a single nucleotide polymorphism (SNP) rs527221 G>C located in exon 10 of *DMPK* to distinguish the mutant (CUG)^exp^ RNA from the wild-type *DMPK* RNA [44, 45]. In individuals affected by DM1 who also carry the heterozygous rs527221 SNP, the guanine (G) is linked with the CTG expansion allele, whereas the cytosine (C) is associated with the normal allele. Here, we employed Nanoplate Digital PCR (dPCR) to distinguish and quantify the two types of *DMPK* transcripts: FAM and HEX-labelled probes were designed to bind within the variable site and distinguish the two alleles. The probes had the same nucleotide composition except for the ‘C’ or ‘G’ present in the middle of each probe (**Fig. 1** D). SBTeloMyoD (SB) was established from a donor who did not have DM but who is heterozygous for the rs527221 SNP. As such, it provided a control for the distribution of the two types of *DMPK* transcripts.

Following the double knock-out of *MBNL1* and *2*, the proportion of mutant (G-allele) transcripts in total DMPK mRNA dropped from 48.61±1.34% to 34.38±1.02% (Fig. 1 E, KB and P4C1). This suggests that the absence of MBNL1 and 2 enhances the degradation of mutant *DMPK* transcripts in DM1 cells. Furthermore, **Fig. 1** F shows the ratio of the mutant (G-allele)-to-wild-type (C-allele) *DMPK* transcripts was skewed in the nucleus and cytoplasm. The amount of mutant DMPK transcripts in the DM1 nucleus (KB) was 88.38±1.06% of the total nuclear *DMPK* RNA, compared to 46.50±0.73% in MBNL-deficient cells (P4C1) (**Fig. 1** F, KB and P4C1). Conversely in the cytoplasmic fraction, the proportion of mutant *DMPK* transcripts was 5.43±0.90% in KB, compared to 10.06±2.10% in P4C1. These findings indicate that the absence of MBNL1 and 2 not only promotes the degradation of the mutant *DMPK* transcripts in the nucleus but may also facilitates their export to the cytoplasm.

### Mutant (CUG)^exp^ *DMPK* transcripts are present in the DM cytoplasm

To gain a better understanding of the distribution of *DMPK* transcripts and the composition of nuclear RNA foci, we applied RNA-FISH and STORM to improve their visualisation and quantification. In the first instance, we used an ATTO647n-(CAG)_8_ probe (**Fig. 2** A) to detect the CUG repeat region of expansion RNA in cell lines KB, SB and P4C1 (**Fig. 2** B-D). Data was compiled on the number of (CUG)_8_ probe signals (often referred as probe localisations in SMLM-related literature) and analysed through Bayesian cluster analysis [46] with the corresponding fluorophore coordinates and uncertainties. Signals from non-DM1 cells (SB) were utilised to identify clusters derived from background or non-(CUG)^exp^ signals. Specifically, we found that 95% of clusters identified with the (CAG)_8_ probe in non-DM1 (SB) cells contained ≤ 10 signals (***SI Appendix,* Fig. S7** *A* and *B*) (**Fig. 2** E). Additionally, the distribution of these clusters clearly distinguishes DM1 (KB) and MBNL-deficient DM1 (P4C1) nuclei (***SI Appendix,* Fig. S7** *A*) (**Fig. 2** E). Based on these findings, we categorised the clusters into three distinct groups (**Fig. 2** E): 1) Large (CUG)^exp^ foci: These clusters containing over 250 signals were observed exclusively in the DM1 (KB) nuclei. They exhibit a range of several hundred to several thousand signals and are equivalent to the nuclear foci visualised using conventional fluorescent microscopy. 2) Micro (CUG)^exp^ foci containing between 11 and 250 signals which were detected in the nucleus and cytoplasm of both DM1 and MBNL-deficient DM1 (P4C1) cells. 3) Non-(CUG)^exp^ signals consisting of clusters with ≤ 10 signals which were considered as background and excluded from the further consideration. To quantify the number of clusters in a representative cytoplasmic window, we computed the average number of clusters per 100 μm^2^ cytoplasmic area. No large (CUG)^exp^ foci were detected in the cytoplasm of KB or P4C1 cells, (***SI Appendix,* Fig. S7** *B*) (**Fig. 2** E), but intriguingly, we observed the presence of micro (CUG)^exp^ foci in the cytoplasm of both cell lines (***SI Appendix,* Fig. S7** *B*) (**Fig. 2** E).

**Fig. 2.**
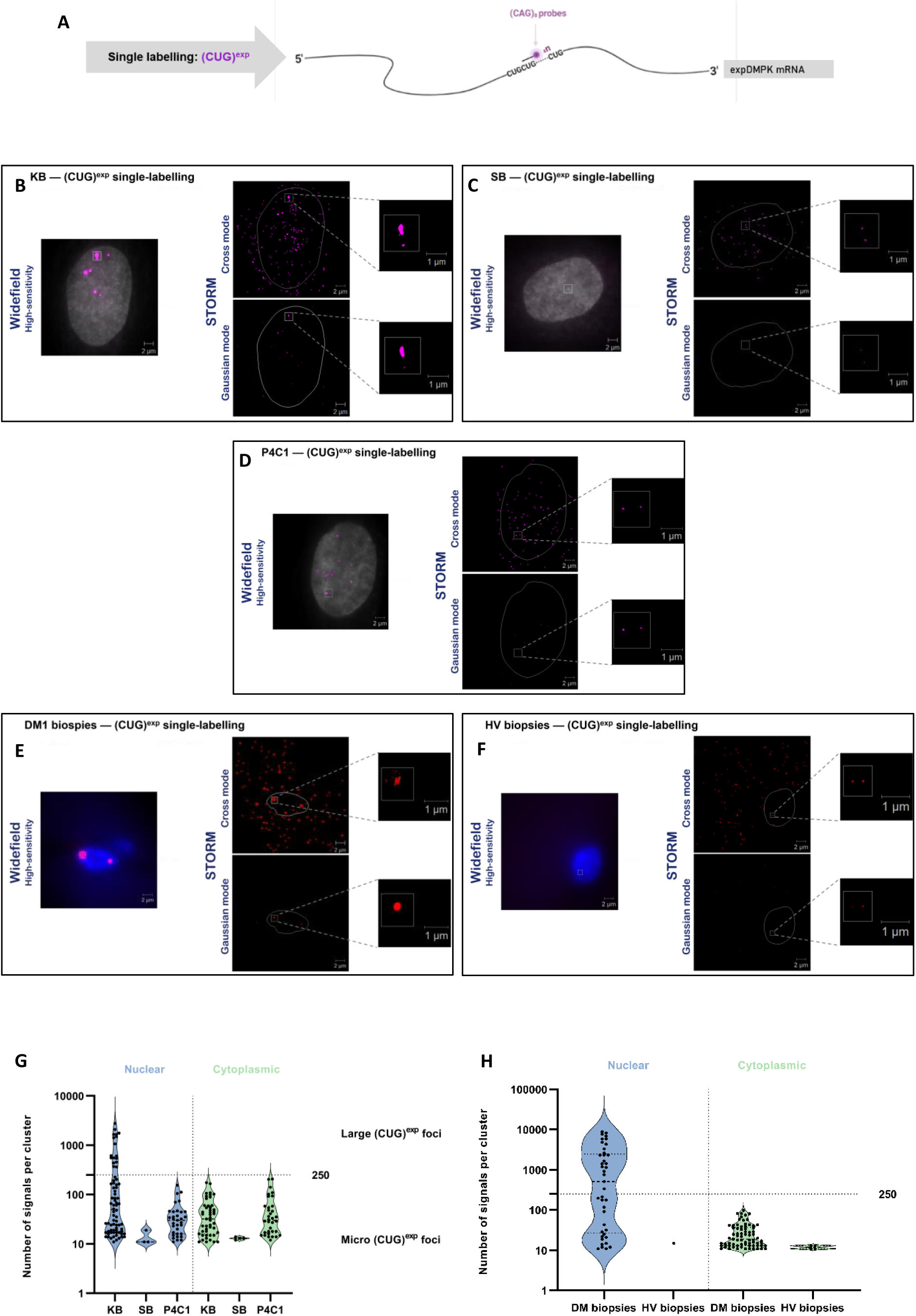
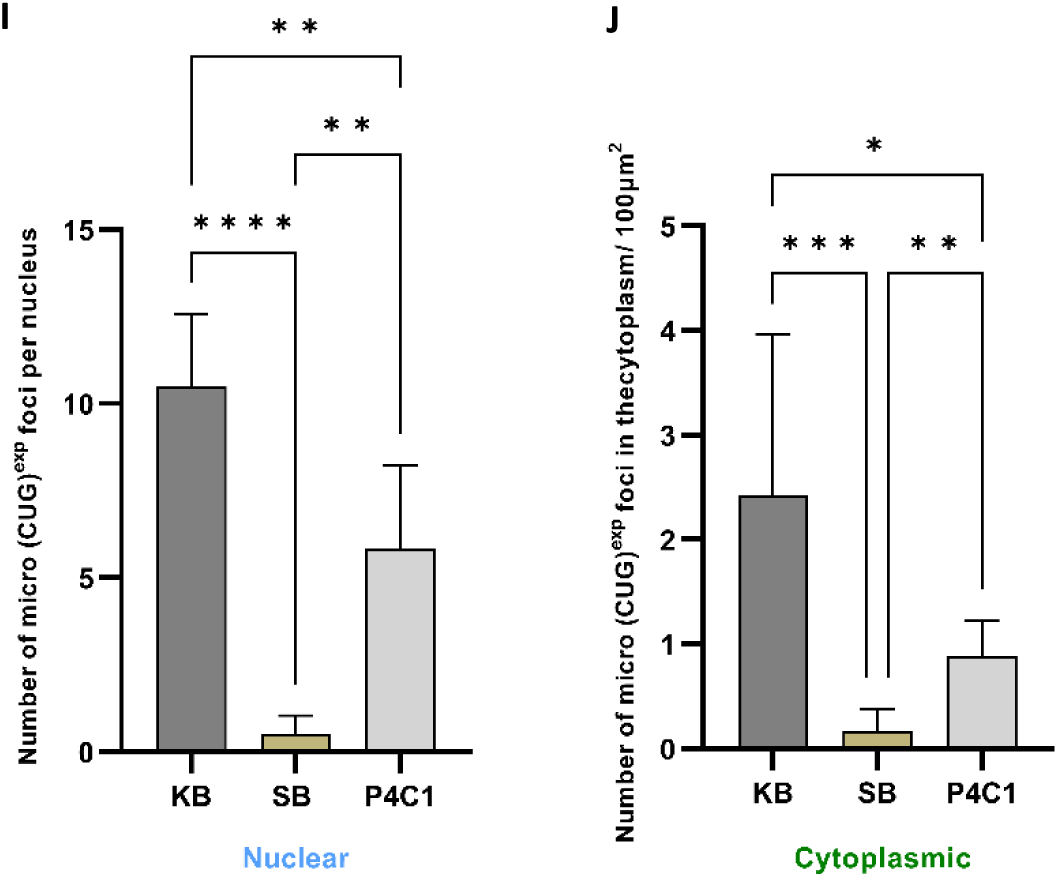
Detecting CUG repeat expansion in the *DMPK* transcript with STORM. (A) An ATTO647n-(CAG)_8_ probe was designed to detect the CUG repeat expansion in the mutant *DMPK* transcript. (B, C, D, E, F) Conventional high-sensitivity widefield images (left) and STORM images (right) of DM1 fibroblast cells (KB, n=6), non-DM fibroblast cells (SB, control, n=6), MBNL-deficient DM1 cells [P4C1, (n=6)] derived from KB, skeletal muscle tissues from DM1 patients [DM31.1 (n=5) and DM34.1 (n=5)] and skeletal muscle tissues from healthy volunteers [HV05.1 (n=3) and HV10.1 (n=3)], following RNA-FISH using the ATTO647n-(CAG)_8_ probe (magenta/red) for the localisation of CUG repeat expansion and Hoechst (grey/blue) to indicate the cell nuclei. Individual fluorophore localisation coordinates were recorded by STORM: under cross mode, each fluorophore signal (blinking) is shown as a cross; under gaussian mode, fluorophore signals are reconstructed to be shown in the way by mimicking the visual effect of human eyes. n indicates the total number of cells analysed. (G, H) Violin plots showing the distribution of clusters identified with the (CAG)_8_ probe in cells (G) and muscle biopsies (H). Each data point represents a cluster and the Y-axis indicates the number of fluorophore signals within each cluster. The violins are color-coded to distinguish between clusters detected in the nucleus (blue) and cytoplasm (green). The dashed lines within each violin indicate the median value of the cluster distribution, while the dotted lines represent the interquartile range (IQR), encompassing the middle 50% of the data. (I) Histograms show the mean number of micro (CUG)^exp^ foci per nucleus. (J) Histograms show the mean number of micro (CUG)^exp^ foci per 100 μm^2^ cytoplasmic area. Statistical significance was determined by one-way ANOVA (mean ± SD, * for P<0.05, ** for P<0.01, *** for P<0.001, **** for P<0.0001) or indicated as not significant (NS) for P>0.05.

On average, each KB nucleus contained 2.83 ± 1.17 large (CUG)^exp^ foci, which accounted for 78.96 ± 17.48% of the total (CUG)^exp^ nuclear signals (***SI Appendix,* Table S6** *A*). We also observed 10.50 ± 2.07 micro foci per nucleus and 2.43 ± 1.54 micro foci per 100 μm² cytoplasmic area in KB cells (***SI Appendix,* Table S6** *B*) (**Fig. 2** F and G). In addition to the absence of large (CUG)^exp^ foci in P4C1, the number of micro (CUG)^exp^ foci as significantly decreased in both the nuclear and cytoplasmic compartments of P4C1 compared to those of KB (**Fig. 2** F and G). Specifically, P4C1 had 5.5 ± 2.40 micro foci per nucleus and 1.57 ± 0.35 micro foci per 100 μm² cytoplasmic area (***SI Appendix,* Table S6** *B*) (**Fig. 2** F and G). Thus, the absence of MBNL proteins impacts on the formation of large (CUG)^exp^ foci and on the micro (CUG)^exp^ foci in DM1 cells. These findings suggest that the MBNL proteins strengthen the nuclear retention of mutant *DMPK* transcripts facilitating the formation of large foci, and in the absence of MBNLs, the mutant *DMPK* transcripts can only form micro foci and are processed more rapidly.

To further corroborate our findings, we performed (CUG)^exp^ single-labelling STORM experiments using an ATTO647n-(CAG)_8_ probe in skeletal muscle tissue from individuals with DM1 (DM31.1 and DM34.1) as well as healthy volunteer (HV) controls (HV05.1 and HV10.1) (**Fig. 2** H and I). The same thresholds were applied to define foci in the muscle biopsies as for cultured cells. These experiments support our observations in KB cells, we detected both large and micro (CUG)^exp^ foci in the nuclei of DM1 muscle biopsies (***SI Appendix,* Fig. S7** *C* and *D*) (***SI Appendix,* Table S6** *C* and *D*) (**Fig. 2** I). Notably, the size of some large foci, in terms of (CAG)_8_ probe signals, exceeded the sizes we previously observed in DM1 fibroblast cells (KB). This difference can likely be attributed to the generally higher *DMPK* expression levels in muscle cells compared to fibroblast cells. Moreover, we observed micro foci in the cytoplasm of DM1 muscle biopsies, suggesting their potential for migration from the nucleus to the cytoplasm.

### Large foci consist of multiple *DMPK* transcripts and many micro foci are (CUG)n fragments

To investigate the integrity of the mutant *DMPK* mRNA, we designed probes from three intervals of the *DMPK* transcript (Fig. 3 A). We used an ATTO550-(CAG)_8_ probe to bind to the CUG repeat region in the *DMPK* transcript. For the 5’ and 3’ labelling, we designed and tested a collection of 8 non-overlapping ATTO647n-labelled single copy probes for each side of the repeat. We performed two dual-labelling STORM experiments (**Fig. 3** A-G). In one, we used the 5’ probe set with the (CAG)_8_ probe in a dual-labelling experiment (**Fig. 3** A-D). In the other experiment, we performed dual-labelling with the 3’ probe set and the (CAG)_8_ probe (**Fig. 3** A, E-G). As shown in the single labelling experiment above, the (CAG)_8_ probe complementary to the *DMPK* repeat expansion does not detect the wild-type *DMPK* transcript if we set the background threshold at 10 signals. However, this is not the case for the probe sets complementary to the 5’ and 3’ to the repeat expansion. ***SI Appendix,* Fig. S8** *B-E* and **Fig. 3** H and I illustrate the distribution of clusters identified with the 5’ or 3’ single-copy probe sets. To determine an appropriate signal threshold for these clusters we utilised an additional ATTO647n-labelled scramble probe as a negative control to identify false or background signals. 95% of the clusters identified with the scramble probe contained 2-7 fluorophore signals (***SI Appendix,* Fig. S8** *A*). Thus, clusters identified with the ATTO647n-labelled 5’ or 3’ probe sets containing 7 signals or less were considered background and were excluded from further consideration. We found that the numbers of 5’ or 3’ clusters were significantly increased in KB nuclei compared to those detected in SB and P4C1 nuclei (**Fig. 3** J) indicating elevated levels of nuclear *DMPK* transcripts in DM1 cells with MBNL proteins present. The number of cytoplasmic 5’ or 3’ clusters remained similar or decreased in P4C1 compared to KB (**Fig. 3** K).

**Fig. 3.**
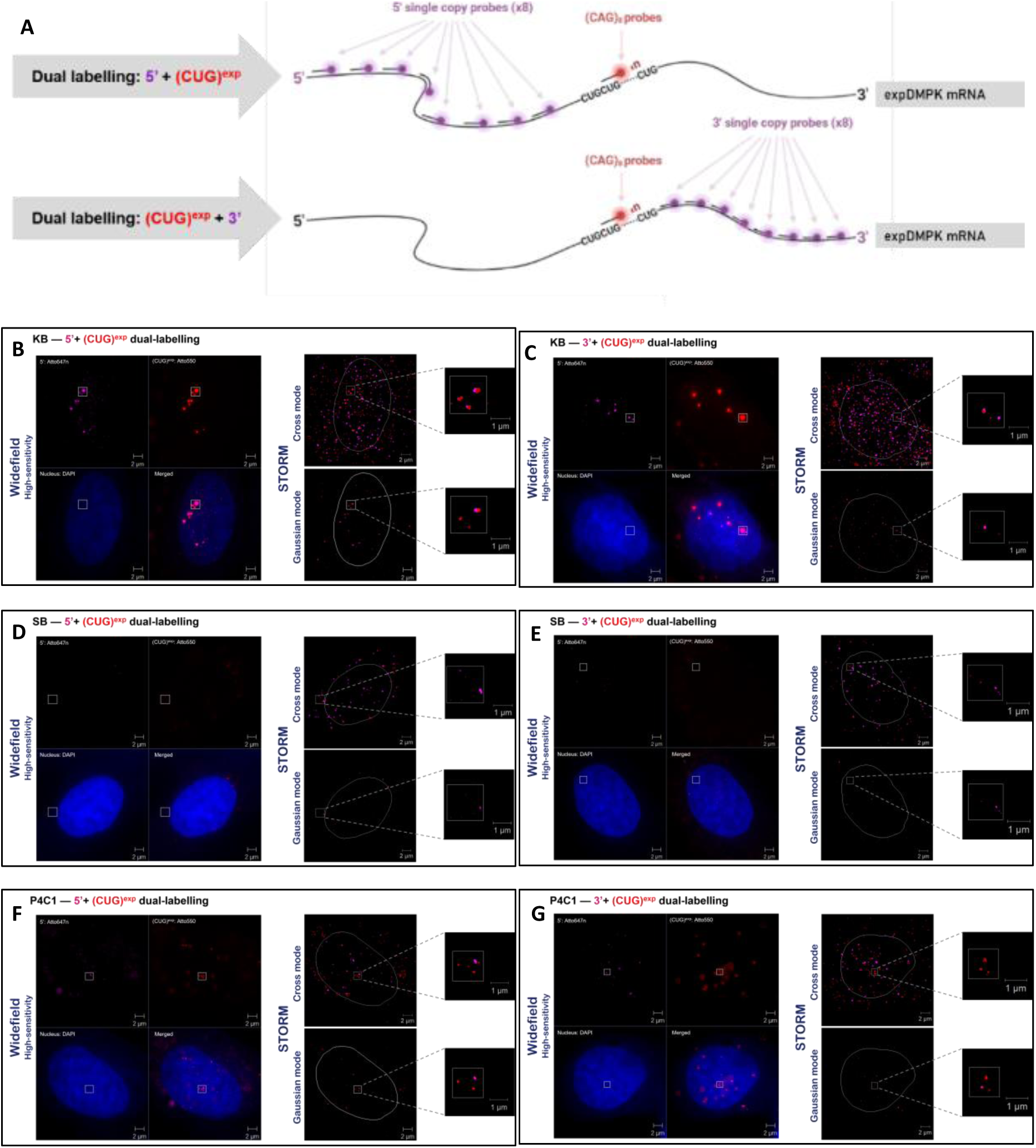
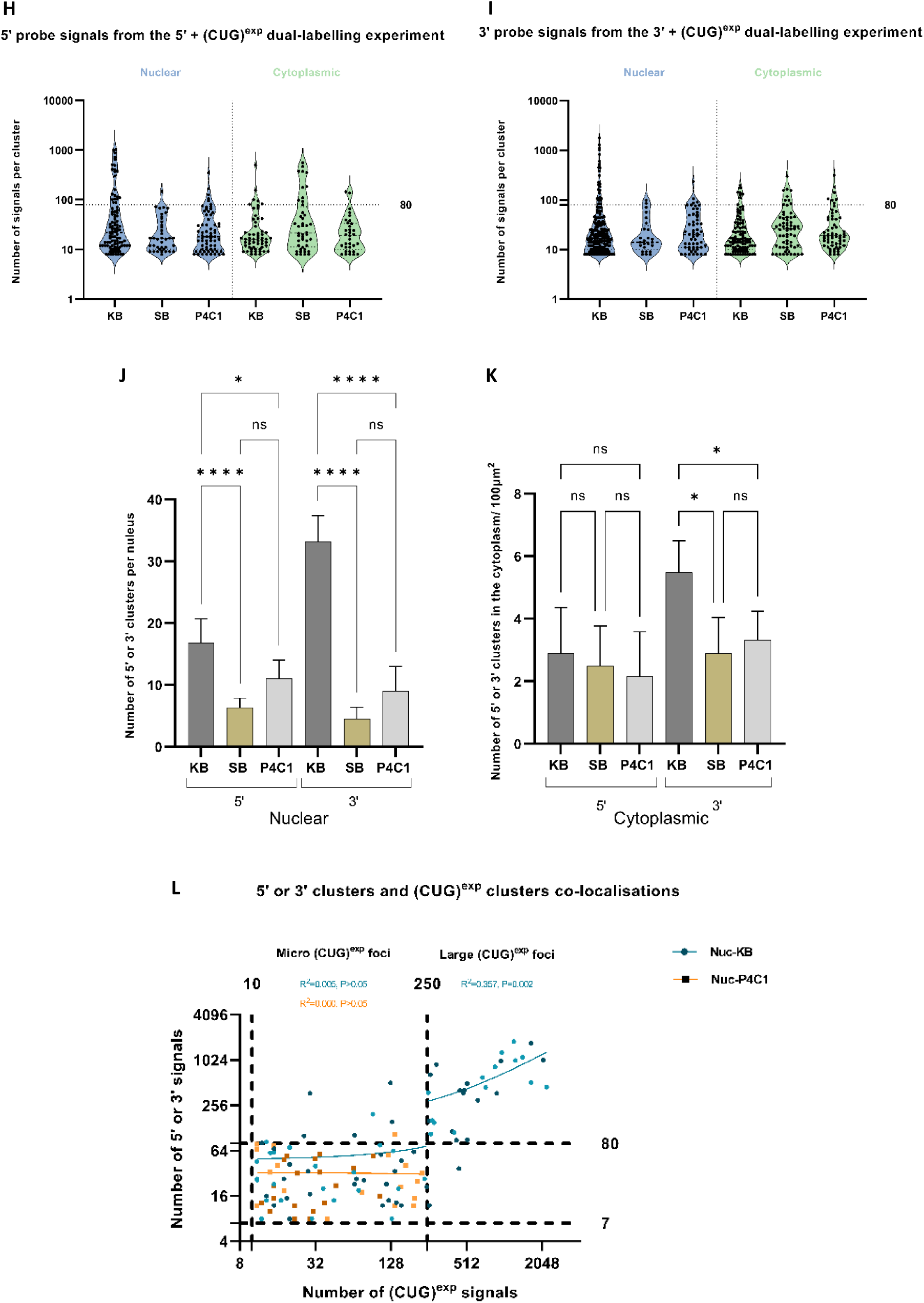
Detecting CUG repeat expansion and its 5’ or 3’ end flanking regions in the *DMPK* transcript with STORM. (A) An ATTO550-(CAG)_8_ probe was designed to bind to the CUG repeat region and 8 ATTO647n-labelled 5’ or 3’ single copy probes were designed to bind to the 5’ or 3’ flanking region of the CUG repeats in the *DMPK* transcript, respectively. (B, C, D, E, F, G) Conventional high-sensitivity widefield images (left) and STORM images (right) of DM1 (KB, n=6), Non-DM (SB, n=6) and MBNL-deficient DM1 (P4C1, n=6) cells following RNA-FISH using the ATTO647n-5’ or 3’ single copy probe set (magenta) for the localisations of 5’ or 3’ flanking regions; ATTO550-(CAG)_8_ probe (red) for the localisation of CUG repeat expansion and Hoechst (blue) to indicate the cell nuclei. Individual fluorophore localisation coordinates were recorded by STORM: under cross mode, each fluorophore signal (blinking) is shown as a cross; under gaussian mode, fluorophore signals are reconstructed to be shown in the way by mimicking the visual effect of human eyes. n indicates the total number of cells analysed. (H, I) Violin plots showing the distribution of clusters identified with the 5’ or 3’ single-copy probe sets. Each data point represents a cluster and the Y-axis indicates the number of fluorophore signals within each cluster. The violins are color-coded to distinguish between clusters detected in the nucleus (blue) and cytoplasm (green). The dashed lines within each violin indicate the median value of the cluster distribution, while the dotted lines represent the interquartile range (IQR), encompassing the middle 50% of the data. (J) Histogram shows the mean number of 5’ or 3’ clusters per nucleus. (K) Histogram shows the mean number of 5’ or 3’ clusters per 100 μm^2^ cytoplasmic area. Statistical significance was determined by two-way ANOVA (mean ± SD, * for P<0.05, **** for P<0.0001) or indicated as not significant (NS) for P>0.05. (L) Scatter plots showing (CUG)^exp^ foci and 5’ or 3’ clusters co-localisations. Each dot represents a co-localisation event in KB cells with dark blue dots indicating 5’ clusters and (CUG)^exp^ foci co-localisations and light blue dots indicating 3’ clusters and (CUG)^exp^ foci co-localisations. Similarly, each square represents co-localisation events in P4C1 cells with dark orange squares indicating 5’ clusters and (CUG)^exp^ foci co-localisations and light orange squares indicating 3’ clusters and (CUG)^exp^ foci co-localisations. The data points are fitted with a linear regression line (solid line). The X-axis indicates the number of signals in each co-localising (CUG)^exp^ focus and the Y-axis indicate the number of signals in each co-localising 5’ or 3’ cluster. The R^2^ values indicate the goodness of fit. A p-value of <0.05 indicates statistical significance.

Next, we examined the association of (CUG)^exp^ foci with the 5’ or 3’ clusters and found that a substantial proportion (86.94 ± 13.06%) of the large foci in the KB nucleus displayed co-localisation with 5’ or 3’ clusters (***SI Appendix,* Table S7** *A* and *C*), the majority of which were highly dense clusters, characterised by approximately 81 or more fluorophore signals (**Fig. 3** L). Moreover, linear regression analysis demonstrated that a significant positive correlation between the number of (CAG)_8_ probe signals and the number of 5’ or 3’ probe signals in each co-localisation event within the zone of large (CUG)^exp^ foci (**Fig. 3** L). These observations imply that the co-localising high-signal 5’ or 3’ clusters represent an exceptional accumulation of mutant *DMPK* transcripts in large (CUG)^exp^ foci, such that larger foci, as indicated by the number of (CAG)_8_ signals, contain a greater number of *DMPK* mRNA molecules. Conversely, only a small proportion of micro (CUG)^exp^ foci co-localised with 5’ or 3’ clusters (***SI Appendix,* Table S7** *A-D*). In the KB nucleus the figure was 33.24 ± 18.72% and in the cytoplasm, 14.33 ± 10.82%. In the P4C1 nucleus 25.20 ± 20.70% of micro foci co-localised with 5’ or 3’ clusters and 18.94 ± 12.22% in the cytoplasm. Notably, the majority of these co-localising 5’ or 3’ clusters exhibited counts of 80 signals or fewer (**Fig. 3** L). Additionally, the results of the linear regression analysis demonstrated that there is no correlation between (CAG)_8_ probe signals and 5’ or 3’ probe signals in the micro foci that possess 5’ or 3’ flanking sequences, (**Fig. 3** L). These findings suggest that while the majority of micro foci may originate from standalone (CUG)n repeat fragments lacking flanking regions, among the remaining micro foci, their size in terms of (CAG)_8_ probe signals does not correspond to an increased number of 5’ or 3’ probe signals, indicating diverse forms of micro foci, possibly containing DMPK mRNAs at different degradation states. Moreover, ∼ 18.9% of cytoplasmic micro foci are flanked by 5’ or 3’ sequences in P4C1 compared to ∼ 14.3% in KB, indicating a greater proportion of relatively intact expansion transcripts present in the cytoplasm when MBNLs are absence.

### The pools of free MBNL1 proteins are similar in DM1 nuclei to those in controls

Next, we examined the distribution of MBNL1 proteins in cultured DM1 fibroblast cells and muscle tissues from DM1 patients, through Immunofluorescence (IF) + RNA-FISH and STORM experiments. The captured images provided clear evidence of the co-localisation of MBNL1 clusters and RNA foci (**Fig. 4** A-D). ***SI Appendix,* Fig. S9** *B-E* and **Fig. 4** E illustrate the distribution of fluorophore signals in MBNL1 clusters in various cell and tissue types, including DM1 fibroblast cells (KB), non-DM fibroblast cells (SB), skeletal muscle tissues from DM1 patients, and healthy volunteers. To account for non-specific binding, we conducted an additional control experiment by performing IF + RNA-FISH in cells where the primary antibody was replaced with dH_2_O during the incubation step. Images and data obtained from these cells were used as negative controls, in which, the majority (95%) of clusters contained 8 or fewer fluorophore signals (***SI Appendix,* Fig. S9** *A*). Therefore, clusters with 9 or more signals were considered as MBNL1 clusters, while clusters with 8 or fewer signals were attributed to non-specific binding and were excluded from further consideration. In KB cells and DM1 muscle biopsies more highly dense MBNL1 clusters, characterised by approximately 125 or more fluorophore signals, were observed in the nucleus compared to their respective controls (**Fig. 4** E), and 74.80 ± 25.80% of these clusters were co-localised with (CUG)^exp^ foci in DM1 nuclei (***SI Appendix,* Table S8** *A* and *C*). Linear regression analysis (**Fig. 4** F) demonstrated a significant positive correlation between the number of (CAG)_8_ probe signals and the number of MBNL1 signals within the zone of large (CUG)^exp^ foci. This is consistent with previous studies suggesting that MBNL1 binding is proportional to the length of CUG repeats [47]. However, no such correlation was observed within the zone of micro (CUG)^exp^ foci. ***SI Appendix,* Fig S9** *G* shows that the total number of MBNL1 signals per nucleus is substantially greater in KB cells and DM1 biopsies compared to their respective controls, indicating an increased accumulation of MBNL1 in these nuclei. However, the total number of MBNL1 signals per 100 μm^2^ cytoplasmic area is similar in KB cells and DM1 biopsies to their respective controls (***SI Appendix,* Fig S9** *H*). Furthermore, only 6.93±6.41% and 11.41±9.02% of MBNL1 signals co-localised with micro (CUG)^exp^ foci in the cytoplasm of KB and DM1 biopsies, respectively (***SI Appendix,* Table S8** *B* and *D*), suggesting the nuclear accumulation of MBNL1 does not affect the distribution of MBNL1 in the cytoplasm. In KB nuclei, 39.52±11.50% of MBNL1 signals were associated with the (CUG)^exp^ foci, while in the nuclei of DM1 muscle biopsies, this figure is 73.17±19.52% (**Fig. 4** G). However, the number of free MBNL1 signals [those not co-localising with (CUG)^exp^ foci] in the nuclei of KB and DM1 biopsies was similar or higher to the number in their respective controls (**Fig. 4** H). This suggests that despite a substantial portion of MBNL1 proteins being sequestered by the (CUG)^exp^ *DMPK* transcripts, there appears to be a sufficient pool of free MBNL1 proteins in DM1 nuclei.

**Fig. 4.**
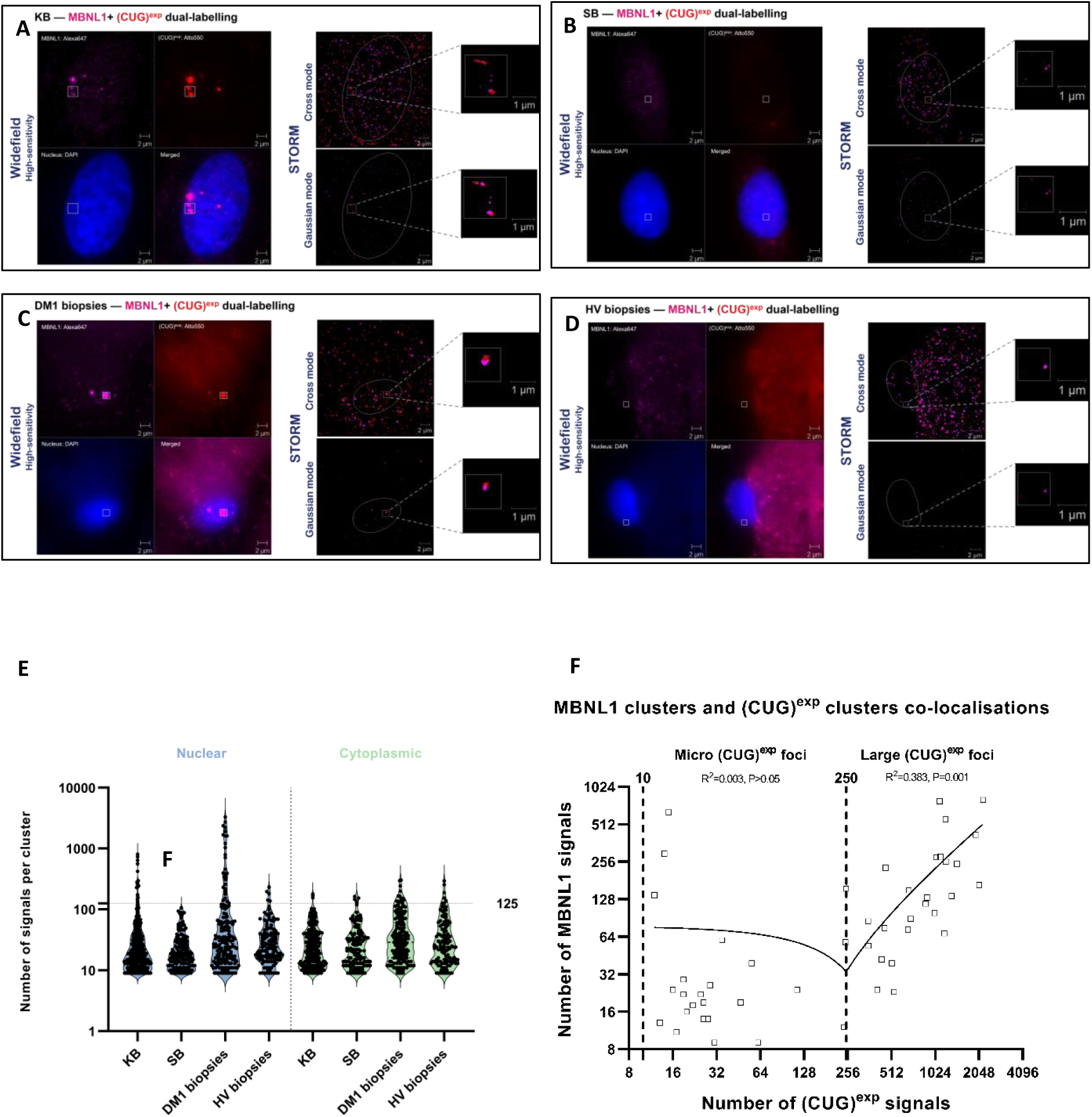
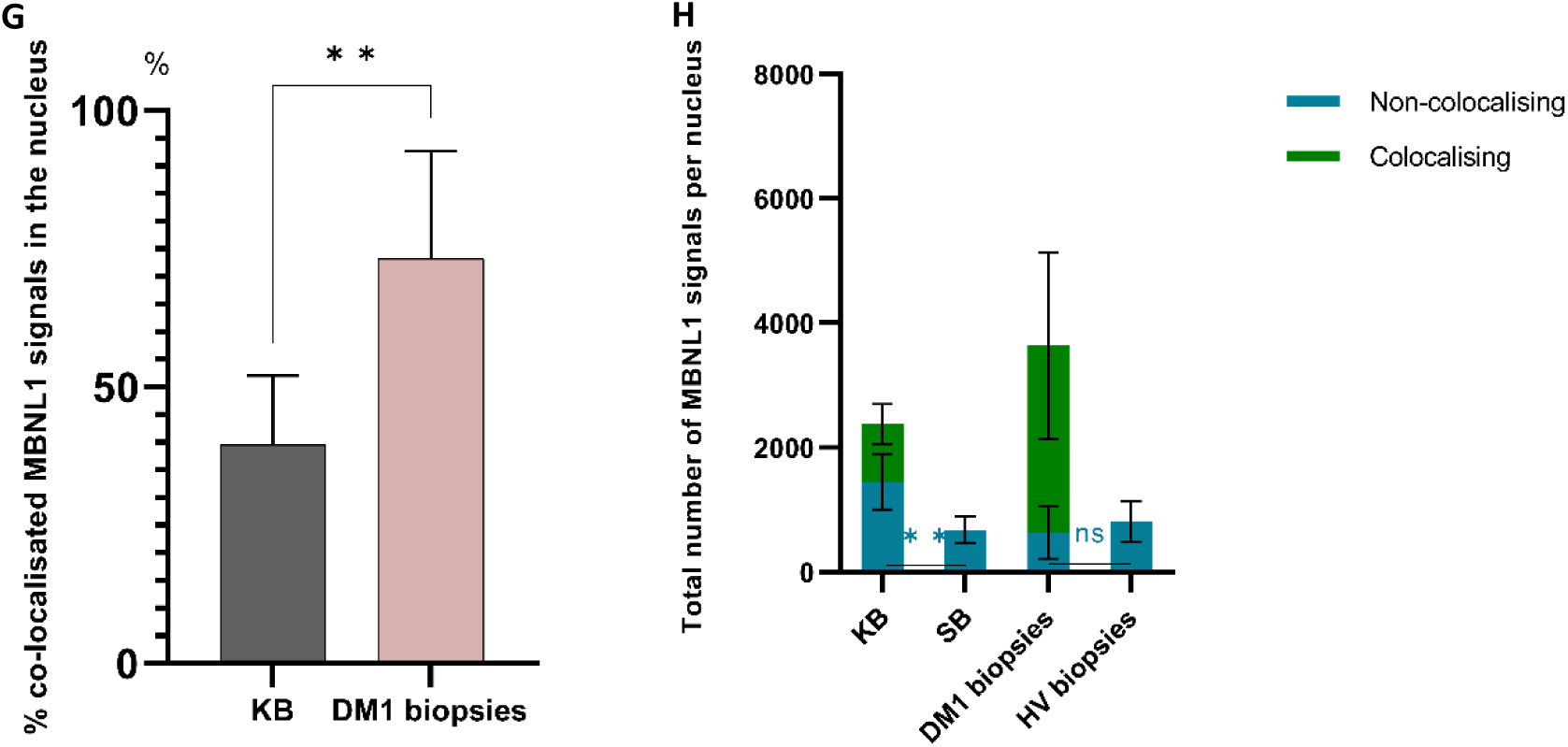
STORM—MBNL1 and (CUG)^exp^ dual-labelling. (A, B, C, D) Conventional high-sensitivity widefield images (left) and STORM images (right) of DM1 fibroblast cells (KB, n=7), non-DM fibroblast cells (SB, control, n=6), skeletal muscle tissues from DM1 patients [DM31.1 (n=5) and DM34.1 (n=5)] and skeletal muscle tissues from healthy volunteers [HV05.1 (n=3) and HV10.1 (n=3)], following RNA-FISH using a mouse anti-MBNL1 primary antibody and an anti-mouse Alexa647-labelled secondary antibody (magenta) for the localisations of MBNL1 proteins; ATTO550-(CAG)_8_ probe (red) for the localisation of CUG repeat expansion and Hoechst (blue) to indicate the cell nuclei. n indicates the total number of cells analysed. Individual fluorophore localisation coordinates were recorded by STORM: under cross mode, each fluorophore signal (blinking) is shown as a cross; under gaussian mode, fluorophore signals are reconstructed to be shown in the way by mimicking the visual effect of human eyes. (E) Violin plots showing the distribution of clusters identified with a mouse anti-MBNL1 primary antibody and an anti-mouse Alexa647 secondary antibody. Each data point represents a cluster and the Y-axis indicates the number of fluorophore signals within each cluster. The violins are color-coded to distinguish between clusters detected in the nucleus (blue) and cytoplasm (green). The dashed lines within each violin indicate the median value of the cluster distribution, while the dotted lines represent the interquartile range (IQR), encompassing the middle 50% of the data. (F) Scatter plot showing (CUG)^exp^ foci and MBNL1 clusters co-localisations in KB cells. The data points are fitted with a linear regression line (solid line). The R^2^ values indicate the goodness of fit. A p-value of <0.05 indicates statistical significance. (G) Histogram showing the percentage of MBNL1 signals co-localised with (CUG)^exp^ signals in the nucleus, relative to the total number of MBNL signals. (H) Histogram showing the average total number of MBNL1 signals detected per nucleus, signals that co-localise with (CUG)^exp^ foci are represented in green, while those not co-localising with (CUG)^exp^ foci are shown in blue. Statistical significance was determined by two-tailed student t-test or one-way ANOVA (mean ± SD, ** for P<0.01) or indicated as not significant (NS) for P>0.05.

### Multiple RNA decay pathways are involved in the degradation of mutant (CUG)^exp^ *DMPK* mRNAs

To investigate the role of XRN2, EXOSC10, UPF1, and STAU1 in the degradation of mutant (CUG)^exp^ *DMPK* transcripts, we knocked down these RNA decay factors in DM1 (KB) and MBNL-deficient DM1 (P4C1) cells using lentiviral shRNAs. Western blot analyses were conducted to assess the knock-down efficiency. An equivalent reduction of protein levels was observed in both KB and P4C1 cells, with XRN2 reduced by ∼ 76.8%, EXOSC10 by ∼ 50.0%, UPF1 by ∼ 56.5% and STAU1 by ∼ 69.9% (***SI Appendix,* Fig. S11** *A-L*).

The effect of decay factor knock-down on RNA foci was examined using RNA-FISH (**Fig. 5** A and B) and a significant increase in foci area was observed in KB following knock-down of *XRN2* (∼ 10.9%), *EXOSC10* (∼ 6.5%), *UPF1* (∼ 36.3%) and *STAU1* (∼ 13.1%) compared to scramble controls. A relatively greater increase in foci area was observed in P4C1 following knock-down of *XRN2* (∼ 180.0%), *EXOSC10* (∼ 33.3%), *UPF1* (∼ 182.4%) and *STAU1* (∼92.3%) (**Fig. 5** C-F), which indicates their role in reducing RNA foci formation and in the absence of MBNLs, they act more effectively.

**Fig. 5.**
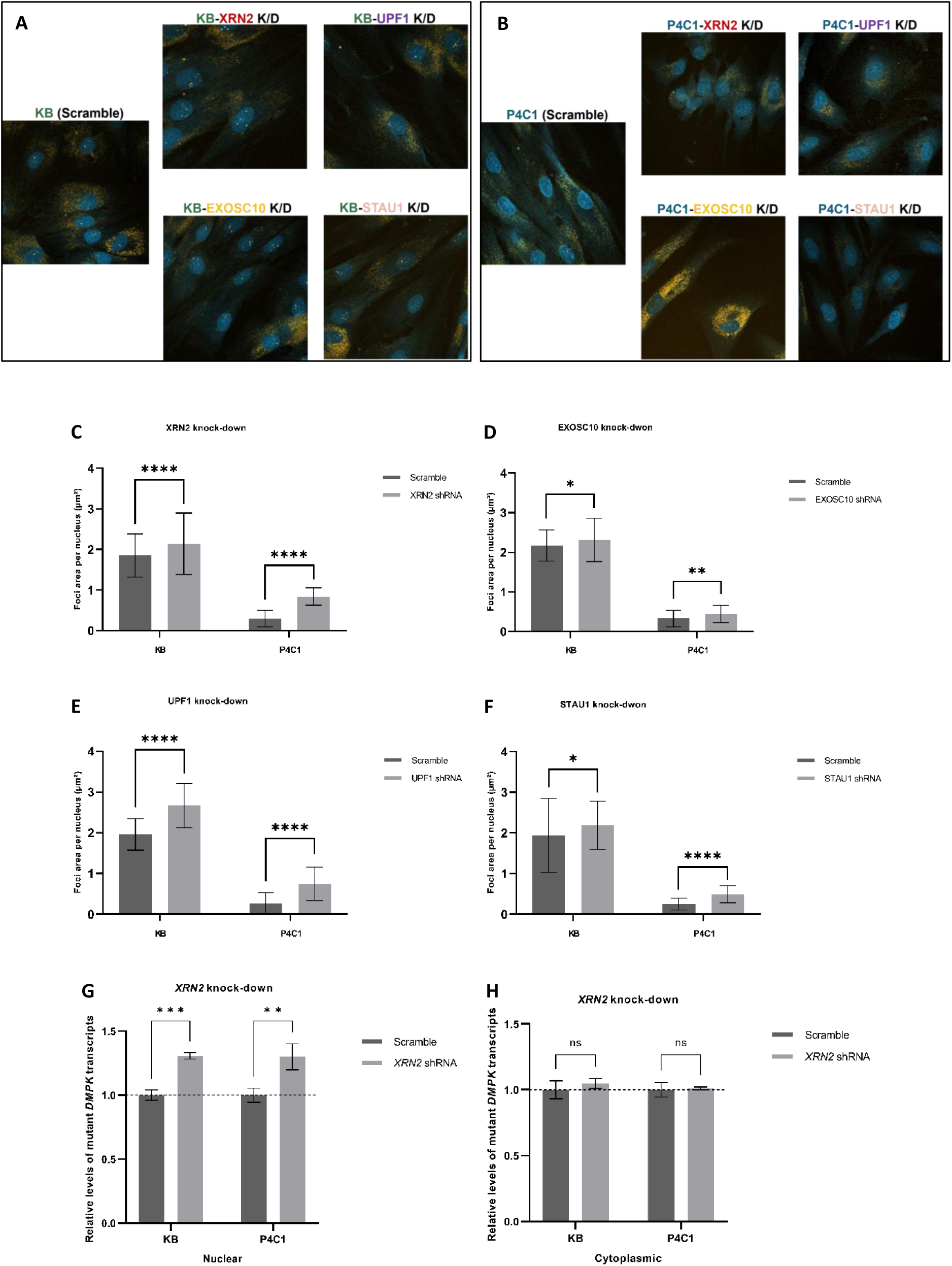
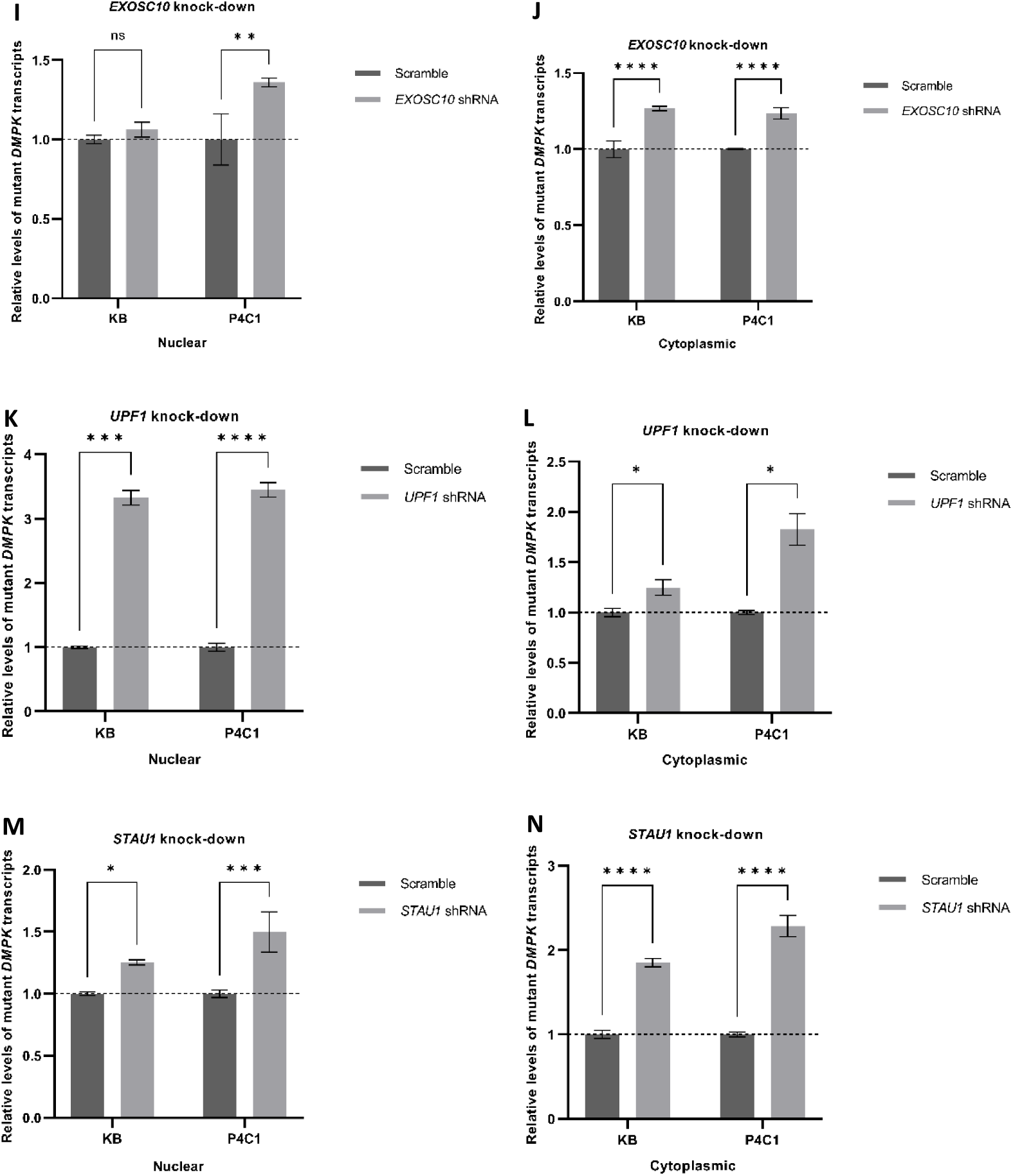
Knock-downs of RNA decay factors in unmodified DM1 cells and MBNL-deficient DM1 cells. **Effect of the knock-down (KD) of decay factors on RNA foci.** (A, B) Images of KB and P4C1 cells following the lentiviral shRNA knock-down of *XRN2*, *EXOSC10*, *UPF1* and *STAU1* or transfection with corresponding scramble shRNA lentiviral particles and RNA-FISH using Cy3-(CAG)_10_ probe (amber) for the localisation of CUG repeats in the mutant *DMPK* mRNA and Hoechst (blue) to indicate the cell nuclei. (C, D, E, F) Quantification of RNA foci area per nucleus with data compiled from the RNA-FISH in above experimental cells. **Effect of the knock-down of RNA decay factors on the mutant *DMPK* transcripts.** (G, H, I, J, K, L, M, N) Quantification of the relative level of nuclear and cytoplasmic (CUG)^exp^ *DMPK* transcripts in KB and P4C1 following the lentiviral shRNA knock-down of *XRN2* (G, H), *EXOSC10* (I, J), *UPF1* (K,L) and *STAU1* (M, N) or transfection with corresponding scramble shRNA lentiviral particles by measuring the abundance of 5’ end to the CUG repeats, normalised against the level of MALAT transcripts for the nuclear fraction and GAPDH transcripts for the cytoplasmic fraction. Scramble shRNA data established baselines at 1.0 for each condition. The experiment was performed in triplicate to ensure the reproducibility of the results. The total number of cells examined by RNA-FISH was listed in ***SI Appendix,* Table S8**. Statistical significance was determined by Two-way ANOVA (mean ± SD, * for P<0.05, ** for P<0.01, *** for P<0.001, **** for P<0.0001) or indicated as not significant (NS) for P>0.05.

Next, we investigated the impact of RNA decay factor knock-downs on mutant *DMPK* transcripts in the nuclear and cytoplasmic fractions of KB and P4C1. *XRN2* knock-down produced a significant increase in mutant *DMPK* transcripts in the nuclei of KB (∼ 30.9%) and P4C1 (∼ 30.6%) but had no significant effect on their cytoplasmic abundance in either cell lines (**Fig. 5** G and H). Knock-down of *EXOSC10* had negligible impact on mutant *DMPK* transcripts in KB nuclei, yet it induced a significant increase in P4C1 nuclei (∼ 38.2%) (**Fig. 5** I). Notably, in cytoplasmic fractions *EXOSC10* knock-down produced an increase mutant *DMPK* transcript abundance in both KB (∼ 20.3%) and P4C1 (∼ 21.8%) cells (**Fig. 5** J).

The individual knock-downs of *UPF1* and *STAU1* produced similar effects. In each case we observed a significant increase in the abundance of nuclear mutant *DMPK* transcripts in both KB (∼ 232.8% for *UPF1* and ∼ 25.3% for *STAU1*) and P4C1 (∼ 246.3% for *UPF1* and ∼ 49.9% for *STAU1*) (**Fig. 5** K and M). Moreover, their knock-downs also resulted in an increase in the abundance of cytoplasmic mutant *DMPK* transcripts. In the KB cytoplasm we observed increases of ∼ 21.3% for *UPF1* and ∼ 86.1% for *STAU1* whereas the impact of knock-downs was significantly greater in P4C1 at ∼ 81.4% for *UPF1* and ∼ 129.0% for *STAU1* (**Fig. 5** L and N).

These findings suggest that XRN2, EXOSC10, UPF1 and STAU1 are implicated in the nuclear accumulation of RNA foci, as well as in the degradation of mutant *DMPK* mRNAs in the nucleus and/or cytoplasm.

Next, we examined the potential therapeutic benefit of enhancing the UPF1 pathway as its involvement in mRNA subcellular localisation [48, 49] and mRNA surveillance [50, 51] is well documented. We over-expressed *UPF1* in KB and P4C1 through lentiviral gene transfer. Western blot analysis showed the UPF1 level in infected KB and P4C1 cells to be twice as high as in the control cells (***SI Appendix,* Fig. S12** *A-C*). *UPF1* over-expression produced a significant reduction in the foci area in both KB (∼ 29.9%) and P4C1 (∼ 82.8%) (**Fig. 6** A and B) when visualised by RNA-FISH. Next, we investigated the impact of *UPF1* over-expression on mutant *DMPK* transcripts. **Fig. 6** C and D shows the over-expression of *UPF1* resulted in a significant reduction in mutant *DMPK* transcript abundance relative to wild-type *DMPK* transcripts in both nuclear (∼ 23.4%) and cytoplasmic (∼ 25.1%) fractions in KB. The effect of *UPF1* over-expression was even greater in P4C1 cells with a reduction in the relative abundance of the mutant transcript of ∼ 60.6% and ∼ 40.1% in the nucleus and cytoplasm, respectively, suggesting that *UPF1* over-expression has a more pronounced effect on the degradation of mutant *DMPK* transcripts in MBNL-deficient DM1 cells compared to unmodified DM1 cells.

**Fig. 6.**
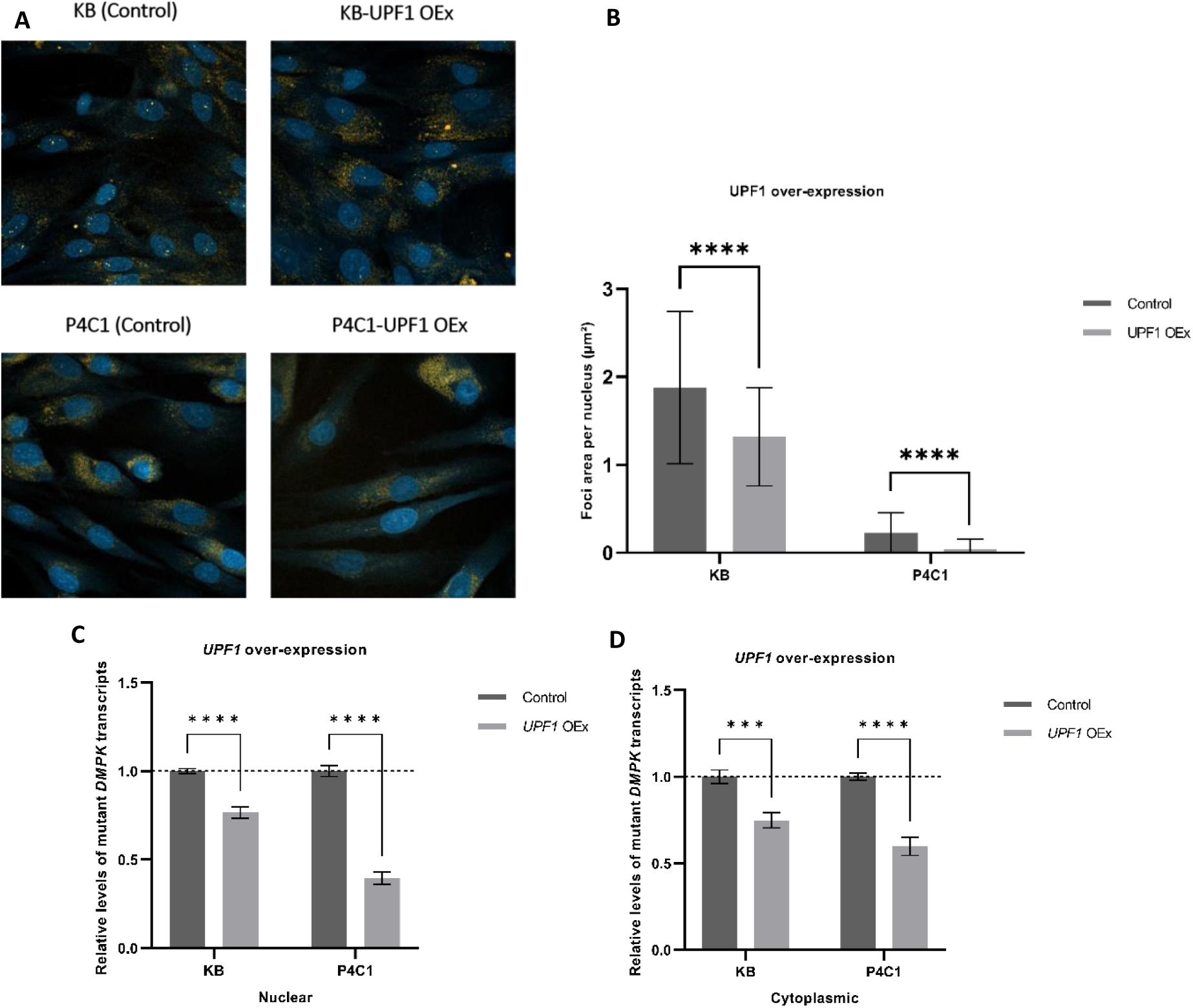
U*P*F1 over-expression in unmodified DM1 cells and MBNL-deficient DM1 cells. **Effect of *UPF1* over-expression (OEx) on RNA foci** (A) Images of KB and P4C1 cells following the lentiviral transfection with *UPF1* cDNA or empty control and RNA-FISH using Cy3-(CAG)_10_ probe (amber) for the localisation of CUG repeats in the mutant *DMPK* mRNA and Hoechst (blue) to indicate the cell nuclei. (B) Quantification of RNA foci area per nucleus with data compiled from the RNA-FISH in above experimental cells. **Effect of the *UPF1* over-expression (OEx) on the mutant *DMPK* transcripts.** Quantification of the relative level of nuclear (C) and cytoplasmic (D) (CUG)^exp^ *DMPK* transcripts in KB and P4C1 by measuring the abundance of 5’ end to the CUG repeats, normalised against the level of MALAT transcripts for the nuclear fraction and GAPDH transcripts for the cytoplasmic fraction. The experiment was performed in triplicate to ensure the reproducibility of the results. Scramble shRNA data established baselines at 1.0 for each condition. The total number of cells examined by RNA-FISH was listed in ***SI Appendix,* Table S8**. Statistical significance was determined by Two-way ANOVA (mean ± SD, *** for P<0.001, **** for P<0.0001).

## Discussion

In this study two MBNL-deficient cell lines (P4C1 and P4B6) were generated that allowed us to examine the effect of MBNL loss on nuclear foci and mutant expansion RNA in DM1 cells. MBNL1 was either completely absent or truncated with the loss of domains encoded by several exons, including ZnF 3-4, while MBNL2 was truncated with the loss of domains encoded by constitutive exon 6 in both P4C1 and P4B6 cell lines. Thus, the endogenous expression of both *MBNL1* and *MBNL2* was effectively disrupted in these cell lines.

We found that the knock-out of MBNL1 and 2 produced a significant reduction in the number of nuclear foci. Furthermore, using an allele specific polymorphism to distinguish the wild-type and mutant *DMPK* transcripts in a digital PCR assay, we found a substantial reduction in the proportion of mutant transcripts in the nucleus and a significant increase in that in the cytoplasm following the knock-out of *MBNL1* and *2*. This suggests that the absence of MBNL1 and 2 not only facilitates the nuclear degradation of mutant *DMPK* transcripts, but also promotes their nuclear export.

Using STORM technology, we gained unprecedented insights into (CUG)^exp^ RNA foci in DM1 cells and the impact of MBNL-deficiency in DM1 cells. We used probe sets complementary to the 5’ and 3’ ends of the *DMPK* transcript to investigate the integrity of the mutant transcript. This showed that large (CUG)^exp^ foci are typically composed of multiple *DMPK* transcripts, while micro foci are primarily (CUG)n fragments. Large (CUG)^exp^ foci are only found in the nucleus, while micro foci are present in both nuclear and cytoplasmic compartments. Moreover, our study builds upon and expands the previous notion that the depletion of MBNL1 leads to elimination of RNA foci [20, 52]. We demonstrated that the absence of MBNLs modulates the breakdown of mutant *DMPK* mRNAs. In DM1, mutant *DMPK* mRNAs aggregate into large, compact (CUG)^exp^ RNA foci that can be easily detected under conventional fluorescent microscope. However, in the absence of MBNLs, mutant *DMPK* mRNAs can only form micro (CUG)^exp^ foci. Thus, the lack of MBNL promotes the degradation of mutant *DMPK* mRNAs and it may assist transportation to the cytoplasm. Our data provide a challenge to the paradigm that mutant expansion RNAs sequester MBNL proteins, rather MBNL proteins promote the nuclear retention and restrict the degradation of the mutant *DMPK* transcripts.

Additionally, the co-localisation analysis provided evidence that most of the cytoplasmic micro (CUG)^exp^ foci lack 5’ or 3’ flanking sequences, which indicates that many mutant *DMPK* transcripts exist as fragments in the cytoplasm which could provide a source for RAN translation in DM1 [53, 54].

Our findings are consistent with previous studies, demonstrating that the presence of CUG repeat expansion in the *DMPK* transcript leads to the sequestration of a significant amount of MBNL1 proteins, resulting in an overall increase in MBNL1 protein levels within the DM1 nucleus. However, we found that despite the concentration of MBNL1 proteins, there is an adequate amount of them remaining unbound, which diverges from the prevailing perspective of MBNL loss-of-function in DM1.

In KB cells, we observed that only 39.52±11.50% of MBNL1 proteins were sequestered by the (CUG)^exp^ foci. Interestingly, a positive correlation was found between the number of fluorophore signals in large (CUG)^exp^ foci and the amount of sequestered MBNL proteins in KB cells. However, the scenario was notably different in the cells of DM1 muscle biopsies. In DM1 biopsies, a much larger proportion (73.17±19.52%) of MBNL1 was sequestered by the (CUG)^exp^ foci. Surprisingly, we did not observe a significant correlation between the number of CAG probe signals and the number of MBNL1 signals within the zone of large (CUG)^exp^ foci in the DM1 biopsies (***SI Appendix,* Fig. S9** *J*). One plausible explanation for this discrepancy could be that in DM1 biopsies, MBNL1 sequestration might have reached a saturation point due to the limited availability of unbound MBNL1 proteins. Alternatively, the binding sites may be compacted in dense structures in the muscle tissues, limiting accessibility for MBNL binding. These data suggests that the dynamics of MBNL1 sequestration and its association with (CUG)^exp^ foci differ between cultured DM1 fibroblast cells and DM1 muscle biopsies, possibly reflecting the varying disease contexts and molecular interactions at play in these cellular environments.

The pathogenesis of DM1 is associated with nuclear retained CUG expansion *DMPK* mRNA. However, little is known about the degradation of it, and what enzymes are responsible. Although several therapeutic approaches [12, 14-16] have appeared to eliminate nuclear foci when visualised using *in situ* hybridisation, they have not successfully reversed the entire pathogenesis of DM1 as it is not clear what happens to the mutant RNA transcripts after the dissolution/disappearance of nuclear foci. There is no evidence showing that the foci are the sites of mutant RNA decay. Additionally, our investigation into the Poly(A) and 5’ cap status of mutant *DMPK* mRNA revealed that a significant portion of the mutant *DMPK* mRNAs were deadenylated or decapped (***SI Appendix,* Fig. S10**), indicating that their degradation may be initiated but impeded to a large extent. To gain a better understanding of the control of mutant DMPK mRNA turnover, we employed shRNA interference on 4 decay factors: XRN2, EXOSC10, STAU1 and UPF1 in conjunction with probe digital PCR to quantify the mutant *DMPK* transcripts based on the SNP situated in exon 10, 5’ to the repeat expansion region.

XRN2 is a processive 5’ → 3’ exonuclease and primarily located in the nucleus [55, 56]. Its knock-down had a significant effect on the abundance of nuclear mutant *DMPK* transcripts in KB and P4C1. In contrast, EXOSC10 is a distributive 3’ → 5’ exonuclease and human EXOSC10 is present not only in the nucleus but also in the cytoplasm [57]. Knock-down of *EXOSC10* had no significant effect on the abundance of nuclear mutant *DMPK* transcripts in KB, but resulted in a significant increase in the abundance of nuclear mutant *DMPK* transcript in P4C1 and in that of cytoplasmic mutant *DMPK* transcripts in both KB and P4C1 cells. These results suggest that both XRN2 and EXOSC10 are involved in the degradation of mutant *DMPK* transcripts. In the KB nucleus, the 3’ → 5’ degradation activity of EXOSC10 appears to be impeded. As a result, degradation by EXOSC10 probably does not proceed to the 5’ end of the repeat expansion region. However, in P4C1 where MBNLs are absent, nuclear degradation of the mutant RNAs is more effective. The absence of an equivalent 3’ SNP restricts our capacity to measure accurately the abundance of the 3’ end of mutant *DMPK* transcripts.

Overall, although both enzymes are involved in the decay of mutant *DMPK* transcripts, the functional differences between XRN2 and EXOSC10 suggest that their stalling by MBNL proteins may lead to distinct effects on the decay of the transcripts.

Nonsense-mediated decay (NMD) and Staufen-mediated decay (SMD) are two essential pathways for mRNA quality control that prevent the translation of defective transcripts [40, 51, 58-60]. The efficiency and specificity of these pathways rely on key factors, including UPF1 for NMD and STAU1 for SMD [42, 43, 61, 62]. UPF1 mainly operates on the 3’UTRs of mRNAs that are directed for NMD in the cytoplasm, but a recent study offered evidence that UPF1 also facilitates the release of mRNAs from transcription sites and their export from the nucleus to the cytoplasm [48]. In addition to its role in Staufen-mediated decay (SMD), STAU1 has also been reported to promote the nuclear export of specific mRNAs, including mutant (CUG)^exp^ *DMPK* mRNA [41]. Our results demonstrated that knocking down *UPF1* or *STAU1* in DM1 cells enhances RNA foci accumulation and inhibits the cytoplasmic degradation of mutant *DMPK* transcripts. These effects are more pronounced in MBNL-deficient DM1 cells, suggesting that both NMD and SMD pathways are essential for the degradation of mutant *DMPK* mRNA, while the sequestration of MBNLs impedes this degradation process and in the absence of MBNLs, UPF1 and STAU1 act more effectively.

In our study, we observed that inhibition of *UPF1* had a comparable effects on the nuclear mutant *DMPK* mRNA processing in both KB and P4C1, but over-expression of *UPF1* seemed to have a more profound impact on that in P4C1 than in KB. Furthermore, our results showed that knock-down of *STAU1* led to a significant increase in nuclear mutant *DMPK* transcript levels, which was even more pronounced in MBNL-deficient DM1 cells. Unlike UPF1, STAU1 is more likely to interact directly with the CUG repeats within the mutant *DMPK* mRNA and promote its nuclear export through the assistance of NXF (nuclear export factor) family proteins [40, 41, 63]. Therefore MBNL sequestration may inhibit this event to some extent. Additionally, the observation that over-expression of *UPF1* reduces RNA foci and mutant *DMPK* transcripts in DM1 cells provides support for the idea that UPF1 may be a promising target for developing therapies for DM1. However, more research would be needed to fully comprehend the underlying mechanisms of expanded *DMPK* mRNA processing by UPF1 and STAU1, and to determine the optimal level of UPF1 expression and phosphorylation for therapeutic purposes.

Taken together, our study indicates that XRN2, EXOSC10, UPF1 and STAU1 are implicated in the RNA foci accumulation and the degradation of mutant *DMPK* mRNAs. Moreover, UPF1 and STAU1 may have additional roles beyond degradation, impacting the nuclear processing of these mRNAs. Our study also highlights the critical role of MBNL proteins in regulating mutant *DMPK* mRNA metabolism. The absence of MBNLs in DM1 appears to destabilise the mutant transcripts, expediting their degradation or nuclear processing. Thus, targeting these pathways and enzymes could be a potential strategy for developing therapies for DM1. For example, drugs that can promote the activity of these enzymes could be used to facilitate the degradation of mutant *DMPK* mRNA and alleviate its toxic effects. Recent studies showed that small molecules and decoys that impact on MBNL binding [64-66] appear to facilitate the endogenous degradation of repeat expansion mRNAs. Furthermore, studying the degradation mechanisms of mutant *DMPK* mRNA in myotonic dystrophy may also provide a foundation for understanding the mechanisms of RNA degradation in other diseases caused by short tandem repeat (STR) mutations, such as Huntington’s disease, Fragile X syndrome, and several types of ataxia. The mechanisms that regulate RNA stability and degradation are complex and involve multiple pathways and RNA-binding proteins. As we gain a better understanding of the molecular mechanisms involved in RNA processing and degradation, we may be able to develop new therapies and interventions for a wide range of diseases caused by RNA dysregulation.

### Methods and Materials Cell culture

Fibroblast cells were grown in Dulbecco’s modified Eagle’s medium (DMEM) with penicillin and streptomycin and 10% fetal calf serum (FCS) (Sigma-Aldrich). Cell lines used in this study include human fibroblasts from a non-DM (SBTeloMyoD) person, a DM1 (KBTeloMyoD - 400 CTG repeats) patient.

### Sample use

The human biological samples were obtained in accordance with ethical guidelines, and their utilization for research purposes adhered to the stipulations of informed consent. DM31.1, DM34.1, HV05.1 and HV 10.1 are human vastus lateralis skeletal muscle biopsies. DM31.1 was sourced from a male DM1 patient aged 48 with an average modal lymphocyte CTG repeat size of 451. DM34.1 was sourced from a female DM1 patient aged 40 with an average modal lymphocyte CTG repeat size of 602. HV05.1 was sourced from a healthy female volunteer aged 66. HV10.1 was sourced from a healthy female volunteer aged 19.

### RNA-FISH for cells

Cells were grown in 96 well-plates or in 6-well plates with a cover glass in each well for 72 hours before fixation. Media was removed from cells and they were washed thrice with DPBS. The fixation step was performed by incubating cells in 4% (v/v) paraformaldehyde (PFA) for 30 minutes at room temperature. Cells were washed again thrice with PBS and then permeabilised in 80% (v/v) ethanol for 3 hours at room temperature. After another three washes with PBS, cells were incubated in the pre-hybridisation solution for 15 minutes at room temperature. The pre-hybridisation solution contains: 40% (v/v) formamide, 10% (v/v) 20x SSC and 50% (v/v) DEPC water. The prehybridisation solution was then replaced with the hybridisation solution. Each 10ml of hybridisation solution contained: 40% (v/v) formamide, 10% (v/v) 20x SSC, 50% (v/v) DEPC water, 10 µl of 200mM vanadyl ribonucleoside complex (VRC, RNase inhibitor), 100 µl of 10 mg/ml ssDNA (sperm salmon DNA), 20 µl of 20 mg/ml BSA and 500ng of Cy3-labelled (CAG)_10_ probes (imaging with Celldiscoverer 7) or 50ng of ATTO647n-labelled or ATTO550-labelled probes (for STORM experiments, probe details in ***SI Appendix,* Table S3** and **Fig. S13**. Cells were incubated overnight at 37℃ in the incubator. The next day, cells were washed thrice with PBS + 5 mM MgCl_2_, stained for 40 minutes at room temperature with Hoechst 33342 (fluorescent stain for DNA) and finally left in PBS + 5 mM MgCl_2_. Cells were then ready for imaging using the ZEISS Celldiscoverer 7 microscope or the Zeiss Elyra Super Resolution Microscope.

### RNA-FISH for human muscle samples

Snap-frozen human skeletal muscle samples were used. 7-µm cryostat sections were thawed onto a PLL (Poly-l-lysine)-coated coverslip. Slides were fixed in 2% (v/v) PFA in PBS for 30 min at 4°C. After two brief washes in PBS at room temperature, slides were permeabilised in 2% acetone in PBS for 5 min at 4°C, followed by two brief washes in PBS. Pre-hybridisation [40% (v/v) formamide, 10% (v/v) 20x SSC and 50% (v/v) DEPC water] was conducted at room temperature before hybridisation at 42°C [40% (v/v) formamide, 10% (v/v) 20x SSC, 50% (v/v) DEPC water, 200 µM VRC, RNase inhibitor, 100 µl of 100 µg/ml ssDNA, 40 µg/ml BSA and] for 2 hours. A Post-hybridisation wash [40% (v/v) formamide, 10% (v/v) 20x SSC and 50% (v/v) DEPC water] was performed at 45°C for 30 min followed by 2× SSC washes at room temperatures for 5 × 5 min. Slides were incubated in PBS with 5 mM MgCl_2_ for 15 min at room temperature. After two brief washes in PBS with 5 mM MgCl_2_, the stained sections were then ready for imaging using the Zeiss Elyra Super Resolution Microscope.

### IF + RNA-FISH

Samples were fixed and permeabilized with a 50:50 mixture of ice-cold methanol and acetone for 30 minutes at room temperature. Subsequently, to block nonspecific binding, samples were treated with a solution of 5% (w/v) BSA and 5% (v/v) sheep serum in PBS for 1 hour at room temperature. Next, the samples were incubated with an anti-MBNL1 mouse monoclonal antibody at a dilution of 1:1000, and the incubation was carried out overnight at 4°C. Afterward, the samples were stained with the secondary antibody, AlexaFluor647 goat anti-mouse (diluted at 1:500), for 30 minutes. Following the immunostaining, the samples were processed for RNA-FISH using appropriate RNA-FISH protocols. Finally, the samples were imaged using the Zeiss Elyra Super Resolution Microscope.

### Fluorescence Microscopy

#### Celldiscoverer 7

Fixed cells were visualized using a Zeiss Celldiscoverer 7 Microscope (Carl Zeiss, Germany) equipped with a ZEISS Plan-Apochromat 20x/0.95 Autocorr objective. Images were captured and analysed using the ZEN (blue version) software (Carl Zeiss, Germany) at room temperature. DAPI staining was excited with a 405-nm laser, while Cy3 samples were imaged using a 561-nm excitation. For image analysis, the Image Analysis tool in the ZEN software was employed. Threshold settings were carefully adjusted and applied to ensure clear distinction between RNA foci and nuclei against the background. To enhance the precision of threshold setting, SB (non-DM) cells were employed as a negative control. Additionally, object size and circularity parameters were used to exclude noise and artifacts from the analysis. The same parameter criteria were consistently applied to all samples within each 96-well plate. However, minor adjustments to the parameters might have been made when analysing samples from different plates to account for potential variations in experimental conditions.

### Super Resolution Microscopy

#### Image recording

Fixed cells or tissue adhered to 18 mm coverslips (Zeiss) were subjected to a single rinse with distilled water. To ensure controlled mounting, we used an imaging spacer (Grace Bio-Labs SecureSeal™) on a slide. We then added 9 µl of STORM buffer (Abbelight) to the central cavity of the imaging spacer. The imaging process utilised a Zeiss Elyra PS1 Super-Resolution Microscope (Carl Zeiss, Germany), equipped with a Zeiss alpha Plan-Apochromat 100x/1.46 oil immersion objective. Image acquisition was facilitated through the ZEN software (black version) from Carl Zeiss, Germany. STORM imaging was conducted using the laser widefield mode. Prior to each STORM recording, a field of view snapshot was acquired, employing noise reduction through the averaging of 16 images. The actual STORM recording encompassed the acquisition of 10,000 to 20,000 frames, each exposed for 25 ms. Sequential utilisation of 642-nm and 561-nm lasers enabled the detection of probes or secondary antibodies labelled with ATTO/Alexa647n and ATTO550, respectively. Laser power was judiciously modulated based on target characteristics, with a 100% high laser power for densely populated target molecules and 25% laser power for sparsely distributed ones. To facilitate channel separation without the need for filter cube adjustments, a dedicated filter set (Laser blocking filter LBF -561/642) was employed. This strategy ensured precise optical alignment of the two channels. Any observed x-y mismatch resulted solely from drifting, which was rectified during post-processing. The microscope’s focus remained locked in a predefined position throughout the imaging procedure.

#### Image processing

The raw image files, saved in the CZI format by Zeiss, underwent initial processing within the dedicated PALM module of the Zeiss Black 2012 software suite. To identify localizations with precision, the Gaussian fitting module was employed. Parameters for point spread function (PSF) detection were configured; the size was set to 9 pixels, and the peak intensity-to-noise ratio was established at 6. As a result, each localization was characterised by its x and y coordinates, along with additional characteristics such as photon count, localization precision, and the half-width of the PSF. To ensure a refined dataset and minimize potential false detections, a stringent filtration process was applied based on these characteristics.

#### Image analysis

Image analysis was performed using the Bayesian Cluster Analysis Package (generated by Prof. Dylan Owen and his team [46]). For characterisation of cytoplasmic/nuclear clusters, we manually segmented the cells (based on DAPI to identify nuclei and faint background fluorescence signal to identify cell body).

### Western blotting

Cells were washed with PBS and then lysed with RIPA lysis buffer (25 mM Tris-HCl pH = 7.5, 150 mM NaCl, 1% (v/v) NP-40, 1% (w/v) sodium deoxycholate, 0.1% (w/v) SDS) with protease and phosphatase inhibitors (Thermo Scientific, 78442). The lysate was homogenized by passing through a 22-gauge syringe 10 times and then incubated on ice for 30 min with vortexing every 5 min. Cell debris was removed by centrifugation at 500× g for 10 min at 4°C and lysates were mixed with 4× NuPAGE™ lithium dodecyl sulfate (LDS) sample buffer (Invitrogen, NP0007) with 10× NuPAGE™ sample reducing agent (Invitrogen, NP0004) and heated at 70°C for 10 min. Samples were loaded onto a NuPAGE™ 10% Bis-Tris polyacrylamide gel (Invitrogen, NP0301) and transferred to a PVDF membrane using iBlot 2 dry blotting system (Invitrogen, IB21001). Membranes were blocked in 5% (w/v) skim milk in tris-buffered saline with 0.1% (v/v) Tween-20 (TBST) for 1 h at room temperature. Membranes were then incubated with primary antibodies (***SI Appendix***, **Table S1**) that were diluted 1:1,000 in 1% (w/v) skim milk in TBST at 4°C overnight. After three TBST washes, membranes were incubated with HRP-conjugated secondary antibodies (***SI Appendix***, **Table S1**) at a 1:10,000 dilution in 1% (w/v) skim milk in TBST for 1 h at room temperature. Membranes were washed three times with TBST and chemiluminescence signals were detected using with the Clarity™ western ECL Substrate (BIO-RAD). The protein bands were then visualised using a LAS-3000 Imaging System. Band intensities were quantified using FIJI after subtracting background signal.

### Genome editing by CRISPR/Cas 9

For our CRISPR projects, we focused on two target genes, *MBNL1* and *MBNL2*. To design the guide RNAs (gRNAs), we utilised ChopChop version 2, an online tool known for predicting potential gRNAs based on target sequence and off-target analysis. Plasmids containing the Cas9 protein and the gRNAs were prepared for the experiments. Prior to transfection, we harvested two million cells and formed a cell pellet. Next, 100 µl of nucleofector mix (comprising 82 µl of nucleofector solution and 18 µl of supplement 1 solution) was added to the cell pellet, and the cells were carefully resuspended. The cell and nucleofector mix were then combined with the CRISPR/Cas9 vector mix, and the entire mixture was transferred into a cuvette. The cuvette was placed into the Nucleofector Amaxa Electroporator, and electroporation was conducted using the program: U23 NHDF Human Adult High Efficiency. Following electroporation, the cells were transferred to a flask with appropriate media to facilitate their growth and maintenance.

Transfected cells were subjected to genomic DNA extraction followed by PCR amplification and Sanger sequencing of PCR products to examine the efficiency of genome editing at expected target sites in the *MBNL1* and *MBNL2* gene. Fragments harbouring the target site for each gRNA were PCR amplified using the primers MBNL1-F’: 5’-GTGTGCCCTTCTCATTGTATCA- 3’ and MBNL1-R’: 5’-GAAAGTATTTGCACTTTTCCCG- 3’ for MBNL1; MBNL2-F’: 5’-TTGCCGCTGTTTAAATT- 3’ and MBNL2-R’: 5’- CAAAAGCCCTAACTGCTGTTC- 3’ for MBNL2.

### Lentiviral shRNA/expression vector transduction

Cells were seeded on a 6-well plate (0.3 million per well) at 37 ℃ in a humidified 5% CO_2_ incubator overnight. On the transduction day, the virus was thawed and mixed gently prior to use. Old media were removed from the plate and 1 ml of fresh medium was added to each well. Appropriate amount of virus (***SI Appendix,*** Table S2) and 10 µg of Polybrene were placed in each well to achieve the desired MOI. After that, the plate was swirled gently to mix and incubated at 37 ℃ in a humidified 5% CO_2_ incubator overnight. Virus-containing media were remove and replaced with fresh media after 12 hours of incubation. Cells were harvested 5 days post transduction for western blot, RNA extraction or RNA-FISH.

### Measuring the *DMPK* transcript levels

Total RNA from fibroblast cells was extracted using the Purelink^TM^ RNA mini Kit (12183018A), according to the manufacturer’s protocols. Nuclear and Cytoplasmic RNA fractions from fibroblast cells were extracted using the Norgen Cytoplasmic & Nuclear RNA Purification Kit (21000), according to the manufacturer’s protocols. RNA was converted to cDNA using Verso cDNA Synthesis Kit with random hexamers (Thermo Scientific™, AB1453). *DMPK* cDNA copies were quantified using Nanoplate Digital PCR (dPCR) (QIAGEN). Primers and probes used in the digital PCR experiments were configured and purchased on the QIAGEN GeneGlobe website using the dPCR LNA® probe PCR Assays or the QuantiNova® LNA® Probe PCR Assays. To estimate the *DMPK* cDNA abundance, data were normalized to *MALAT1* for the nuclear fraction and *GAPDH* for the cytoplasmic fraction.

## Appendix/Supplementary data

Data represented in SI Appendix.

## Acknowledgments

This project was supported by the Myotonic Dystrophy Foundation, Alan Webster Endowment, Myotonic Dystrophy Support Group and University of Nottingham. We would like to thank T. Self and S. Bagia for their invaluable technical assistance. We would like to thank D. Owen at the University of Brimingham for his expert contributions to data analysis. We would like to thank S. Saam for providing the muscle biopsies.

## Author Contributions

XX, MW, JDB conceived and designed the experiments. XX, RM, TG, SB conducted the experiments. XX, RM, NJV, JDB performed data analysis and interpretation. XX, RM, JDB wrote the paper. All authors actively participated in result discussions and contributed to the manuscript.

