## Supplementary material for "Regulation of Toxic RNA Foci and Mutant *DMPK* Transcripts: Role of MBNL Proteins and RNA Decay Pathways": SI Appendix

**Extended methods**

**RNA-seq**

Total RNA was extracted from the cultured cells using the Purelink RNA Mini Kit (Life Technologies) with DNase I treatment. Total RNA integrity was confirmed by Agilent 2100 Bioanalyzer RNA nano chip. RNA-seq libraries were constructed using the NEBNext Ultra II Directional RNA library prep kit for Illumina, using ribosomal RNA depletion followed by strand-specific RNA-seq preparation. Samples were amplified with PCR for 9-11 cycles and sequenced using the Illumina NextSeq 500 v2 with 75 nucleotide paired end reads. The sequencing generated millions of short reads representing fragments of the transcriptome. Raw sequencing reads obtained from the sequencer underwent quality trimming to remove adapters and low-quality bases. Next, the trimmed reads were mapped to the reference genome using the HISAT2 alignment tool. Subsequently, read counts were quantified for each annotated feature, including genes or exons, using the featureCounts program. Principal component analysis (PCA) was applied to assess the global patterns and identify potential sample outliers. To identify differentially expressed (DE) genes, the DESeq package was employed, which implements rigorous statistical methods. Moreover, to examine isoform abundance and detect differential exon usage, de novo transcriptome assembly was conducted using StringTie.

**Nanopore DNA sequencing**

For targeted nanopore DNA sequencing with the enrichment of regions of interest (ROI) using CRISPR/Cas9, genomic DNA from fibroblast cells was extracted using the QIAGEN DNeasy Blood & Tissue kit according to the manufacturer's protocol. The DNA concentration and quality were assessed using a NanoDrop spectrophotometer and agarose gel electrophoresis. Guide RNAs (gRNAs) were designed using ChopChop version 2. Appropriate gRNAs were then purchased from IDT and resuspended in nuclease-free water at a concentration of 100 µM. The gRNA/Cas9 ribonucleoprotein (RNP) complex was prepared by mixing the gRNA and Cas9 protein at a ratio of 1.5:1 and incubating at 37°C for 10 minutes. The purified genomic DNA was then mixed with the gRNA/Cas9 RNP complex and incubated at 37°C for 1 hour. This allowed the RNP complex to bind and cleave the Region of Interest (ROI) in the genomic DNA. The DNA was purified using a PCR cleanup kit to remove any residual gRNA/Cas9 RNP complex, unbound gRNAs, and other impurities that may interfere with the downstream sequencing. The sequencing adapters were ligated to the Cas9 cut sides of the ROI using the Ligation Sequencing Kit (SQK-LSK109, Oxford Nanopore Technologies) according to the manufacturer's protocol. The ligation reaction was incubated at 20°C for 10 minutes, followed by 65°C for 10 minutes. Excess unligated adapters and other short DNA fragments were removed using the Ligation Sequencing Kit (SQK-LSK109, Oxford Nanopore Technologies) according to the manufacturer's protocol. The purified DNA was quantified using a Qubit fluorometer. The 1D2 sequencing kit (SQK-LSK308, Oxford Nanopore Technologies) was used for construction of sequencing libraries with 500 ng input DNA. The sequencing library was prepared according to the manufacturer's protocol and sequenced on a MinION sequencing device (Oxford Nanopore Technologies) using R9.5 flow-cells for 24–48 hours. The reads were base-called using the MinKNOW software and the resulting fastq files were analysed using a variety of bioinformatics tools, such as Minimap2, Samtools, and GATK, to identify and extract the relevant information. The Nanopore sequencing service and data analysis were provided by colleagues from the University of Nottingham Next Generation Sequencing Facility, who performed quality control and helped with data interpretation.

Table S1 List of primary and secondary antibodies used in this paper.

| Antibody | Source | Catalog | Working concentration |
| --- | --- | --- | --- |
| Mouse monoclonal anti-MBNL1 | Gift from Dr. Ian Holt) | MB1a | 1:1000 for WB; 1:500 for IF |
| Mouse monoclonal Anti-MBNL2 | Gift from Dr. Ian Holt) | MB2a | 1:1000 for WB; 1:500 for IF |
| Mouse monoclonal anti-Lamin-B (A-11) | Santa Cruz  Biotechnology | sc-377000 | 1:1000 for WB |
| Mouse monoclonal anti-α-tubulin (TU-02) | Santa Cruz  Biotechnology | sc-8035 | 1:1000 for WB |
| Anti-mouse IgGκ BP-HRP | Santa Cruz  Biotechnology | sc-516102 | 1:5000 for WB |
| Mouse monoclonal anti-XRN2 (H-3) | Santa Cruz  Biotechnology | sc-365258 | 1:1000 for WB |
| Mouse monoclonal anti-EXOSC10 (B-8) | Santa Cruz  Biotechnology | sc-374595 | 1:1000 for WB |
| Rabbit monoclonal  anti-STAU1 | abcam | ab137100 | 1:1000 for WB |
| Anti-rabbit IgG HRP-linked Antibody | Cell Signalling | 7074 | 1:5000 for WB |
| Rabbit monoclonal anti-RENT1/hUPF1 | abcam | ab133564 | 1:1000 for WB |
| Mouse monoclonal anti-GAPDH | Santa Cruz  Biotechnology | sc-47724 | 1:2000 for WB |
| Anti-Mouse IgG (H+L), Superclonal™ Recombinant Secondary Antibody, Alexa Fluor™ 647 | Thermo Fisher | A28181 | 1:500 for IF |

Table S2 List of lentiviral particles used in this paper.

| Type | Gene name | Target sequence 5’-3’ | Vector | Source | MOI |
| --- | --- | --- | --- | --- | --- |
| Over-expression | UPF1 | cDNA: UPF1-201 (ENST00000262803.10) | pLenti-P2A-tGFP | Origene | 5 |
| Knock-down | XRN2 | TACATAGCTGATCGTTTAAAT | pLKO.1-CMV-neo | Sigma-Aldrich | 4 |
|  | EXOSC10 | sh1:AGTTGACTTGGAGCACCACTCTTACAGGA  sh2:TGATACTCATCAGGCAGCACGCCTTCTTA  sh3:TCAGGCAGCGAAGTTCGATCCATCAACCA  sh4:CTTACGACTACAGCCAGTCAGACTTCAAG | pGFP-C-shLenti | Origene | 4 (in total) |
|  | UPF1 | sh1:ATCACGGCACAGCAGATCAACAAGCTGGA  sh2:CACAACCAGATCAGGAACATGGACAGCAT  sh3:TGCCTGCGTGGTTTACTGTAATACCAGCA  sh4:CGTGAGAGCCTCATGCAGTTCAGCAAGCC | pGFP-C-shLenti | Origene | 4 (in total) |
|  | STAU1 | CCAACAGTGATCTGTATTCTT | pLKO.1-CMV-neo | Sigma-Aldrich | 4 |
| Negative Control | Scramble | Non-target: TATGTGCGGCAAACCAAGCG | pLKO.1-CMV-neo | Sigma-Aldrich | 4 |
|  | Scramble | Non-target: GCGCGATAGCGCGAATATAC | pGFP-C-shLenti | Origene | 4 |
|  | Empty | / | pLenti-P2A-tGFP | Origene | 5 |

Table S3 List of probes used in the STORM experiments.

|  | | ID | Probe sequence 5’󠄐-3’ | Tm (℃) |
| --- | --- | --- | --- | --- |
| Single-labelling | | ATTO647n-(CAG)_8_ | ATTO647n-CAGCAGCAGCAG CAGCAGCAGCAG | 75.1 |
| Dual-labelling | Set 1 | ATTO550-(CAG)_8_ | ATTO550-CAGCAGCAGCAG CAGCAGCAGCAG | 75.1 |
|  |  | 5’-DMPK-1 | ATTO647n-CGCTCGGAGCGGTTGTGAACTG | 66.2 |
|  |  | 5’-DMPK-2 | ATTO647n-ACAGAACAACGGCGAACAGGAGCA | 64.7 |
|  |  | 5’-DMPK-3 | ATTO647n-ACTGCCACTTCAGCTGTTTCATCCTGT | 65.6 |
|  |  | 5’-DMPK-4 | ATTO647n-CCTTCCCGAATGTCCGACAGTGTCT | 64.3 |
|  |  | 5’-DMPK-5 | ATTO647n-AGAGGTGCTCCTTGTAGTGGACGAT | 64.2 |
|  |  | 5’-DMPK-6 | ATTO647n-GGTCCCCATTCACCAACACGTCC | 64.4 |
|  |  | 5’-DMPK-7 | ATTO647n-AGTCGGACCTCCTTAAGCCTCACC | 64.4 |
|  |  | 5’-DMPK-8 | ATTO647n-ACATGTTGGACAGGCAGCACCATG | 65.2 |
|  | Set 2 | ATTO550-(CAG)_8_ | ATTO550-CAGCAGCAGCAG CAGCAGCAGCAG | 75.1 |
|  |  | 3’-DMPK-1 | ATTO647n-GGGCCTTTTATTCGCGAGGGTC | 65.4 |
|  |  | 3’-DMPK-2 | ATTO647n-CGGGGTCTCAGTGCATCCAAAACG | 65.3 |
|  |  | 3’-DMPK-3 | ATTO647n-GGGTCCTGTAGCCTGTCAGCGA | 64.8 |
|  |  | 3’-DMPK-4 | ATTO647n-TAAATATCCAAACCGCCGAAGCGGGC | 64.6 |
|  |  | 3’-DMPK-5 | ATTO647n-CCCCAGAGCAGGGCGTCATGCA | 65.2 |
|  |  | 3’-DMPK-6 | ATTO647n-GGGGTGCGTGGAGGATGGAACAC | 65.5 |
|  |  | 3’-DMPK-7 | ATTO647n-GCAGTTTGCCCATCCACGTCAGG | 66.1 |
|  |  | 3’-DMPK-8 | ATTO647n-GCCTCAGCCTGGCCGAAAGAAAGA | 64.9 |
| Scramble | | ATTO647n-Scramble | ATTO647n-AGTCCTGCAATGTGACATCGAGTG | 64.3 |

**Characterisation of MBNL-deficient DM1 cell lines: P4C1 and P4B6**

In this section, we elucidated the process and outcomes of CRISPR/Cas9-mediated knock-out targeting exon 4 of the *MBNL1* gene and exon 6 of the *MBNL2* gene. Our investigation involved transfection of cells with plasmids encoding Cas9 protein and guide RNAs, followed by the selection and cultivation of stably modified cell lines. Subsequently, we conducted DNA extraction and PCR amplification to assess the efficacy of genome editing at the anticipated target sites within the *MBNL1* and *MBNL2* genes. We employed distinct PCR primer pairs to amplify fragments encompassing the target regions, located within exon 4 of the *MBNL1* gene and exon 6 of the *MBNL2* gene. The anticipated outcome for unmodified target sites was the generation of a 572-bp fragment from *MBNL1* genomic DNA and a 554-bp fragment from *MBNL2* genomic DNA. For further characterisation, Sanger sequencing was performed on the PCR products. Through these endeavours, we successfully generated two modified cell lines, namely P4C1 and P4B6. Notably, our Sanger sequencing results revealed that in the context of *MBNL1* gene editing, P4C1 cells exhibited a homozygous deletion of four nucleotides within both alleles of the *MBNL1* gene, while P4B6 cells displayed a homozygous deletion of one nucleotide within both alleles. Similarly, concerning *MBNL2* gene editing, P4C1 cells exhibited a homozygous deletion of four nucleotides in both alleles of the *MBNL2* gene, while P4B6 cells showcased a homozygous deletion of two nucleotides within both alleles.

These deletions unequivocally induced frameshift mutations within the *MBNL1* and *MBNL2* gene, and Consequently premature stop codons (PTCs) were introduced into their respective mRNAs. Furthermore, the frameshift mutations may have caused complete skipping of exons. Ultimately, the introduction of PTCs and/or exon skipping can disrupt the overall structure and stability of the mRNA molecules. This disruption may lead to the generation of truncated MBNL1 and MBNL2 proteins (as observed in P4B6 cells with truncated MBNL1 and truncated MBNL2, and P4C1 cells with truncated MBNL2), or degradation and destabilisation of the mRNA (as observed in P4C1 cells with the complete absence of MBNL1 protein) via nonsense-mediated decay (NMD) if the PTC occurs before the last exon-exon junction of the transcript [1]. Alternatively, other faulty mRNA surveillance mechanisms may also contribute to the degradation.

Table S4 gRNA sequences targeting *MBNL1* and *MBNL2*.

|  | Target sequence  5’-3’ | Exon | Efficiency |
| --- | --- | --- | --- |
| MBNL1 | CAACGTGGCAATTGCAACCGAGG | 4 | 0.69 |
| MBNL2 | GCTGGGGTTAAAGACCGCGCTGG | 6 | 0.71 |

The efficiency values were calculated by the CRISPR Design software: https://chopchop.cbu.uib.no/. The protospacer adjacent motif (PAM) sequences are indicated in red.

Fig. S1 Genome Editing of the Human MBNL Locus.

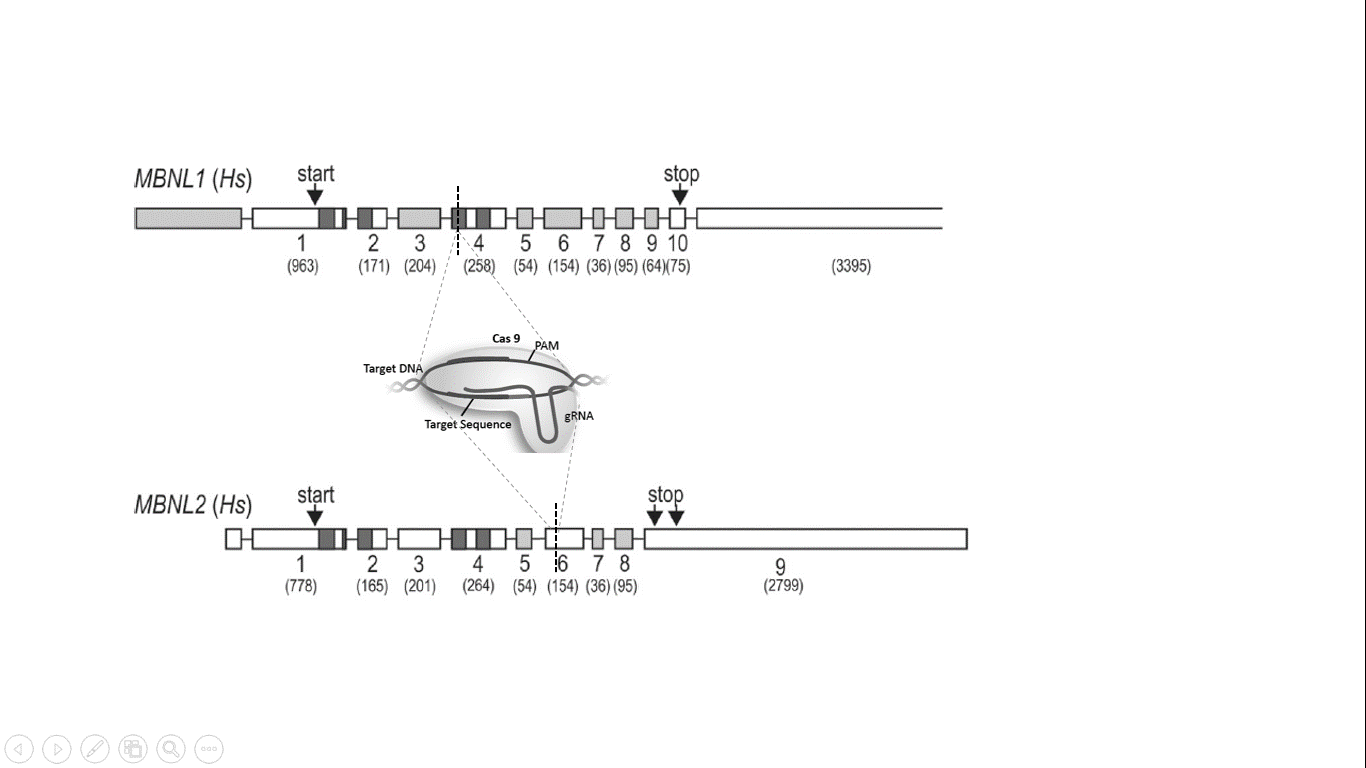

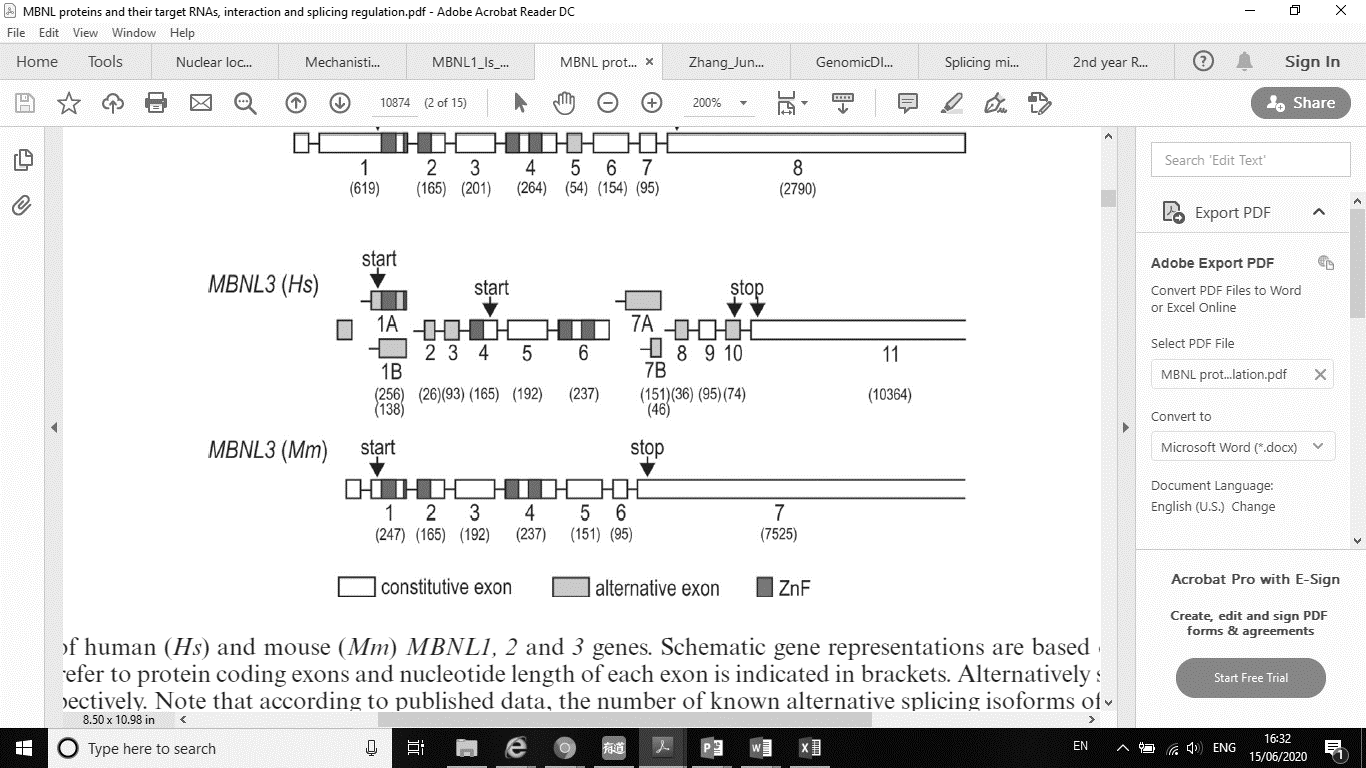

The CRISPR-Cas9 target sites (shown as black dashes) are in the exon 4 (258 nt) of *MBNL1* and in the exon 6 (154 nt) of *MBNL2* respectively. Exon numbers refer to protein coding exons and nucleotide length of each exon is indicated in brackets. Alternatively spliced exons and ZnFs are marked light grey and dark grey respectively.

Fig. S2 Assessment of double knock-out of *MBNL1* and *2* by Western blotting.

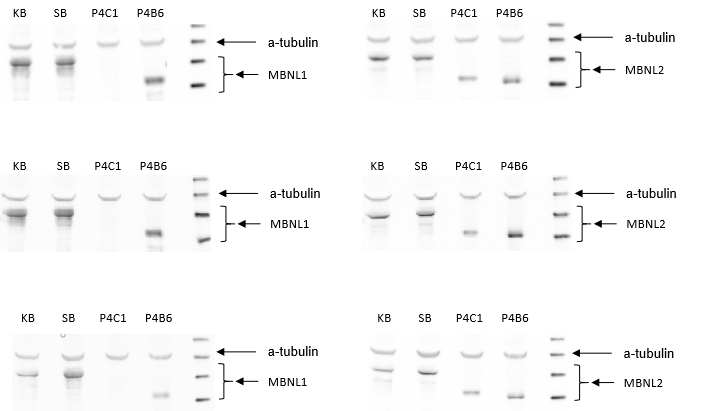

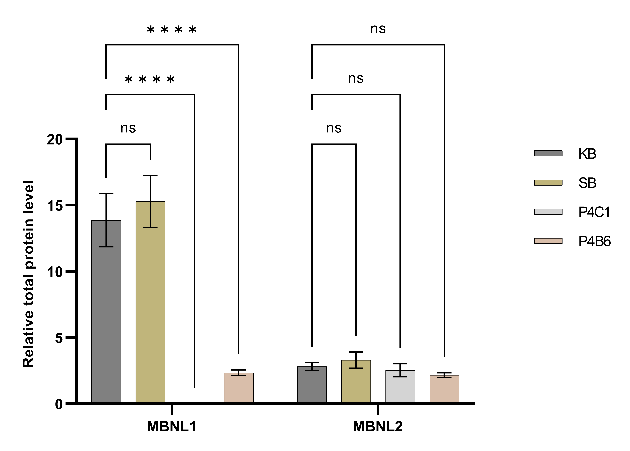

(A) Total protein extracts from KB (a DM1 patient cell line), SB (a non-DM control cell line), as well as P4C1 and P4B6 (MBNL-deficient DM1 cell lines) were analysed by Western blotting to detect the expression of MBNL1 and MBNL2 proteins. α-tubulin was used as a loading control. (B) Bar chart shows relative total MBNL protein level in these cell lines, normalised against the protein level of α-tubulin. The experiment was performed in triplicate to ensure the reproducibility of the results.

MBNL is an RNA-binding protein that regulates its own alternative splicing through a self-regulatory mechanism. There are two primary splicing variants for *MBNL1* mRNA: Variant 1, which lacks alternative exon 7, produces a protein with an approximate molecular weight of 42 kDa; variant 2, lacking alternative exon 5, produces a protein with an approximate molecular weight of 41 kDa. For *MBNL2* mRNA, the two major splicing variants are: variant 1, lacking alternative exon 7, produces a protein weighing approximately 40 kDa; variant 2, which skips both alternative exons 5 and 7, produces a protein weighing approximately 38 kDa.

The specific targeting of exon 4 in the *MBNL1* gene by CRISPR/Cas9 resulted in either complete silencing or significant reduction of MBNL1 protein expression in P4C1 and P4B6 cells. This finding aligns with previous studies by Hale, M.A., et al. (2018), which demonstrated that the ZnF3-4 domain encoded by exon 4 enhances the stability of MBNL1 [2]. Furthermore, the ZnF3-4 domain is responsible for binding to target mRNAs and enabling MBNL1 to regulate their alternative splicing.

Through my RNA-Seq analysis, it was revealed that CRISPR/Cas9-mediated knock-out of *MBNL1* caused complete skipping of exon 4 (**Fig. S3** A and B, **Table S5**), potentially disrupting the normal self-regulation mechanism of MBNL1. Without the ZnF3-4 domain encoded by exon 4, MBNL1 may lose its ability to bind and regulate its own mRNA effectively. Consequently, dysregulation in the alternative splicing of *MBNL1* itself can lead to unintended mis-splicing events. In fact, I observed mis-splicing of *MBNL1* mRNA in P4C1 and P4B6 cells, where exons 5, 6, and 7 exhibited down-regulation (**Fig. S3** A and B, **Table S5**). These downstream exons are reliant on the proper functioning of MBNL1 and its regulatory interactions to be accurately included in the final mRNA transcript. Thus, the absence of functional MBNL1 due to mis-splicing can result in unintended skipping or exclusion of downstream exons.

On the other hand, the expression level of MBNL2 protein with C-terminal truncation remained unaffected in P4C1 and P4B6 cells compared to wild-type MBNL2 in DM1 cells, despite specific targeting of exon 6 in the *MBNL2* gene by CRISPR/Cas9. Similar to targeting exon 4 in the *MBNL1* gene, which caused complete skipping of exon 4 and depletion of downstream exons, targeting exon 6 in the *MBNL2* gene led to the skipping of exon 6 and down-regulation of downstream exon, exon 9 (**Fig. S3** C and D, **Table S5**). It's important to note that exon 6 is a constitutive exon which is essential for maintaining the proper structure and function of the protein, its depletion can result in altered protein structure, loss of functional domains, impaired RNA binding, and disrupted protein-protein interactions.

Fig. S3 Effect of *MBNL1* and *2* double knock-out on the *MBNL1* and *2* mRNA compositions

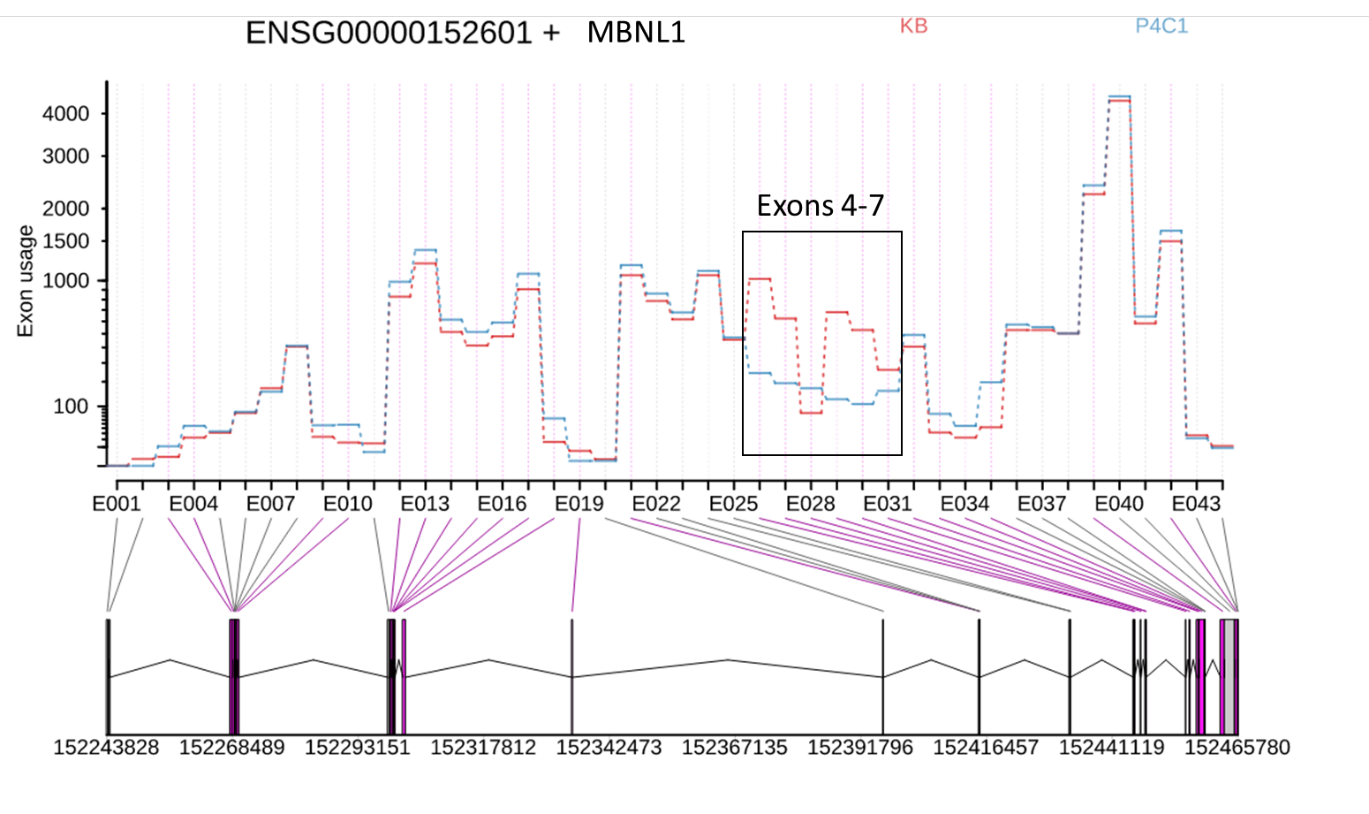

**A**

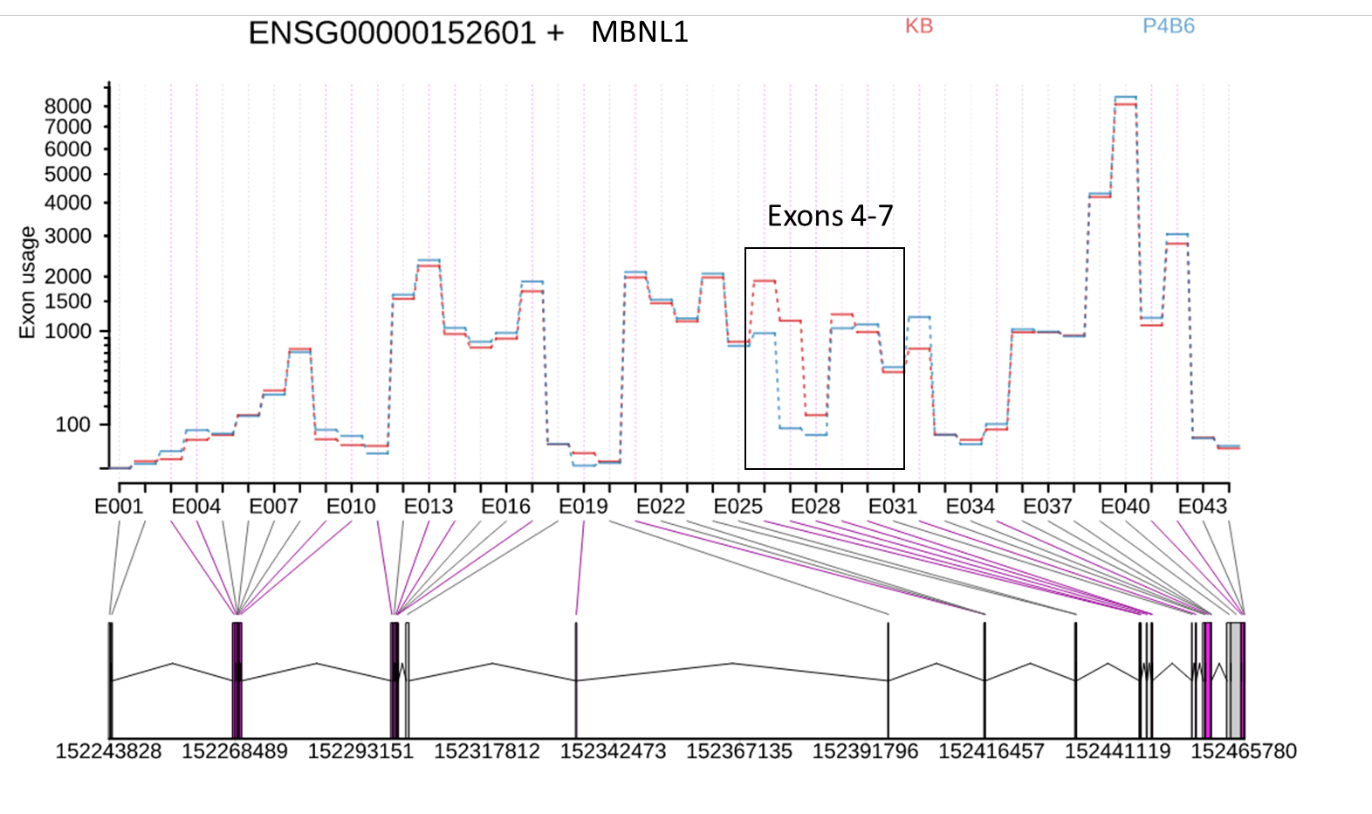

**B**

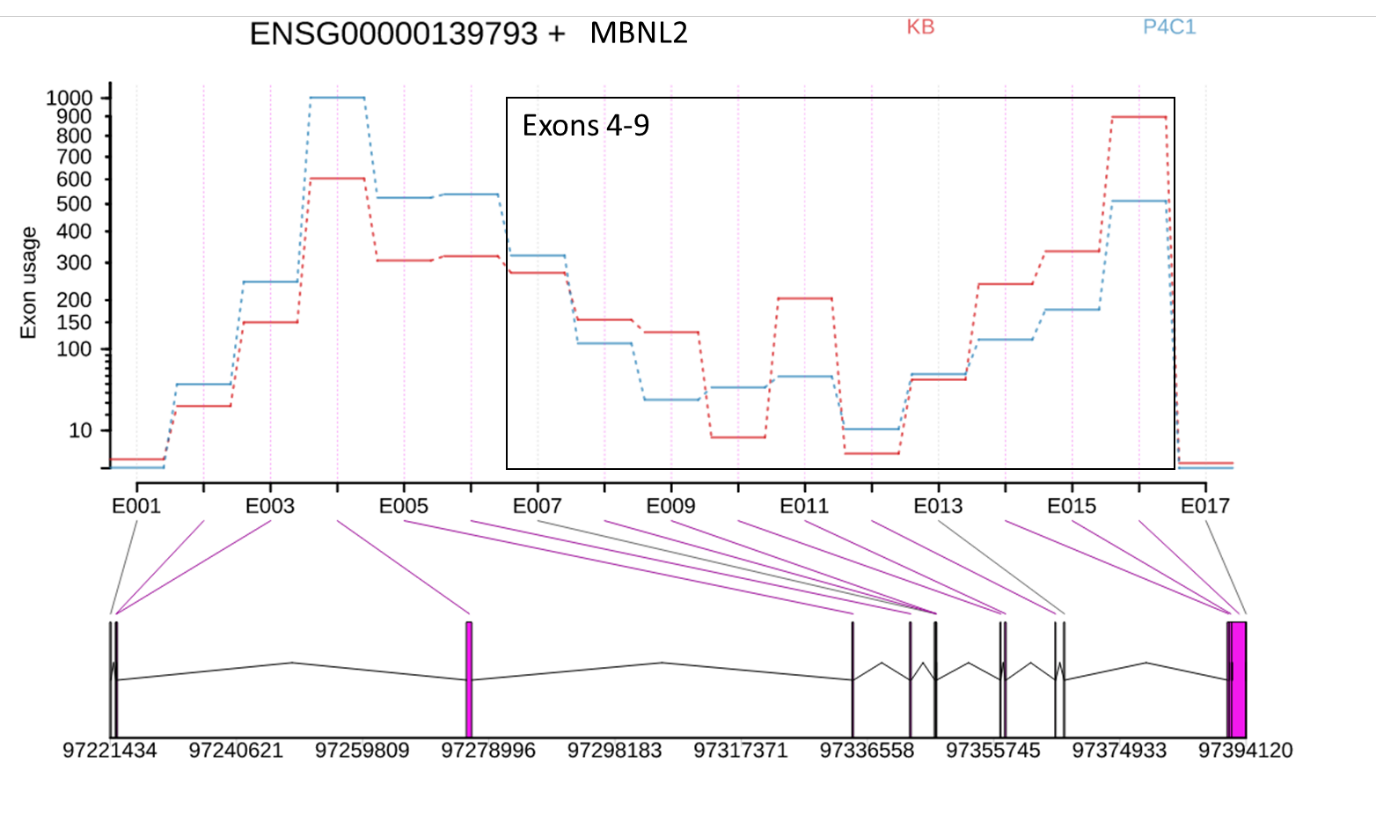

**D**

**C**

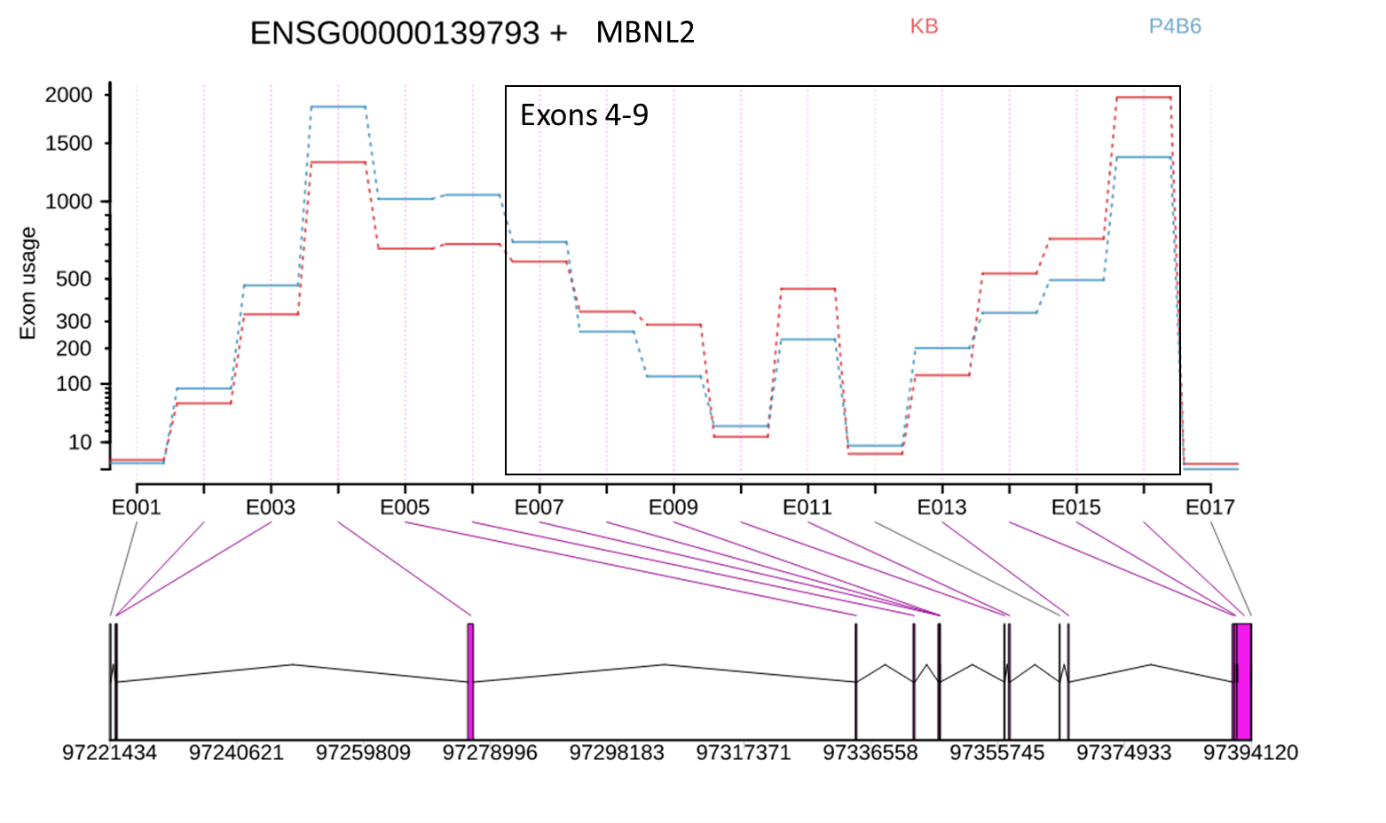
The figures depict the results of RNA-Seq data analysis using DEXSeq, revealing significant differential exon usage of *MBNL1* exon 4-7 (A-B) and *MBNL2* exon 4-7 (C-D) between the DM1 cell line (KB, shown in red) and the MBNL-deficient DM1 cell lines (P4C1 and P4B6, shown in blue). The exon numbering and corresponding counting bins used for quantification are as follows: *MBNL1* exon 4 (counting bin E026-027), exon 5 (counting bin E028), exon 6 (counting bin E029-030), and exon 7 (counting bin E031); *MBNL2* exon 4 (counting bin E007-009), exon 5 (counting bin E010), exon 6 (counting bin E011), exon 7 (counting bin E012), exon 8 (counting bin E013) and exon 9 (counting bin E014-E016). The top panel of the figure presents the fitted values of exon usage based on a linear model, after accounting for overall changes in gene expression. The bottom panel displays the flattened gene model. The purple highlighted counting bin exons indicate regions of significant differential exon usage. To ensure reproducibility, the experiment was performed in triplicate.

Table S5 Aberrantly spliced exons of *MBNL1* and *2* identified in P4C1 and P4B6.

| MBNL | Exon | Expression | Log_2_foldchange P4C1/KB or P4B6/KB |
| --- | --- | --- | --- |
| P4C1- MBNL1 | Exon 4 (encoding ZnF3-4) | ↓ | -2.06 (padj=7.61E-222) |
|  | Exon 5 | ↑ | 1.10 (padj=1.73E-14) |
|  | Exon 6 | ↓ | -2.44 (padj=3.07E-121) |
|  | Exon 7 | ↓ | -0.71 (padj=1.94E-3) |
| P4B6- MBNL1 | Exon 4 (encoding ZnF3-4) | ↓ | -0.97 (padj=1.09E-185) |
|  | Exon 5 | ↓ | -1.34 (padj=1.07E-14) |
|  | Exon 6 | — |  |
|  | Exon 7 | — |  |
| P4C1- MBNL2 | Exon 4 (encoding ZnF3-4) | — |  |
|  | Exon 5 | ↑ | 2.80 (padj=1.24E-18) |
|  | Exon 6 | ↓ | -1.79 (padj=2.41E-34) |
|  | Exon 7 | ↑ | 2.87 (padj=4.52E-4) |
|  | Exon 8 | — |  |
|  | Exon 9 | ↓ | -1.05 (padj=7.39E-24) |
| P4B6- MBNL2 | Exon 4 (encoding ZnF3-4) | — |  |
|  | Exon 5 | ↑ | 0.81 (padj=0.047) |
|  | Exon 6 | ↓ | -0.95 (padj=4.49E-24) |
|  | Exon 7 | — |  |
|  | Exon 8 | ↑ | 0.73 (padj=6.37E-07) |
|  | Exon 9 | ↓ | 0.65 (padj=6.25E-28) |

“↓” indicates down-regulation, “↑” indicates up-regulation, and “—” indicates no significant change. N=3, adjusted P values (padj) of <0.05 were considered to be statistically significant.

**Fig. S4 Speculation of the MBNL mRNA compositions.**

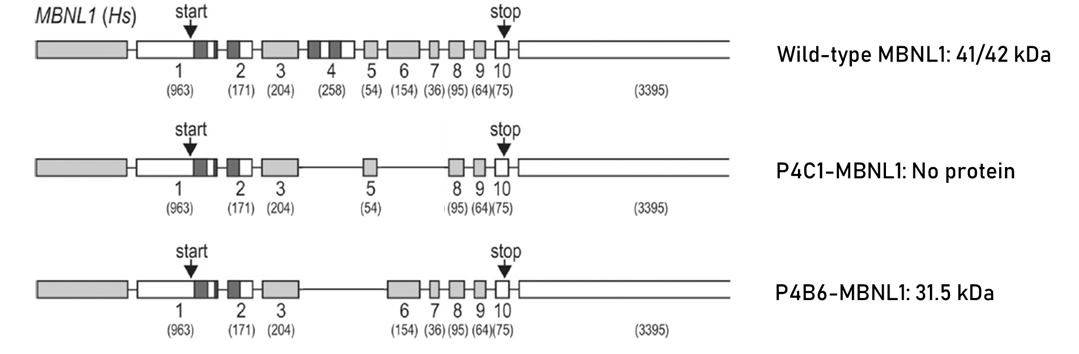

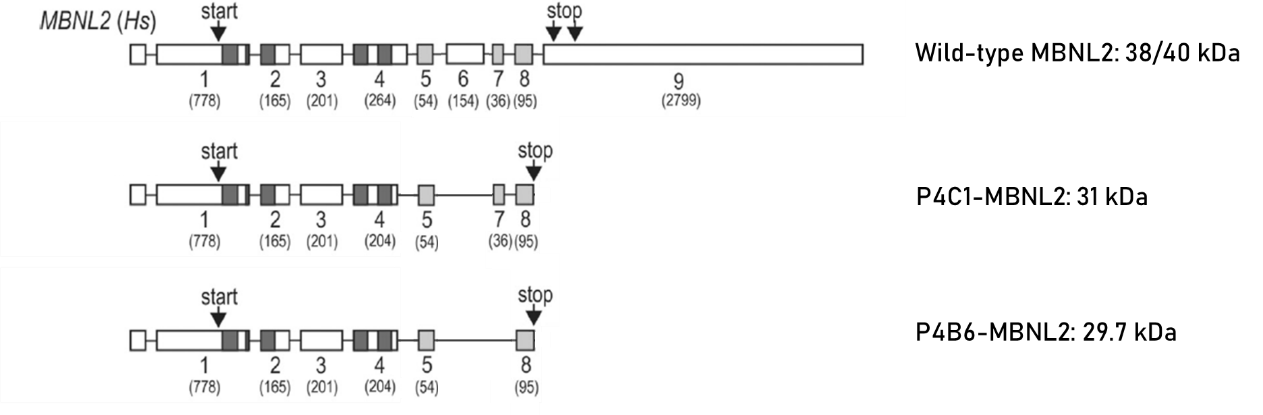

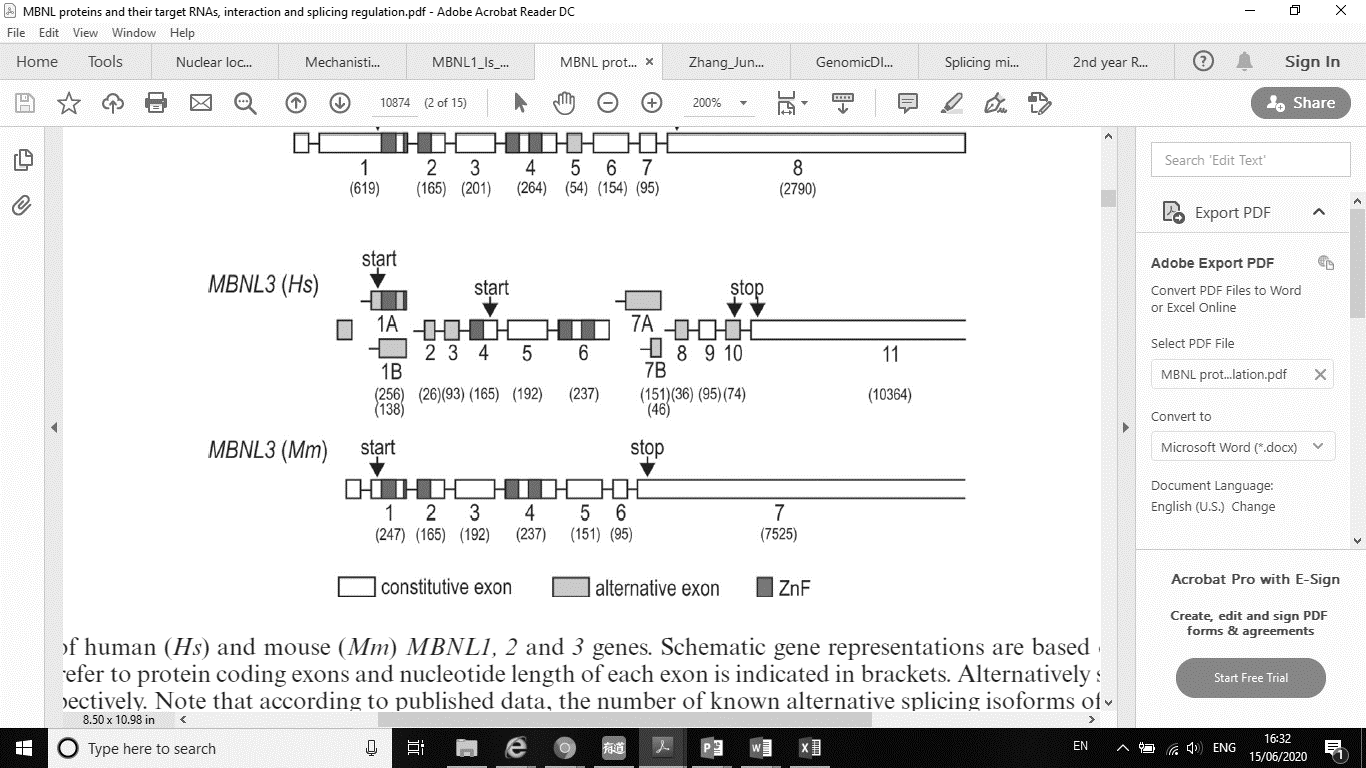

The diagrams show the speculated *MBNL1* and *2* mRNA compositions and molecular weights of the corresponding protein products in the two MBNL-deficient cell lines (P4C1 and P4B6). Alternatively spliced exons and ZnFs are marked light grey and dark grey, respectively.

**Nanopore DNA sequencing**

The length of CTG repeat expansions in the *DMPK* gene is positively correlated with the severity of myotonic dystrophy and increases with successive generations, contributing to anticipation. Southern blot analysis often reveals a broad smear of genomic DNA restriction fragments containing the expanded CTG repeats, indicating somatic mosaicism [3, 4]. Somatic mosaicism can occur due to the instability of the CTG repeat expansion in the *DMPK* gene during cell division. This means that some cells in an affected individual may have a different number of CTG repeats than other cells, leading to variability in symptoms and disease severity and difficulty in measuring allele size.

Although CTG repeat length correlates with disease severity and age of onset, the correlations are imprecise, and measuring allele size alone cannot accurately predict prognosis or anticipate disease progression. Technical difficulties in defining average allele length and measuring small shifts in repeat length over time contribute to this variation [3].

To further our understanding of the correlation between CTG repeat length and anticipation in myotonic dystrophy, more comprehensive and accurate studies are required.

We utilised the Cas9-assisted targeting of chromosome segments (CATCH) technique in conjunction with Nanopore DNA sequencing to accurately measure the CTG repeat lengths within the human *DMPK* gene. This approach enables precise targeting of specific chromosome regions and allows for high-throughput sequencing with long read lengths, enabling more accurate and comprehensive analysis of CTG repeat length variability. The following target sequences by CRISPR/Cas9 were used for isolation of *DMPK* locus: 5’ to CTG repeat region, 5’-GGGCGTGTATAGACACCTGG-3’ and 3’ to the CTG repeat region, 5’-TGCGAACCAACGATAGGTGG-3’. The sgRNA activity was tested in vitro prior to Nanopore sequencing (**Fig. S5**).

**Fig. S5 Assessment of the sgRNA activity by Cas9 in vitro assays.**

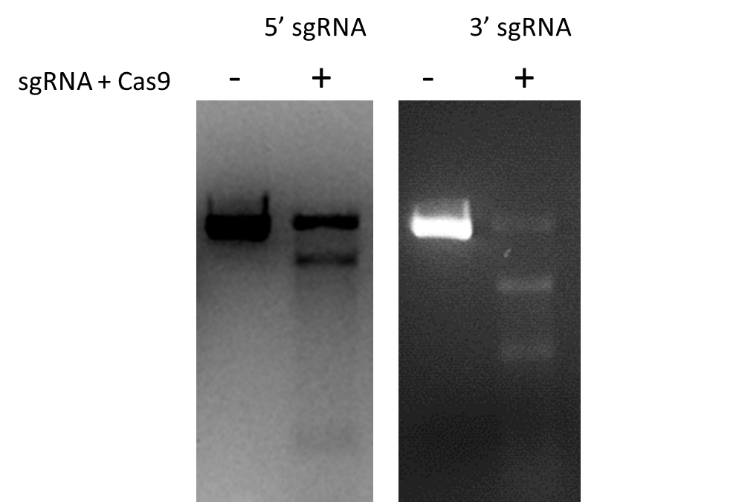

Ethidium bromide-stained gel showing the sgRNA activity following the performance of Cas9 in vitro assays. 5’ sgRNA and 3’ sgRNA recognise the sequences 5’ and 3’ to the CTG region in the *DMPK* gene, respectively.

The *DMPK* gene was cleaved at two sites flanking the CTG repeat region, according to the complementary gRNAs, and the fragment was isolated by PFGE. A large amount of target DNA was extracted from KB, P4C1 and P4B6 cells allowing sequencing library preparation from native DNA without pre-amplification. Nanopore sequencing was then used to analyse the isolated fragments and measure the CTG repeat length. Here, Nanopore sequencing has preliminarily proven to be an effective approach to estimate precise tandem repeat length in each cell with its ability to produce long, single-molecule reads.

Nanopore sequencing generated a total of 213 raw reads in P4C1, out of which 162 were mapped to chromosome 19. Among the mapped reads, 143 contained CTG repeats within the *DMPK* gene. Similarly, P4B6 produced 81 raw reads, with 49 reads successfully mapped to chromosome 19, and 33 of these mapped reads contained CTG repeats. For KB, a total of 138 raw reads were obtained, with 98 mapped to chromosome 19, and 78 of these mapped reads contained CTG repeats.

The analysis of the reads obtained from these three cell lines revealed important findings. Firstly, a substantial amount of reads (points in the bottom rows in the **Fig. S6**) contained CTG repeats ranging from 6 to 8 units in these cell lines. This indicates that these reads represent the non-expanded allele of the *DMPK* gene, which appears to be somatically completely stable in these cell lines. Furthermore, it is evident that all three cell lines exhibited expanded CTG repeats in the *DMPK* gene (**Fig. S6**). P4C1 displayed CTG repeat lengths ranging from 6 to 1553 units, with a median of 940 units for the expanded repeat lengths (>50 units). P4B6 exhibited CTG repeat lengths ranging from 6 to 346 units, with a median of 291 units for the expanded repeat lengths. Additionally, KB demonstrated CTG repeat lengths ranging from 6 to 2574 units, with a median of 284 units for the expanded repeat lengths. Notably, the sizes of CTG repeats varied among the cell lines, indicating somatic mosaicism [3, 4] in DM1 cells. The median values provide valuable information regarding the central tendency of the expanded repeat lengths in each cell line.

**Fig. S6 Distribution of the length of CTG repeats in DM1 cell lines.**

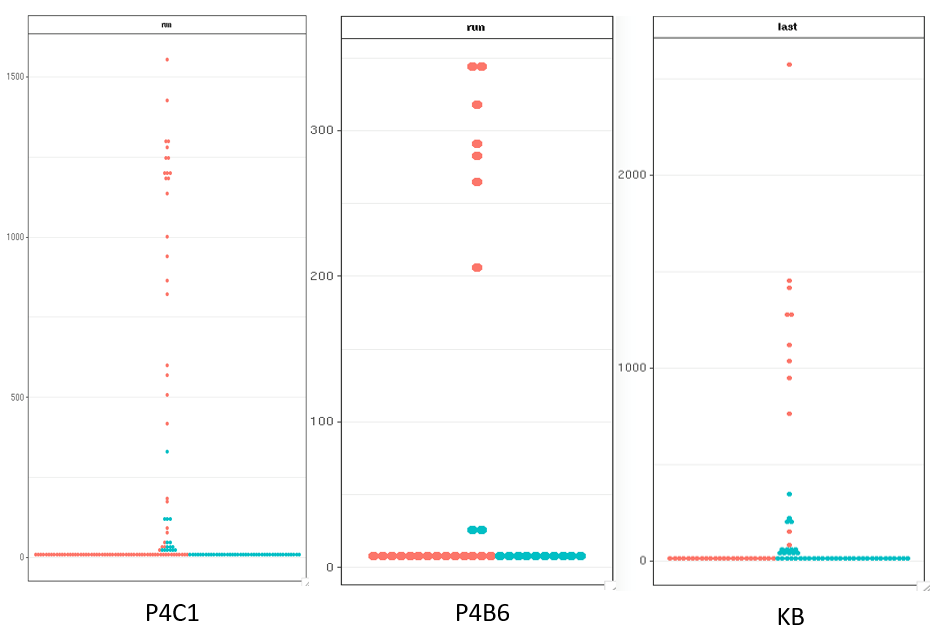

Y-axes indicate the number of CAG∙CTG repeat units. Sequence reads from the forward and reverse strand of DNA are shown in orange and blue, respectively.

These results suggest that Nanopore sequencing is an effective method for accurately estimating precise tandem repeat lengths with its capability of generating long, single-molecule reads. However, there was observed bias towards the forward strand in the nanopore sequencing data. One possible explanation for this bias would be that the library preparation protocol for Nanopore sequencing involves fragmenting the DNA and attaching adapters to the ends of the fragments. If there is bias during this step, such as preferential ligation of the adapters to one end of the DNA fragment, this could result in more reads being generated from one strand than the other. Another possibility could be base-calling errors. The base-calling software used to generate sequence data from the raw nanopore signal can introduce errors. If the base-caller is more accurate for one strand than the other, this could result in more reads being generated from the more accurately base-called strand.

Despite the strand bias, Nanopore sequencing provides several benefits compared to other methods such as PCR-based methods or Southern blot analysis: 1) Long-read sequencing enables the detection of complex repeat structures, including interruptions and expansions in non-canonical regions, that may be missed by other methods. 2) The direct sequencing of the CTG repeat region eliminates the need for amplification, reducing the risk of PCR artifacts and bias. 3) Nanopore sequencing is a powerful technique for individually measuring the CTG repeat length in a single strand of DNA. Due to somatic mosaicism, the size of CTG repeat expansions can vary significantly between cells or tissues. In contrast, PCR-based methods often display CTG repeat fragments as a broad smear band, making it challenging to accurately measure the length. By providing long-read sequencing data, Nanopore sequencing can overcome this limitation, enabling the detection of complex repeat structures and identifying potential somatic mosaicism. 4) The rapid turnaround time of Nanopore sequencing allows for the timely diagnosis of myotonic dystrophy and facilitates research studies.

Overall, Nanopore sequencing offers a promising solution for precise CTG repeat length determination in myotonic dystrophy patients, providing new insights into the genetic mechanisms and variability underlying the disorder.

Fig. S7 Distribution of large and micro (CUG)^exp^ foci from (CUG)^exp^ single-labelling experiments.

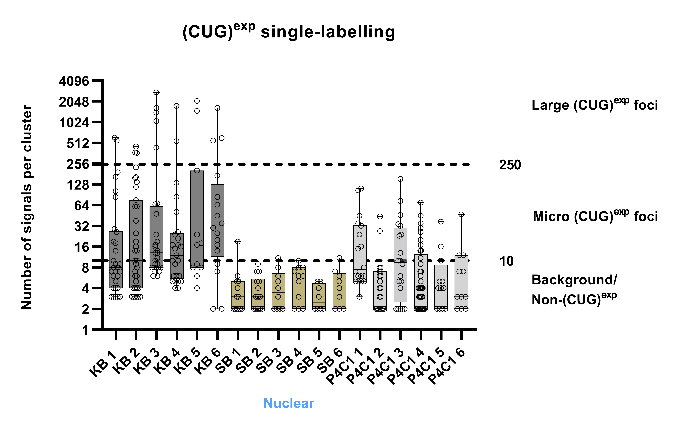

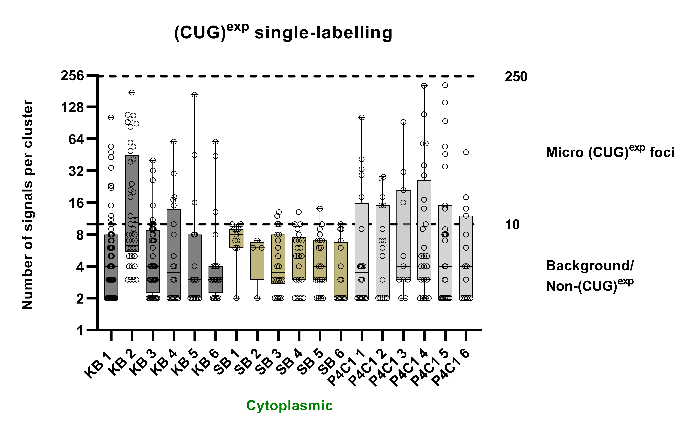

**B**

**A**

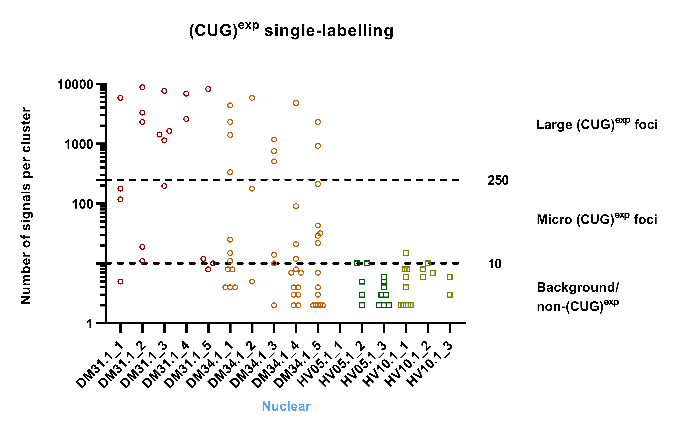

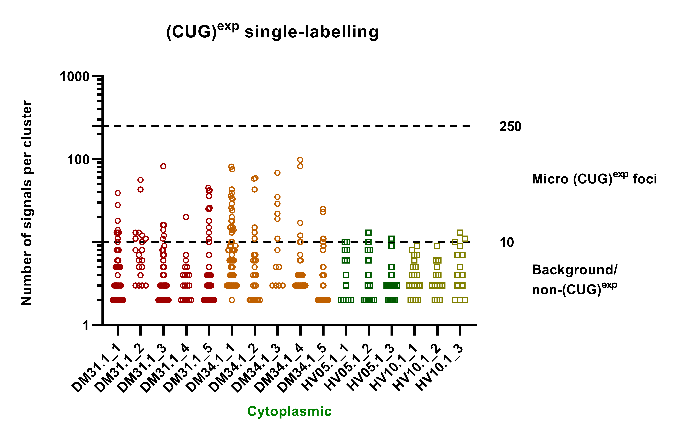

**D**

**C**

Diagrams showing the number of signals in each cluster identified with the (CAG)_8_ probe in the nucleus and in the cytoplasm. X axis, each column indicates a nucleus or a cytoplasm window. Y axis indicates the number of fluorophore signals per cluster. (CUG)^exp^ clusters containing >250 signals are considered as large (CUG)^exp^ foci; (CUG)^exp^ clusters containing 11-250 signals are considered as micro (CUG)^exp^ foci; (CUG)^exp^ clusters containing ≤ 10 signals are considered as background or non-(CUG)^exp^.

Table S6 Counts of large and micro (CUG)^exp^ foci and signals.

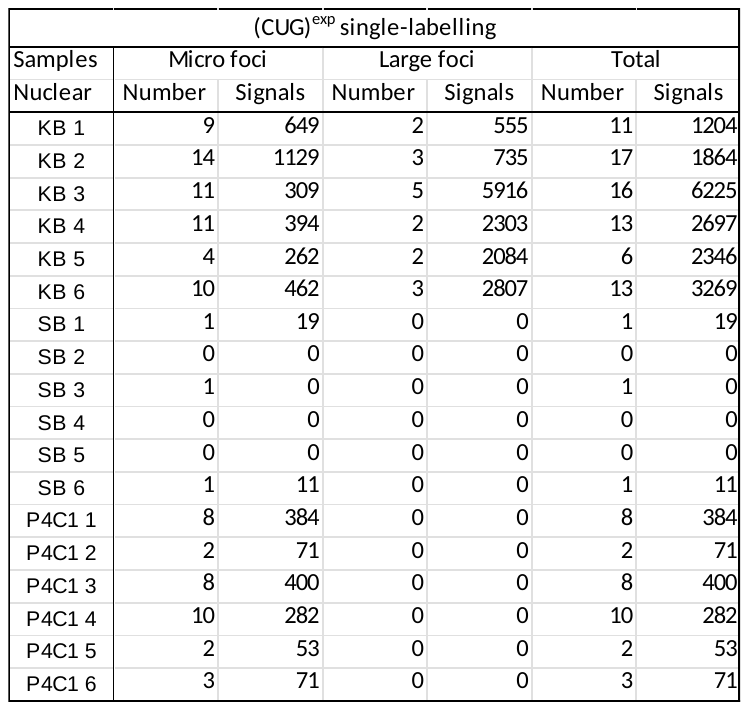

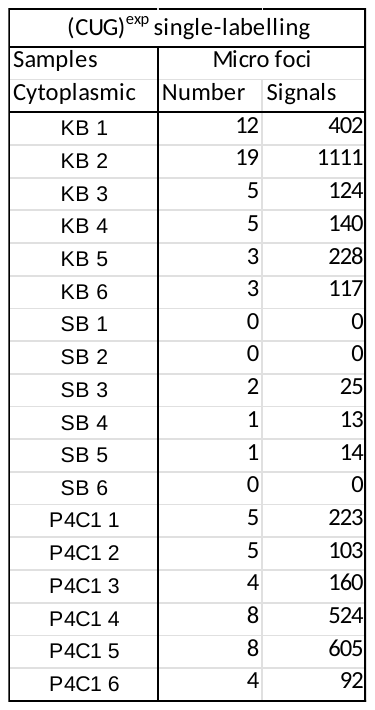

B

A

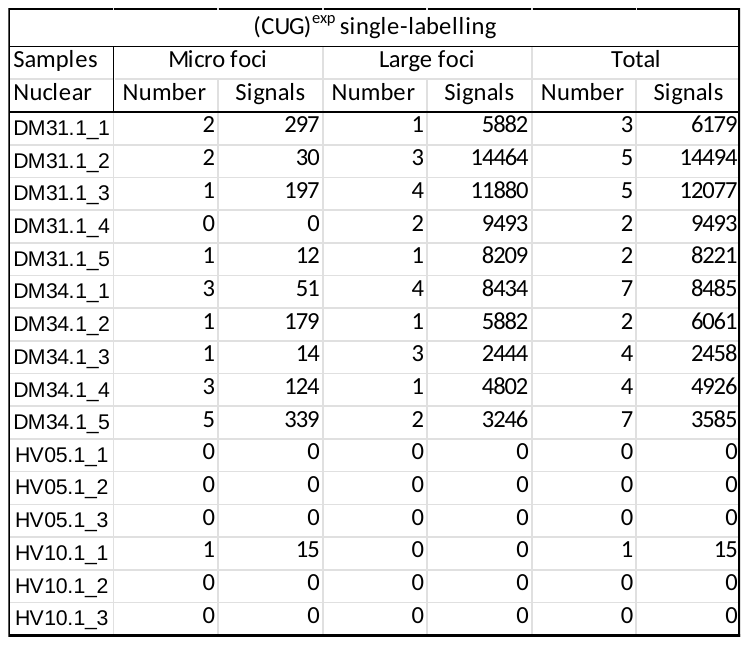

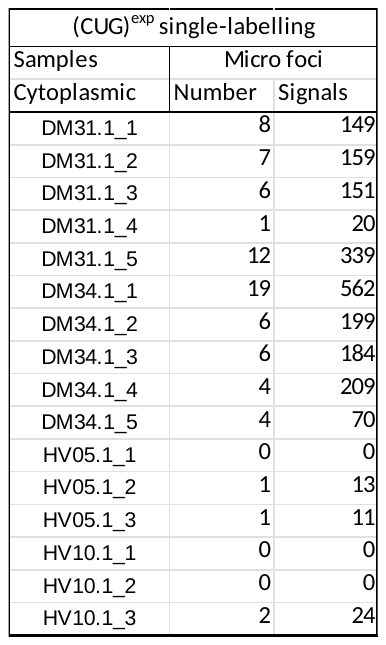

D

C

The “Number” columns show the counts of both large and micro (CUG)^exp^ foci in the nucleus of cell samples (Table A) and muscle biopsies (Table C), as well as in the cytoplasm of cell samples (Table B) and muscle biopsies (Table D). The “Signals” columns present the corresponding total count of probe signals detected within these foci.

Fig. S8 Distribution of 5’ or 3’ clusters from the 5’ or 3’ and (CUG)^exp^ dual-labelling experiments.

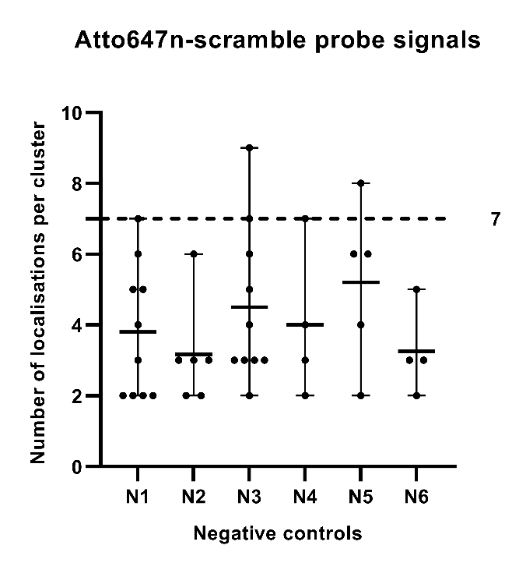

**A**

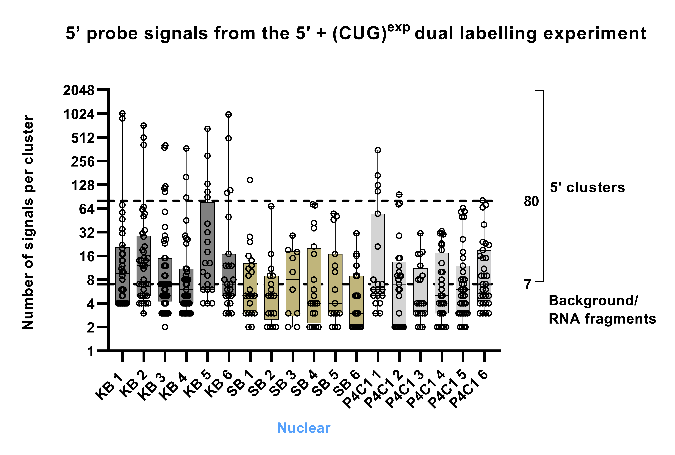

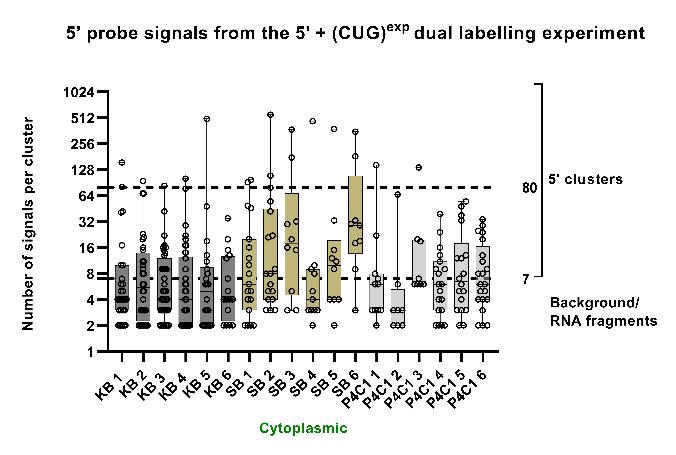

**C**

**B**

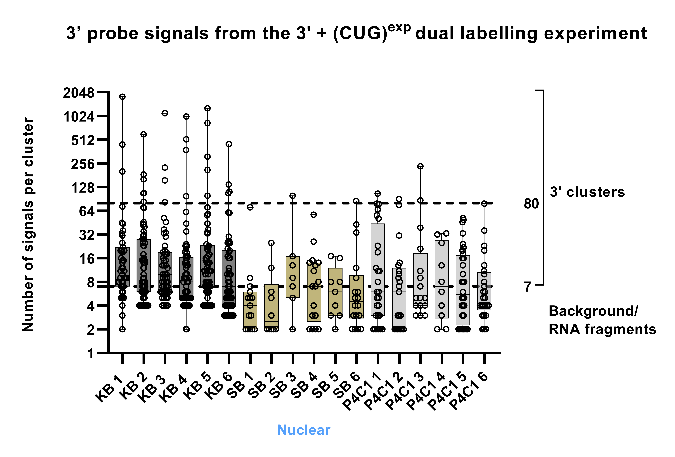

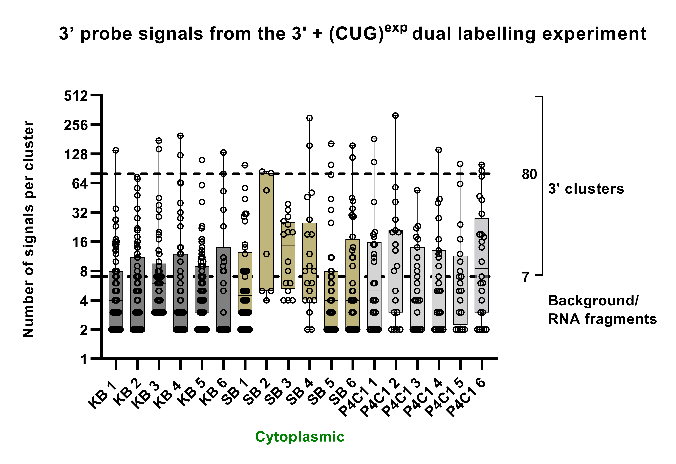

**E**

**D**

(A) Diagram showing the number of signals in each cluster that were detected by using the ATTO647n scramble probe under the same imaging conditions as the ATTO647n-labelled 5’ or 3’ single-copy probe sets. Each dot represents a single cluster, and the X-axis denotes each individual cell sample. (B-E) Diagrams showing the number of signals in each cluster identified with the 5’ or 3’ probe set in the nucleus and in the cytoplasm. X axis, each column indicates a nucleus or a cytoplasm window. Y axis indicates the number of fluorophore signals per cluster. Clusters identified with the 5’ or 3’ probe sets containing ≥8 signals are considered as 5’ or 3’ clusters; clusters identified with the 5’ or 3’ probe sets containing <8 signals are considered as background or RNA fragments.

Table S7 Large and micro (CUG)^exp^ foci co-localising with 5’ or 3’ signals.

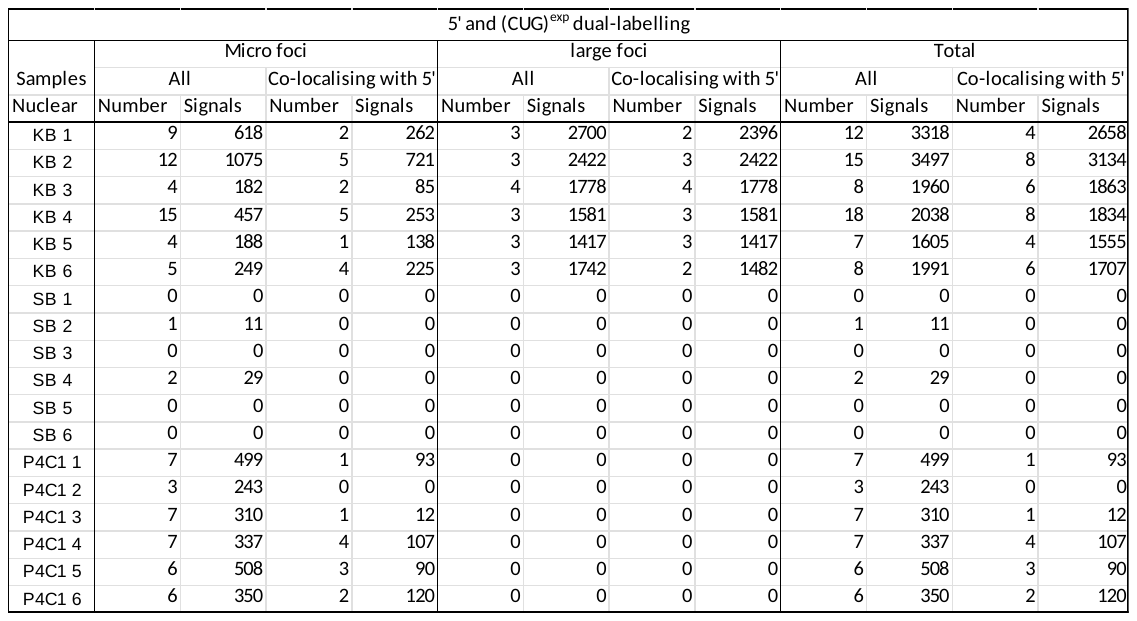

A

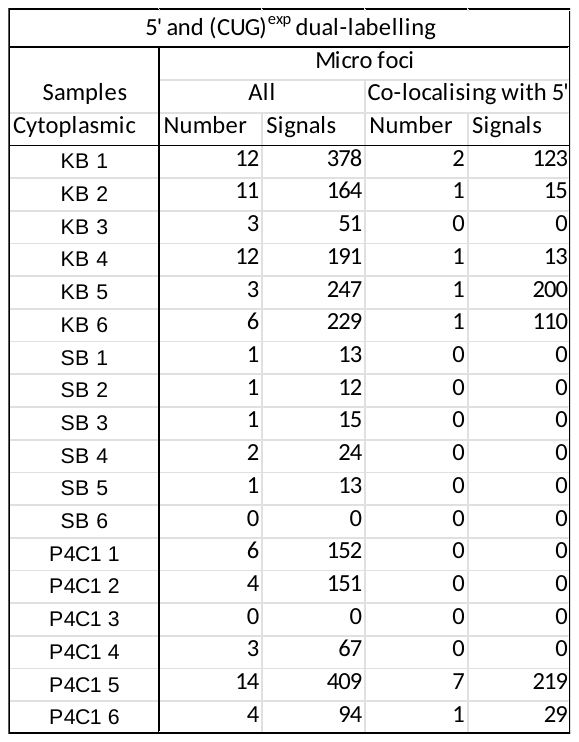

B

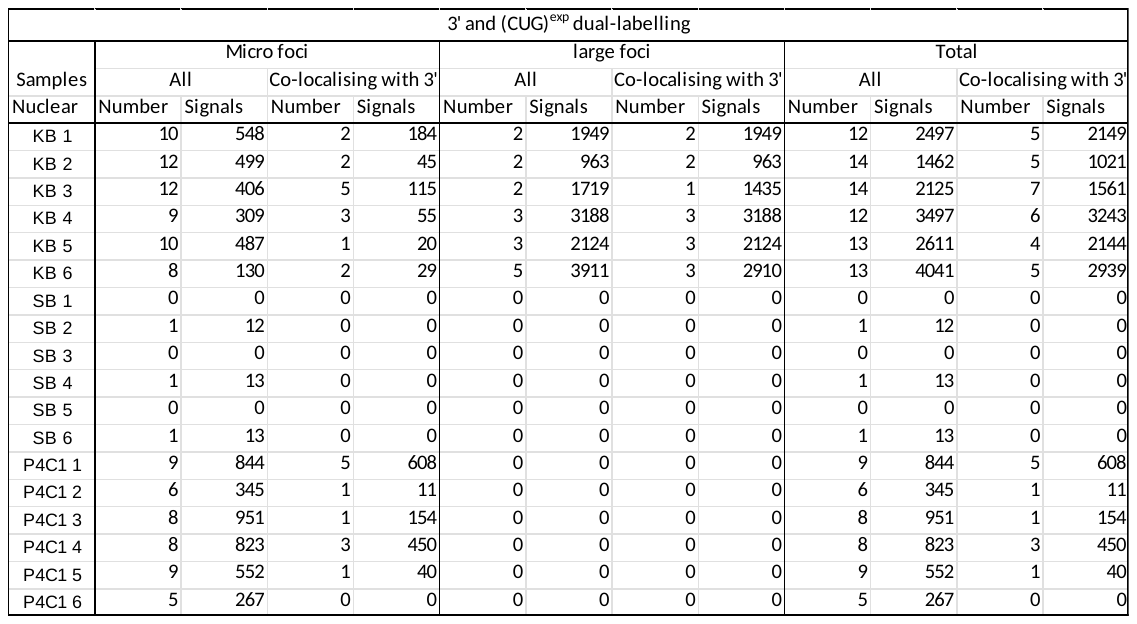

C

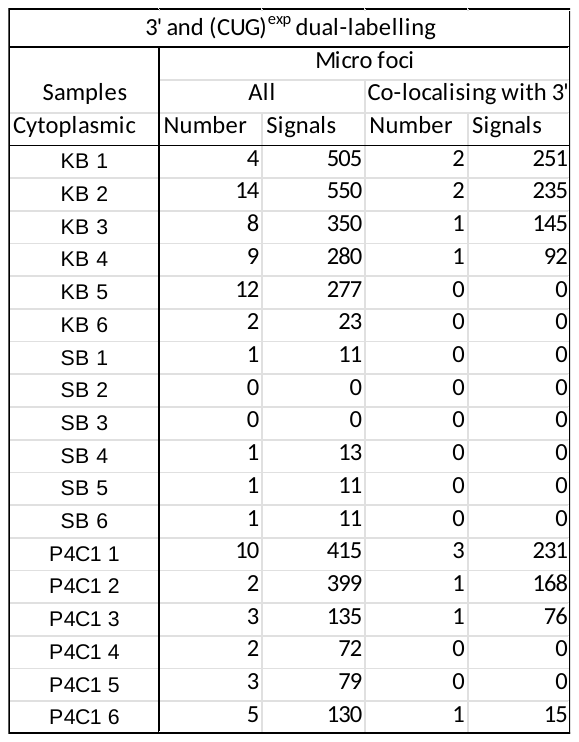

D

The “Number” columns show the counts of large and micro (CUG)^exp^ foci in the nucleus (Tables A and C) and the cytoplasm (Tables B and D), as well as the counts of foci that exhibit co-localisation with either 5’ probe signals or 3’ probe signals. The “Signals” columns present the corresponding total count of probe signals detected within these foci.

Fig. S9 MBNL1 and (CUG)^exp^ dual-labelling.

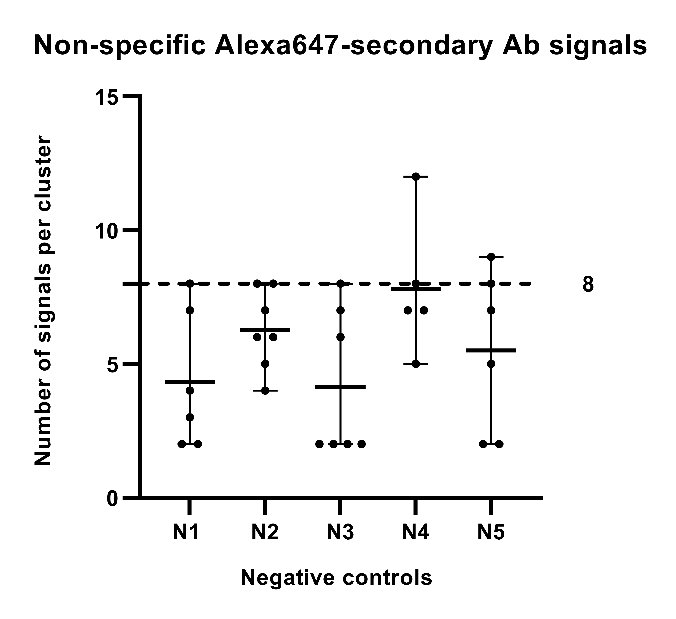

**A**

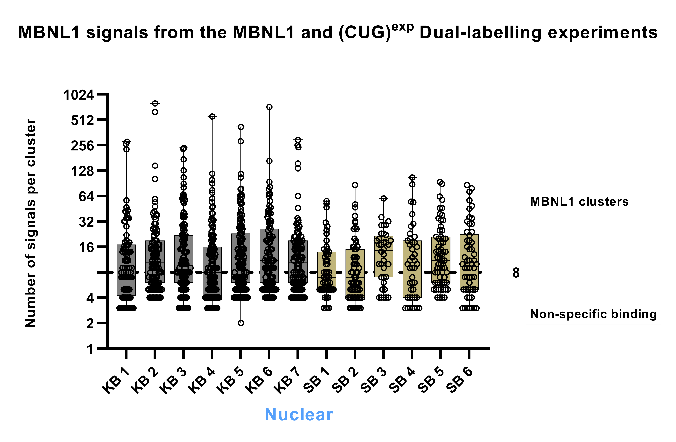

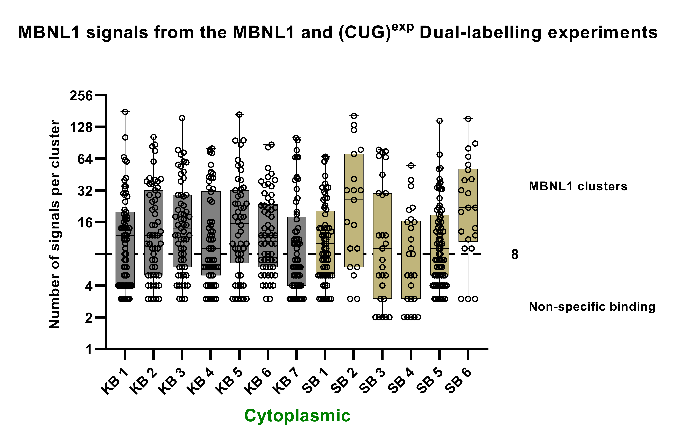

**B**

**C**

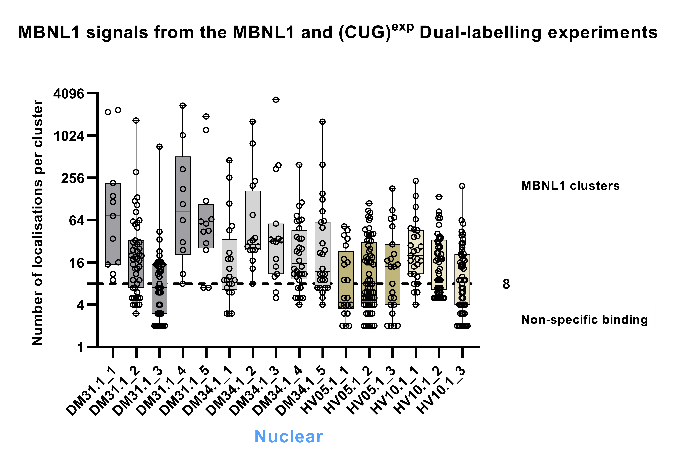

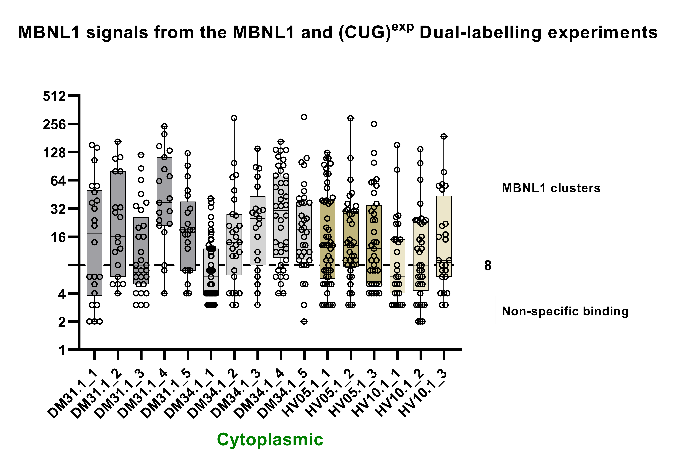

**D**

**E**

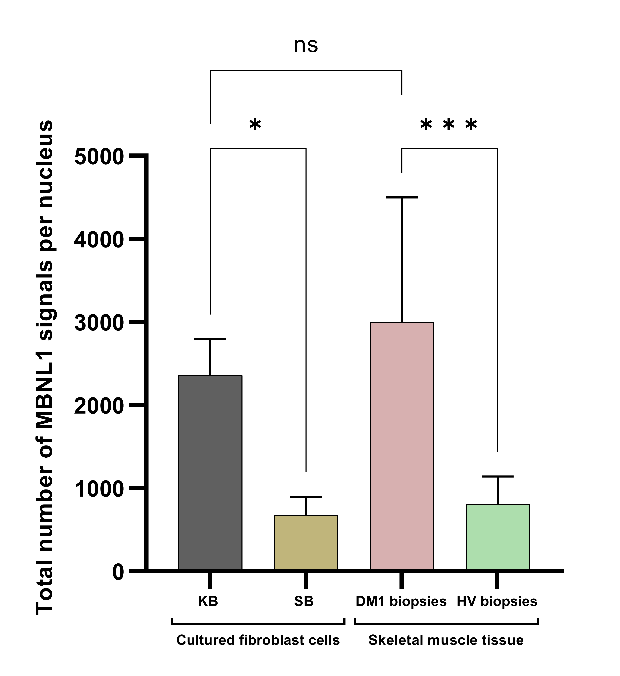

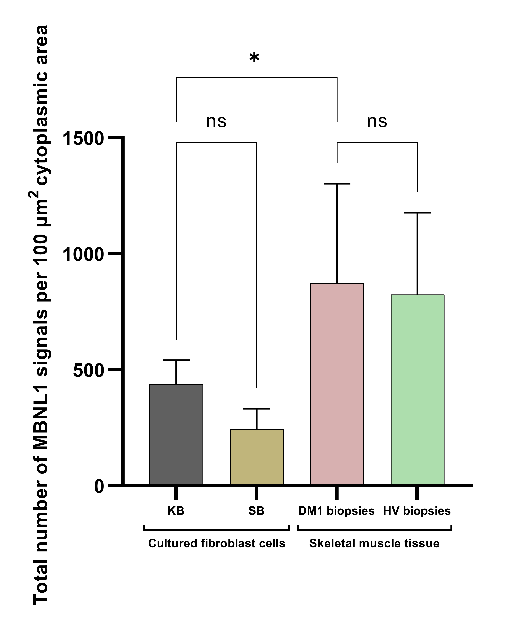

**H**

**G**

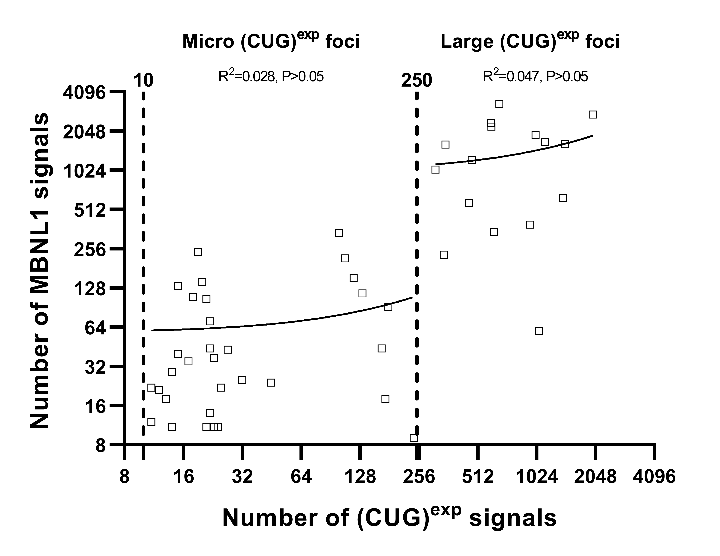

**J**

(A) Diagram showing the number of signals in each cluster that were detected by solely using the Alexa647n-labelled secondary antibody (without any primary antibody) under the same imaging conditions as when using both the anti-MBNL1 primary antibody and the Alexa647-labelled secondary antibody. Each dot represents a single cluster, and the X-axis denotes each individual cell sample. (B-E) Diagrams showing the number of signals in each cluster identified with the anti-MBNL1 antibody in the nucleus and in the cytoplasm. X axis, each column indicates a nucleus or a cytoplasmic window. Clusters containing ≥9 signals are considered MBNL1 clusters; clusters identified containing ≤8 signals are attributed to non-specific binding. (G-H) Histograms showing the average total number of MBNL1 signals detected per nucleus (G) and per 100 μm^2^ cytoplasmic area (H) in cultured DM1 (KB) and non-DM (SB) fibroblast cells, as well as muscle tissues from DM1 patients and healthy volunteers (HV). Statistical significance was determined by one-way ANOVA (mean ± SD, * for P<0.05, *** for P<0.001) or indicated as not significant (NS) for P>0.05. (J) Scatter plot showing (CUG)^exp^ foci and MBNL1 clusters co-localisations in DM1 biopsies. The X-axis indicates the number of signals in each co-localising (CUG)^exp^ foci and the Y-axis indicate the number of signals in each co-localising MBNL1 cluster. The data points are fitted with a linear regression line (solid line). The R^2^ values indicate the goodness of fit. A p-value of <0.05 indicates statistical significance.

Table S8 Counts of MBNL1 clusters and signals.

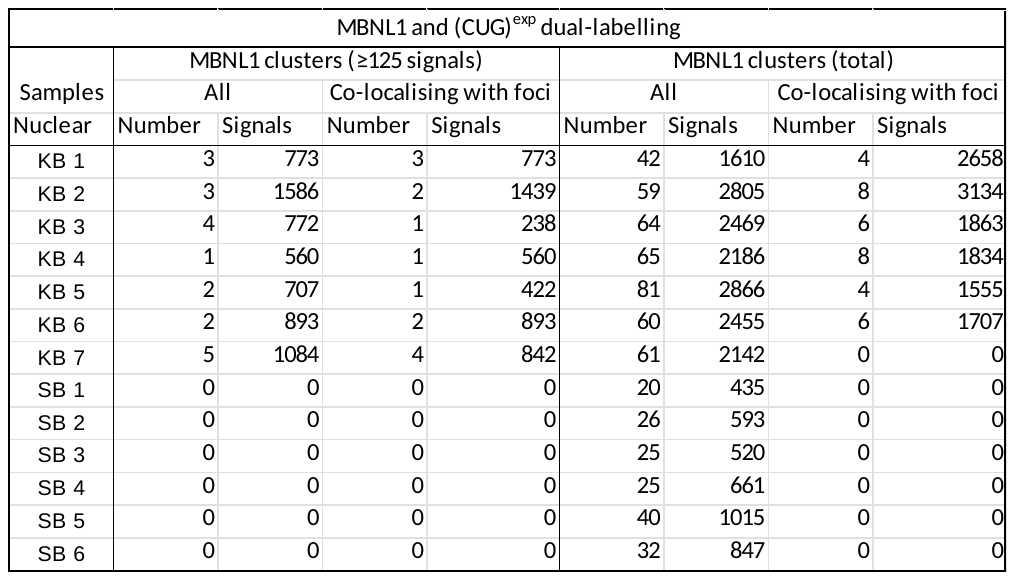

A

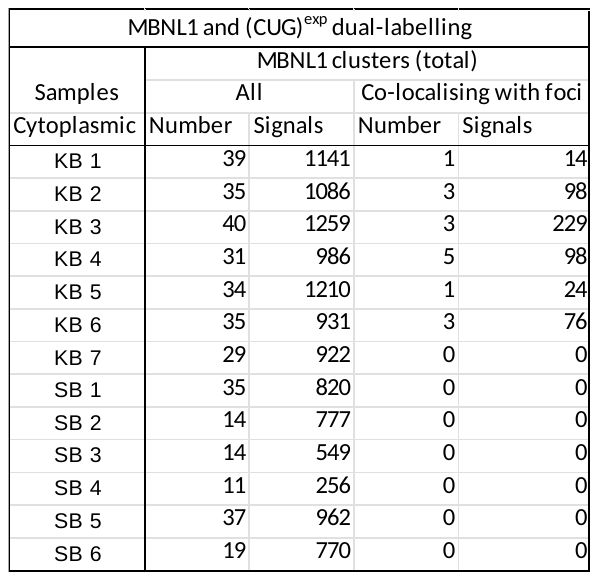

B

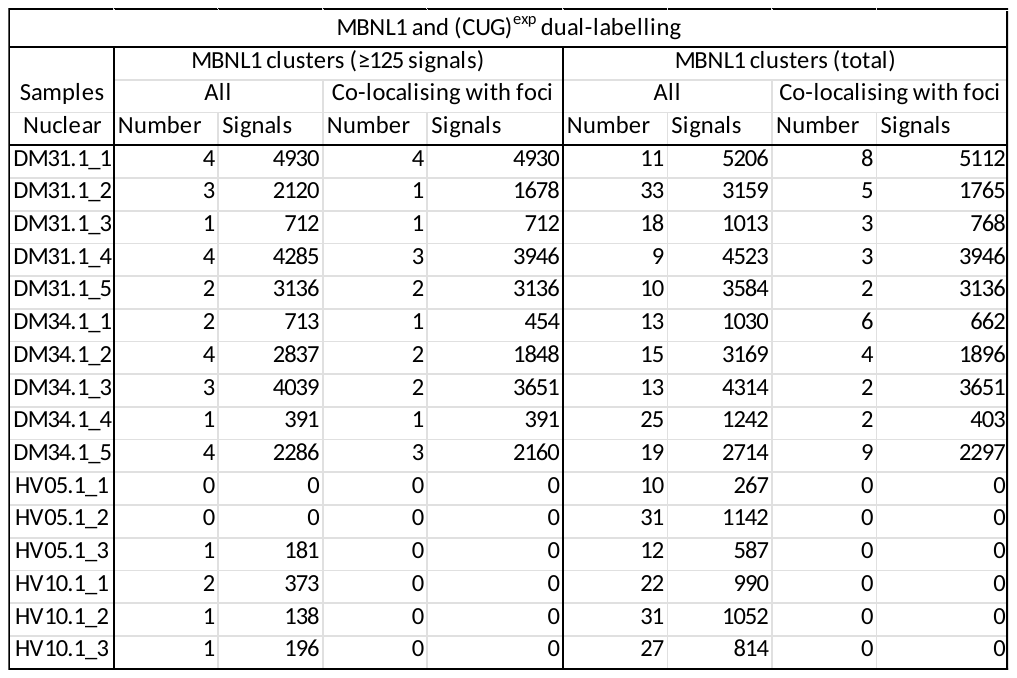

C

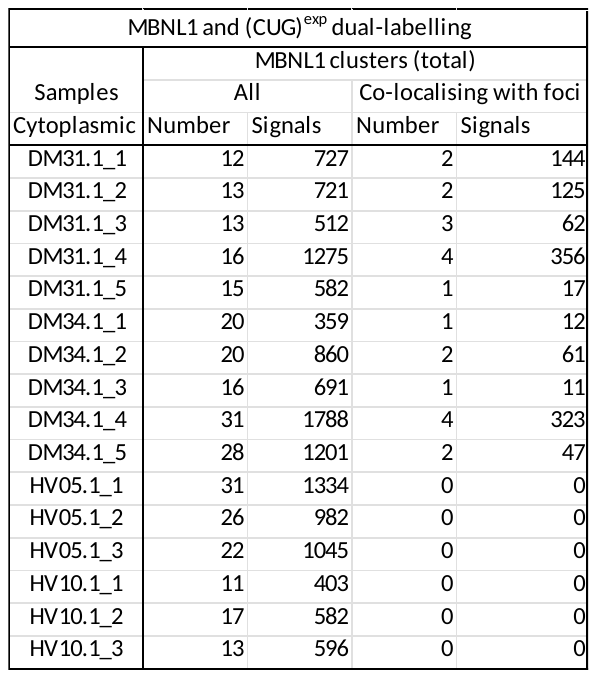

D

The “Number” columns show the counts of MBNL1 clusters in the nucleus of cell samples (Table A) and muscle biopsies (Table C), as well as in the cytoplasm of cell samples (Table B) and muscle biopsies (Table D). Additionally, these columns provide counts of MBNL1 clusters that exhibit co-localisation with (CUG)^exp^ foci. The “Signals” columns present the corresponding total count of probe signals detected within these clusters.

**Poly(A) and 5’ cap status of mutant *DMPK* mRNA**

Hyperadenylation of RNA is known to be associated with nuclear retention, but the cause-consequence relationship between hyperadenylation and regulation of RNA nuclear export is still unclear [5-7]. Additionally, absence of the 7-methylguanosine (m^7^G) cap at the 5’ end of pre-mRNA could cause the 5’->3’ degradation of mRNA by XRN exonucleases [8-10]. To compare both the 3’ polyadenylation and 5’ cap structure status of *DMPK* transcripts in the non-DM cells (SB, control) and DM1 patient cells (KB) and their effect on presence and stabilisation of mutant *DMPK* transcripts, we subjected RNA from KB and SB to poly(A) selection using oligo(dT) beads and 5’ cap selection using 5'-Phosphate-Dependent Exonuclease. 5'-Phosphate-Dependent Exonuclease, a processive 5'->3' exonuclease digests RNA having a 5'-monophosphate. The enzyme does not digest RNA having a 5'-triphosphate, a 5'-cap or a 5'-hydroxyl group. Thus, it can be used to isolate mature mRNA with an m^7^G-cap.

In SB, the poly(A) selection and 5’ cap selection recovered 89.0% and 99.0% of the G-allele *DMPK* transcripts, respectively, compared to untreated G-allele *DMPK* transcripts (**Fig. S10**). In contrast, in KB, only 62.9% and 68.7 % of the mutant (G-allele) *DMPK* transcripts were recovered from the Poly(A) selection and 5’ cap selection, respectively, compared to untreated mutant *DMPK* transcripts (**Fig. S10**). One possible explanation for this observation would be that a significant portion of the mutant *DMPK* mRNA was subject to deadenylation or decapping. Our previous results indicate that the mutant *DMPK* mRNA is primarily retained in the nucleus, which could lead to its recognition as abnormal or faulty RNA, triggering RNA decay (**Fig. 1 G**) [11, 12]. However, due to the complex hairpin structures in the CUG repeats and sequestration of MBNL [13-15], degradation may be initiated but impeded to a large extent. Another explanation might be that a considerable portion of the long CUG expanded *DMPK* mRNA fractured during experimental handling and only part of a transcript harbouring a poly(A)-tail or a m^7^G-cap was captured by the beads and RNA bind columns. Moreover, influence of the expanded CUG repeats on oligo(dT) binding avidity, caused by topological constraints in the expanded transcripts, could not be excluded at this point.

**Fig. S10 Assessment of poly(A) tail and 5’ cap structure status of the mutant (CUG)^exp^ *DMPK* transcripts.**

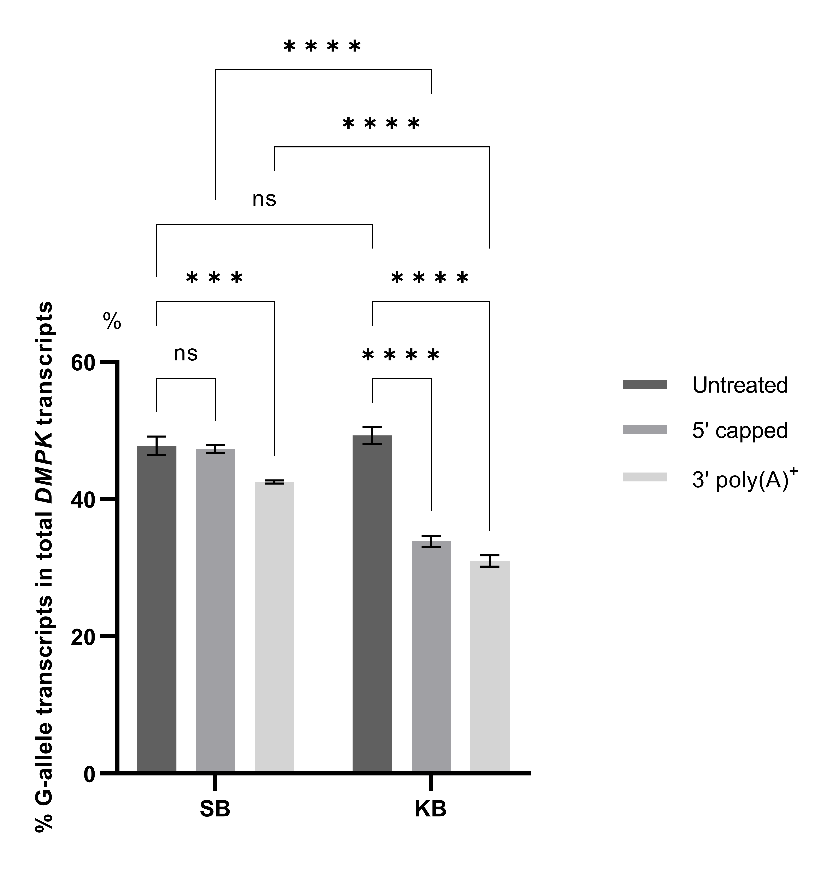

dPCR quantification of mutant (CUG)^exp^ *DMPK* transcripts as a percentage of total *DMPK* RNA, using allele-specific probes following total RNA extraction from KB, SB in conjunction with poly(A) selection or 5’ cap selection. The G-allele *DMPK* transcript represents the mutant (CUG)^exp^ *DMPK* transcript in KB. The experiment was performed in triplicate to ensure the reproducibility of the results. Statistical significance was determined by two-way ANOVA (mean ± SD, *** for P<0.001, **** for P<0.0001) or indicated as not significant (NS) for P>0.05.

Fig. S11 Assessment of the knock-down efficiency of RNA decay factors by Western blotting.

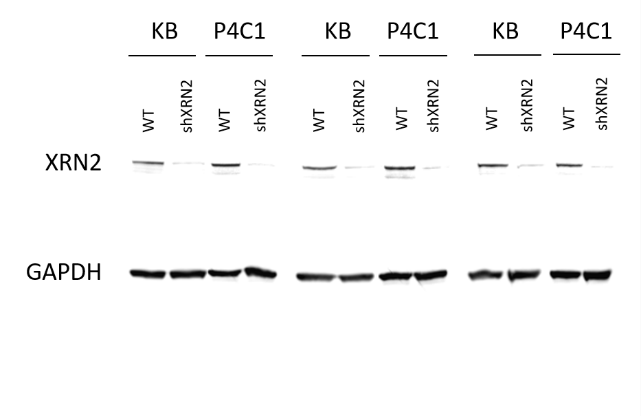

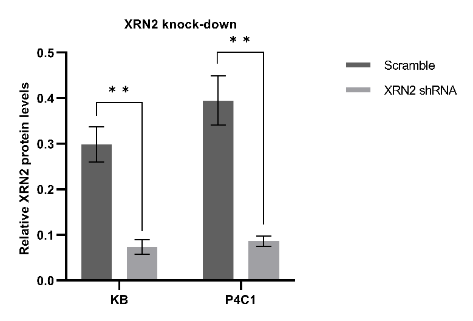

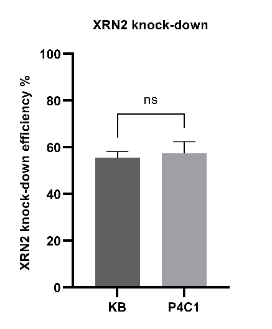

**C**

**B**

**A**

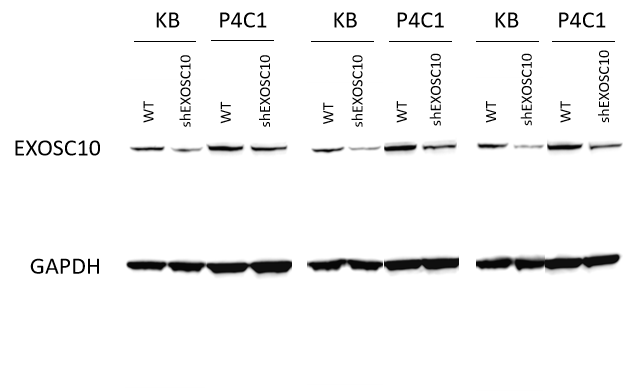

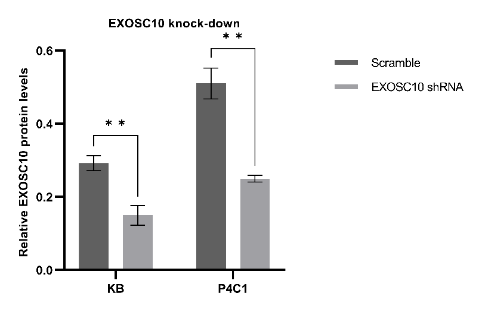

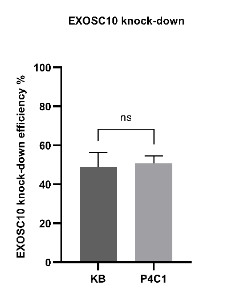

**F**

**E**

**D**

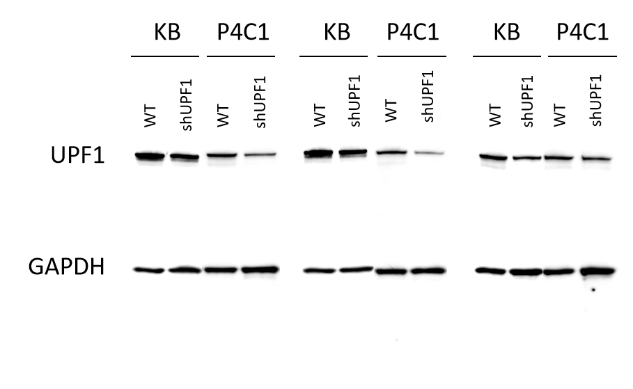

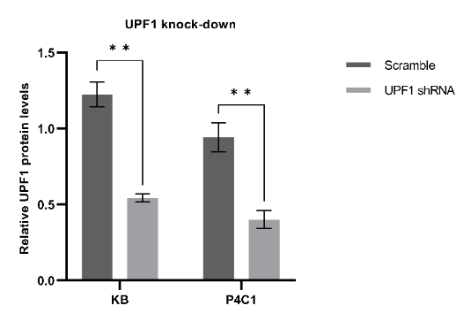

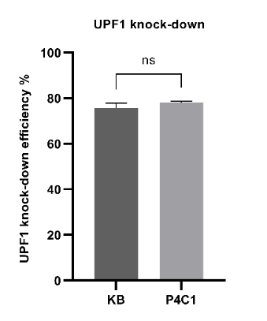

**I**

**H**

**G**

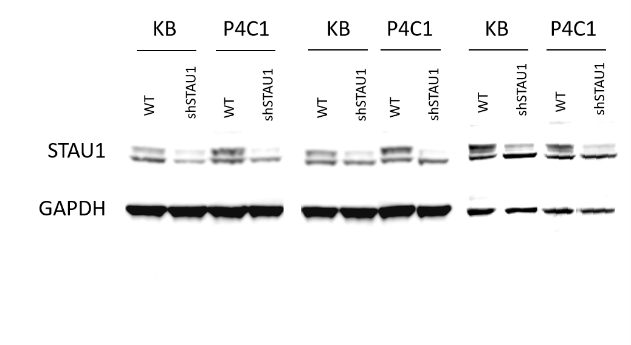

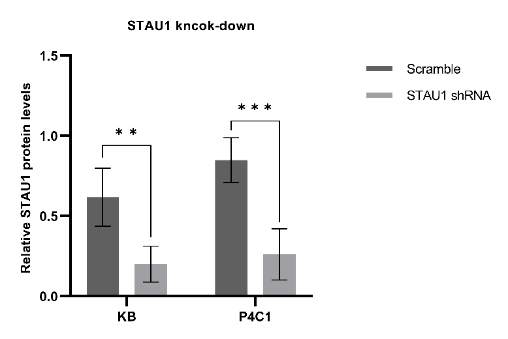

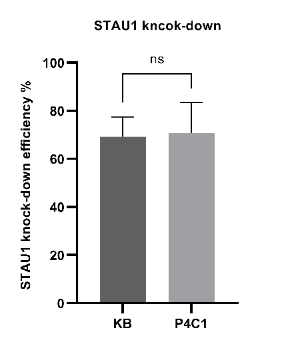

**L**

**K**

**J**

Western blot analyses showing the protein levels in KB (DM1) and P4C1 (MBNL-deficient DM1) cells following the lentiviral transfection with scramble control and shRNAs target *XRN2* (A), *EXOSC10* (D), *UPF1* (G) and *STAU1* (J). The experiment was performed in triplicate to ensure the reproducibility of the results. (B, E, H, K) Quantification of the relative protein levels of XRN2 (B), EXOSC10 (E), UPF1 (H) and STAU1 (K) following the lentiviral shRNA-mediated knock-down in KB and P4C1 cells, normalised against GAPDH values. (C, F, I, L) Comparisons of the knock-down efficiency of XRN2 (C), EXOSC10 (F), UPF1 (I) and STAU1 (L) between KB and P4C1. Statistical significance was determined by two-way ANOVA or two-tailed student t-test (mean ± SD, ** for P<0.01, *** P<0.001) or indicated as not significant (NS) for P>0.05.

Fig. S12 Assessment of the efficiency of *UPF1* over-expression by Western blotting.

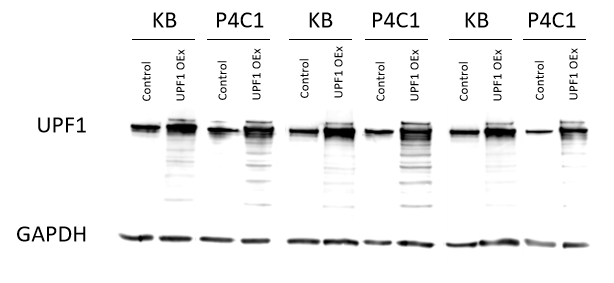

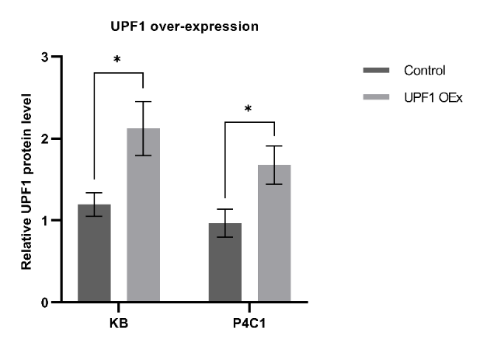

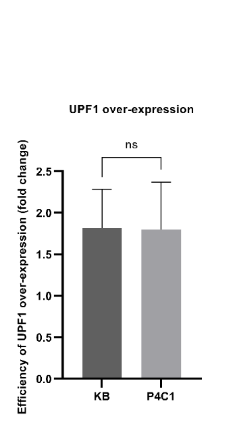

**C**

**B**

**A**

(A) Western blot analyses showing the UPF1 protein levels in KB and P4C1 following the lentiviral transfection with empty control or *UPF1* cDNA. The experiment was performed in triplicate to ensure the reproducibility of the results. (B) Quantification of the relative UPF1 protein levels, normalised against GAPDH values. (C) Comparison of the *UPF1* over-expression efficiency between KB and P4C1. Statistical significance was determined by two-way ANOVA or two-tailed student t-test (mean ± SD, * for P<0.05) or indicated as not significant (NS) for P>0.05.

Table S9 Total number of cells examined by FISH.

|  | KB | | | | | | P4C1 | | | | | |
| --- | --- | --- | --- | --- | --- | --- | --- | --- | --- | --- | --- | --- |
|  | Scramble/  control | | | shRNA/  expression | | | Scramble/  control | | | shRNA/  expression | | |
|  | 1^st^ | 2^nd^ | 3^rd^ | 1^st^ | 2^nd^ | 3^rd^ | 1^st^ | 2^nd^ | 3^rd^ | 1^st^ | 2^nd^ | 3^rd^ |
| XRN2 KD | 113 | 92 | 144 | 99 | 85 | 77 | 98 | 69 | 69 | 18 | 25 | 22 |
| EXOSC10 KD | 37 | 45 | 46 | 35 | 28 | 32 | 104 | 88 | 69 | 34 | 30 | 28 |
| UPF1 KD | 78 | 94 | 107 | 110 | 81 | 65 | 45 | 38 | 33 | 24 | 16 | 27 |
| STAU1 KD | 32 | 41 | 32 | 26 | 27 | 33 | 121 | 72 | 66 | 21 | 17 | 25 |
| UPF1 OEx | 226 | 180 | 182 | 80 | 109 | 88 | 174 | 209 | 145 | 30 | 33 | 28 |

Fig. S13 Probes positioning in the *DMPK* cDNA.

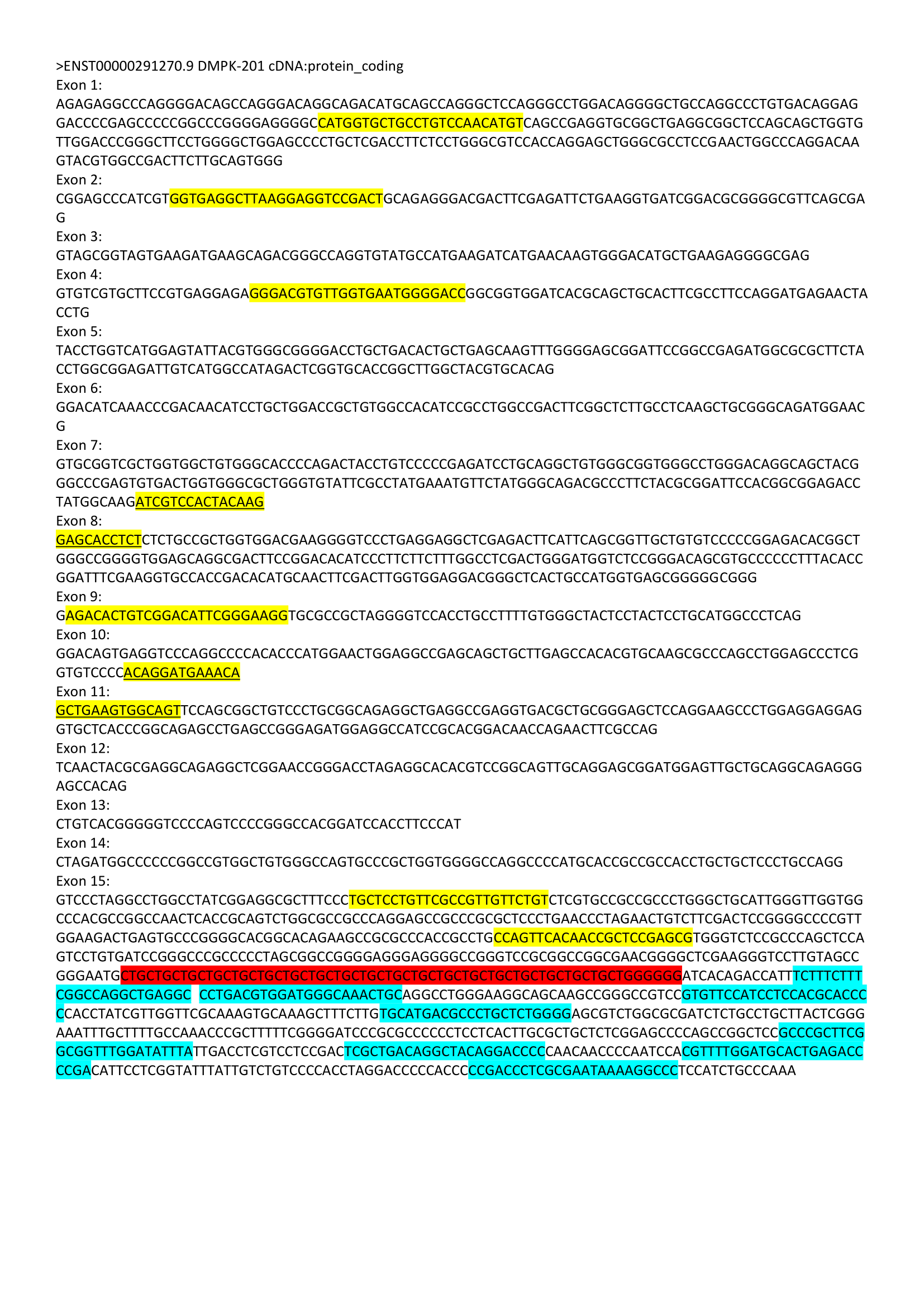

The figure illustrates the positions of 5' and 3' single-copy probes employed in STORM experiments. The *DMPK* cDNA is depicted, with the 5' single-copy probes highlighted in yellow and the 3' single-copy probes highlighted in blue (Probes spanning two exons are also underlined). The CTG repeat region is specifically indicated in red.
